# Genomic consequences of domestication of the Siamese fighting fish

**DOI:** 10.1101/2021.04.29.442030

**Authors:** Young Mi Kwon, Nathan Vranken, Carla Hoge, Madison R Lichak, Kerel X Francis, Julia Camacho-Garcia, Iliana Bista, Jonathan Wood, Shane McCarthy, William Chow, Heok Hui Tan, Kerstin Howe, Sepalika Bandara, Johannes von Lintig, Lukas Rüber, Richard Durbin, Hannes Svardal, Andres Bendesky

**Author notes:** equal contribution (Hannes Svardal, Andres Bendesky). Corresponding authors. (AB), (HS), (RD).

## Abstract

Siamese fighting fish, commonly known as betta, are among the world’s most popular and morphologically diverse pet fish, but the genetic processes leading to their domestication and phenotypic diversification are largely unknown. We assembled de novo the genome of a wild *Betta splendens* and whole-genome sequenced multiple individuals across five species within the *B. splendens* species complex, including wild populations and domesticated ornamental betta. Given our estimate of the mutation rate from pedigrees, our analyses suggest that betta were domesticated at least 1,000 years ago, centuries earlier than previously thought. Ornamental betta individuals have variable contributions from other *Betta* species and have also introgressed into wild populations of those species. We identify *dmrt1* as the main sex determination gene in ornamental betta but not in wild *B. splendens*, and find evidence for recent directional selection at the X-allele of the locus. Furthermore, we find genes with signatures of recent, strong selection that have large effects on color in specific parts of the body, or the shape of individual fins, and are almost all unlinked. Our results demonstrate how simple genetic architectures paired with anatomical modularity can lead to vast phenotypic diversity generated during animal domestication, and set the stage for using betta as a modern system for evolutionary genetics.

**One-Sentence Summary:** Genomic analyses reveal betta fish were domesticated more than 1,000 years ago and the genes that changed in the process.

## Main Text

Domesticated animals have provided important insights into the genetic bases of a wide range of morphological, physiological, and behavioral traits. Because of their intimate relationship with people, domesticates have also furthered our understanding of human history and culture, and of our interactions with other species (*1*). Genetic studies of animal domestication, however, have largely focused on mammals and birds (*1*, *2*), and only few genome-wide analyses of fish domestication have been performed (*3*–*5*).

Siamese fighting fish have been selectively bred for fighting in Southeast Asia for centuries, with reports dating back to as early as the 14th century A.D. in Thailand, making them one of the oldest fish domestications (*6*). Starting in the early 20th century, Siamese fighting fish also began to be bred for ornamental purposes, becoming one of the world’s most popular pet fish, commonly known as betta (*7*). Although it is generally presumed —based on morphology and few genetic markers (*8*, *9*)— that domesticated fighting fish derive mainly from *Betta splendens*, it has been suggested that other closely related species (collectively called the *Betta splendens* species complex) may have contributed to modern varieties (*10*). Ornamental betta have been diversified from their short finned ancestors into an astonishing array of fin morphologies, colors and pigmentation patterns, providing a rich phenotypic repertoire for genetic analysis. This remarkable and long history of domestication for fighting, followed by breeding for ornamental purposes, combined with one of the smallest vertebrate genomes at only ~450 megabase pairs (Mbp) (*11*–*13*), makes betta an appealing subject for evolutionary genetic studies of domestication.

Here, we use a synergistic combination of population and quantitative genetic approaches to investigate the historical processes and molecular changes that lead to the domestication and phenotypic diversification of betta fish.

### A wild *Betta splendens* reference genome

We generated a high-quality reference genome assembly of wild *B. splendens* using long-read PacBio technology, optical mapping with BioNano, scaffolding with 10X Genomics linked reads, and polishing with Illumina short reads. We obtained a genome reference comprised of 441 Mb, of which 98.6% is assigned to the 21 chromosomes expected from its karyotype (*14*), with a contig N50 of 2.50 Mb and scaffold N50 of 20.13 Mb, meeting the standards set forth by the Vertebrate Genomes Project (*15*). To annotate the genome, we performed RNA sequencing from male and female brain, fin, liver, spleen, and gonad. This annotated reference genome is now the representative *B. splendens* reference in NCBI (fBetSpl5.3, GCA_900634795.3).

To discover structural chromosomal rearrangements that may have arisen during domestication, we performed whole genome alignments using three ornamental betta references (*11*–*13*) and our wild *B. splendens* reference, with *Anabas testudineus* (climbing perch) as an outgroup (*8*, *15*). Except for a large intrachromosomal rearrangement of chromosome 16 in ornamental betta, the genome was largely syntenic between wild *B. splendens* and ornamental betta (Suppl. Fig. 1, Note 1).

### Complex evolutionary relationships between *Betta* species

To determine the genetic origin of ornamental betta and understand its relationships with species of the *Betta splendens* complex, we sequenced to ~15× coverage the whole genomes of (i) 37 ornamental betta from different sources, representing a diversity of ornamental traits (Fig. 1A,B; Suppl. Table 1); (ii) 58 wild individuals, including representatives of all species of the *B. splendens* complex (except for *Betta stiktos*), and four populations of *B. splendens* from different parts of its natural range (Fig. 1A); and (iii) an outgroup (*Betta compuncta*). We aligned the sequencing reads to our *B. splendens* reference genome, then called and filtered variants to generate a final set of 27.8 million phased biallelic SNPs.

**Figure 1.**
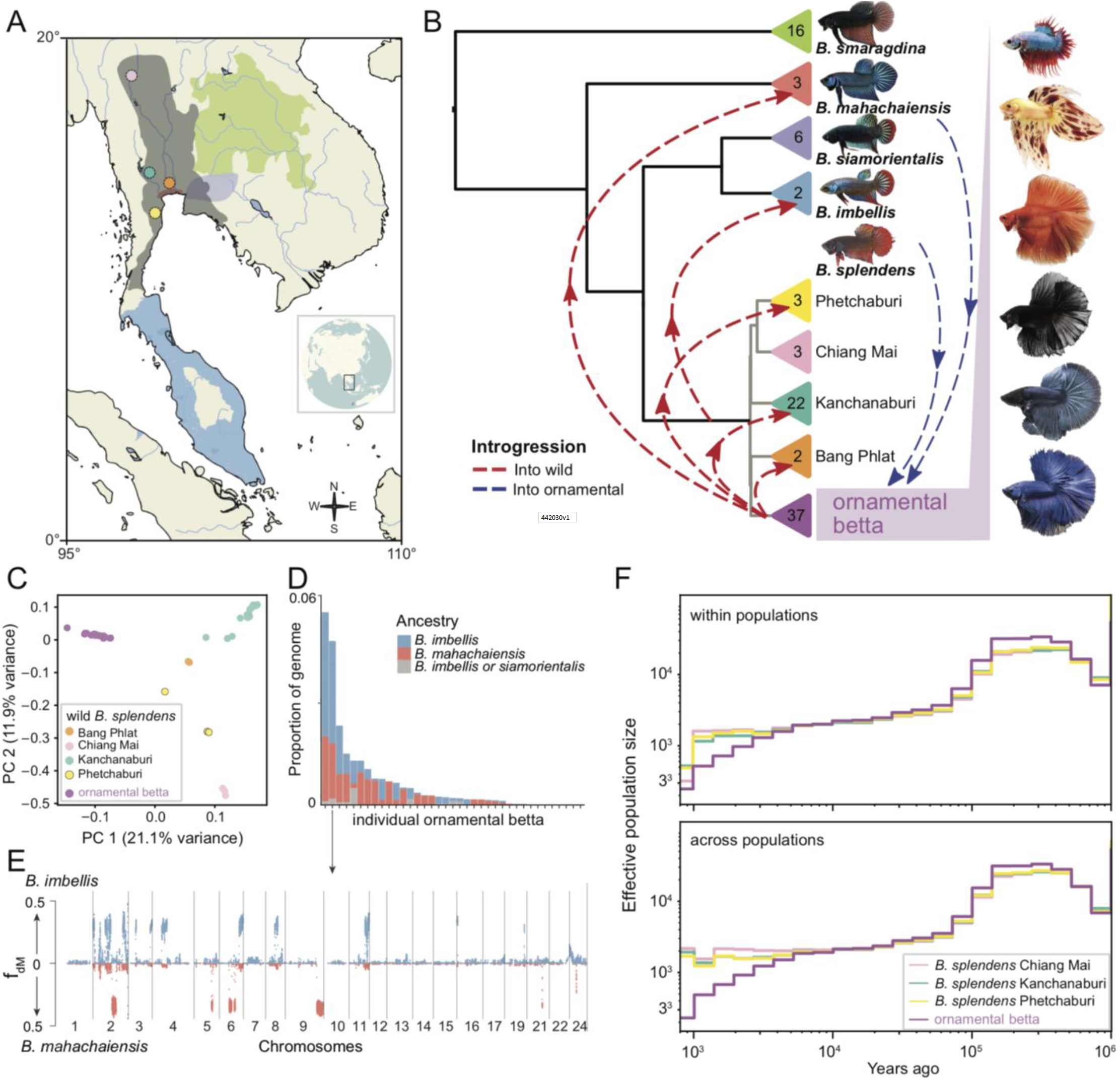
*Betta* phylogeny, gene flow, and demographic history. **(A)** Distribution ranges of *B. splendens* species complex and sampling locations. Colors according to panel B. **(B)** Species and population relatedness based on neighbor-joining of group pairwise genetic differences. Arrows denote introgression events involving specific samples. Triangles contain sample numbers. Photos show representative males of ornamental varieties; from top to bottom: crowntail plakat, dalmatian veiltail, orange doubletail, superblack halfmoon, steel halfmoon, royal blue halfmoon. Image credits, with permission: Frank Sriborirum and Kasey Clark. **(C)** Principal component analysis of *B. splendens* samples. Further principal components are in Suppl. Fig. 7. **(D)** Proportion of genome introgressed from non-*splendens* species in each ornamental individual based on f_dM_ and regional trees. **(E)** Genome-wide fdM plot of ornamental betta Orn45 (p1=other ornamental bettas; p2=Orn45; p3=*B. mahachaiensis* or *imbellis*; outgroup=*B. compuncta*). **(F)** Effective population size as estimated by Relate within populations (top panel) and between sequences from wild *B. splendens* populations and ornamental betta (bottom panel); see Methods.

We first assessed relationships across the wild species of the *B. splendens* complex by constructing neighbor-joining (NJ) and maximum-likelihood (ML) based phylogenies (Fig. 1B; Suppl. Fig. 2A,B). We observed strong bootstrap support for *B. smaragdina* as the outgroup to the other species of the *B. splendens* complex with *B. mahachaiensis* as the outgroup to the remaining species. *B. imbellis* and *B. siamorientalis* together form a sister clade to all wild *B. splendens* populations.

We then tested for evidence of evolutionary processes that violate tree-like species relationships such as hybridization, by computing ABBA-BABA statistics (Patterson’s *D* and f4 admixture ratio (*16*)) for all triplets of individuals organized according to the phylogeny. This analysis revealed widespread patterns of excess allele sharing between non-sister species, suggesting that the speciation history of these groups was complex, involving either structured ancestral populations, cross-species gene flow, or both (Fig. 1B; Suppl. Fig. 3A; Suppl. Note 2). Interestingly, two out of three *B. mahachaiensis* samples and one of the two *B. imbellis* samples showed highly significant excess allele sharing with *B. splendens* populations compared to their conspecifics sampled from different locations, consistent with gene flow from *B. splendens* into particular populations of these species (Suppl. Fig. 3A,B; Suppl. Note 2).

### Ornamental betta derive from *B. splendens* but have variable contributions from other species

Adding the ornamental betta samples to the phylogeny, we found that they cluster with *B. splendens* (Fig. 1B; Suppl. Fig. 2). This result was also observed through principal component analysis (PCA), where ornamental betta showed no apparent loading on axes representing non-*splendens* species (Suppl. Fig. 7). In both phylogenies and PCA, ornamentals form a clearly defined group distinct from all wild *B. splendens* populations (Fig. 1B,C). These results indicate that ornamental betta are genetically most similar to *B. splendens.*

To test whether ornamental betta carry non-*splendens* ancestry, we computed all ABBA-BABA tests of the form *D*(ornamental except focal, focal ornamental; non-*splendens* species, outgroup) (Suppl. Fig. 4G-J). These tests revealed that 76% (28 out of 37) of ornamental betta carry significant ancestry from non-*splendens* species. To examine the chromosomal distribution of non-*splendens* ancestry in these individuals, we computed regional ABBA-BABA statistics (f_dM_) along their genomes and confirmed non-*splendens* ancestry in high-f_dM_ regions by constructing local gene trees (Fig. 1D,E; Suppl. Figs. 8,12). The analyses revealed that signals of excess allele sharing are driven by genomic tracts where one or, more rarely, both haplotypes of the focal sample clustered with *B. imbellis* or *B. mahachaiensis* (Supp. Fig. 8). The genomic locations of these tracts, which encompass between 0 and 6% of the genomes of ornamental betta (Fig. 1D), are generally different among individuals (Suppl. Fig.12). The ornamental sample with the second highest levels of introgression from other species is particularly interesting, since some of its chromosomes are a mosaic of alternating regions of *B. imbellis* and *B. mahachaiensis* ancestry, consistent with a natural or man-made hybrid of those species having been backcrossed into ornamental betta (Fig. 1E). Altogether, our analyses indicate that ornamental betta are clearly derived from *B. splendens*, yet most individuals have relatively recent contributions from *B. mahachaiensis* and *B. imbellis*.

### Ornamental betta introgression is widespread among wild *Betta*

Interestingly, the topology of relationships between wild *B. splendens* populations in NJ-based phylogenies changed after including ornamental bettas (Suppl. Fig. 2A,C; Suppl. Note 3,4). To further investigate this, we computed ABBA-BABA statistics within the framework of the phylogeny including ornamentals (Suppl. Fig. 2C), and assessed each individual’s relationship with respect to the other species of the *Betta splendens* species complex, as well as ornamentals (Suppl. Fig. 5A,7). Together, these analyses revealed strong evidence for ornamental betta ancestry in two out of three wild *B. mahachaiensis* samples and in individuals from three out of four populations of wild *B. splendens* (Suppl. Note 4). Investigating the signals along the genome, we found that for the two *B. mahachaiensis* samples, *mahachaiensis*-like and ornamental-like haplotypes alternate at near-chromosome scale, suggesting an ornamental ancestor only a few generations back (Suppl. Fig. 6B). Conversely, for wild *B. splendens* individuals with ornamental betta ancestry, the genome-wide signals of excess allele sharing with ornamentals were diffusely distributed along the chromosomes with only a few relatively short, clearly distinguishable ornamental haplotypes (Suppl. Fig. 6A), suggesting that there was enough time for introgressed haplotypes to be broken down by recombination. In summary, ornamental introgression into wild *Betta* seems to be geographically diffuse and to have happened both long ago and very recently. This finding is perhaps related to the practice by breeders of releasing excess domesticated betta into the wild and may constitute a conservation threat to wild *Betta* populations.

### Timing the domestication of *B. splendens*

To determine when ornamental betta initially diverged from wild populations, we performed coalescence-based demographic analysis. In order to date events in the domestication of *B. splendens*, we needed to know the germline mutation rate. To determine this, we sequenced an ornamental trio and a quartet to >30× coverage and found the mutation rate to be 3.75×10^−9^ per bp per generation (95% CI: 9.05×10^−10^ to 9.39×10^−9^). This rate is similar to the rate previously inferred for cichlids (*17*) and approximately 3-fold lower than that of humans (*18*). Assuming a generation time of six months (*7*), our demographic analyses suggest that ornamental and wild populations began to split around 4,000 years ago (~1,000 to ~7,000 years based on mutation rate CI). This divergence was coupled to a reduction in population size in ornamental betta as would be expected if a subset of wild individuals began to be bred in captivity (Fig. 1F; Suppl. Fig. 9). Low nucleotide diversity (0.00137 per bp in wild fish and 0.00113 per bp in ornamental betta) and elevated linkage disequilibrium relative to the wild populations further support a decrease in population size throughout domestication that has not fully recovered (Suppl. Figs. 10,12). Even the lower bound (~1,000 years ago) for divergence of the ornamental betta population from wild is earlier than the origin of domestication in the 14th century previously suggested by historical documents (*6*).

### Genetic signals of selection in ornamental betta

Genetic variants that increase fitness in captivity or that are associated with phenotypic traits actively selected by breeders are expected to increase in frequency during domestication. To discover such loci with signatures of selective sweeps in ornamental betta, we searched for extended homozygosity tracts using H-scan (*19*) and for high-frequency haplotypes using G12 (*20*) across 37 ornamental betta (Fig. 2A). Both tests identified concordant loci with strong evidence of selective sweeps in 11 of the 21 *B. splendens* chromosomes, and peaks remained when run on a downsampled set of 24 ornamentals (Supp. Fig 11). Equivalent selection scans using whole-genome sequencing of 24 wild *B. splendens* did not reveal clear signals (Fig. 2A). These results are consistent with footprints of selection in ornamental betta being related to the domestication process.

**Figure 2.**
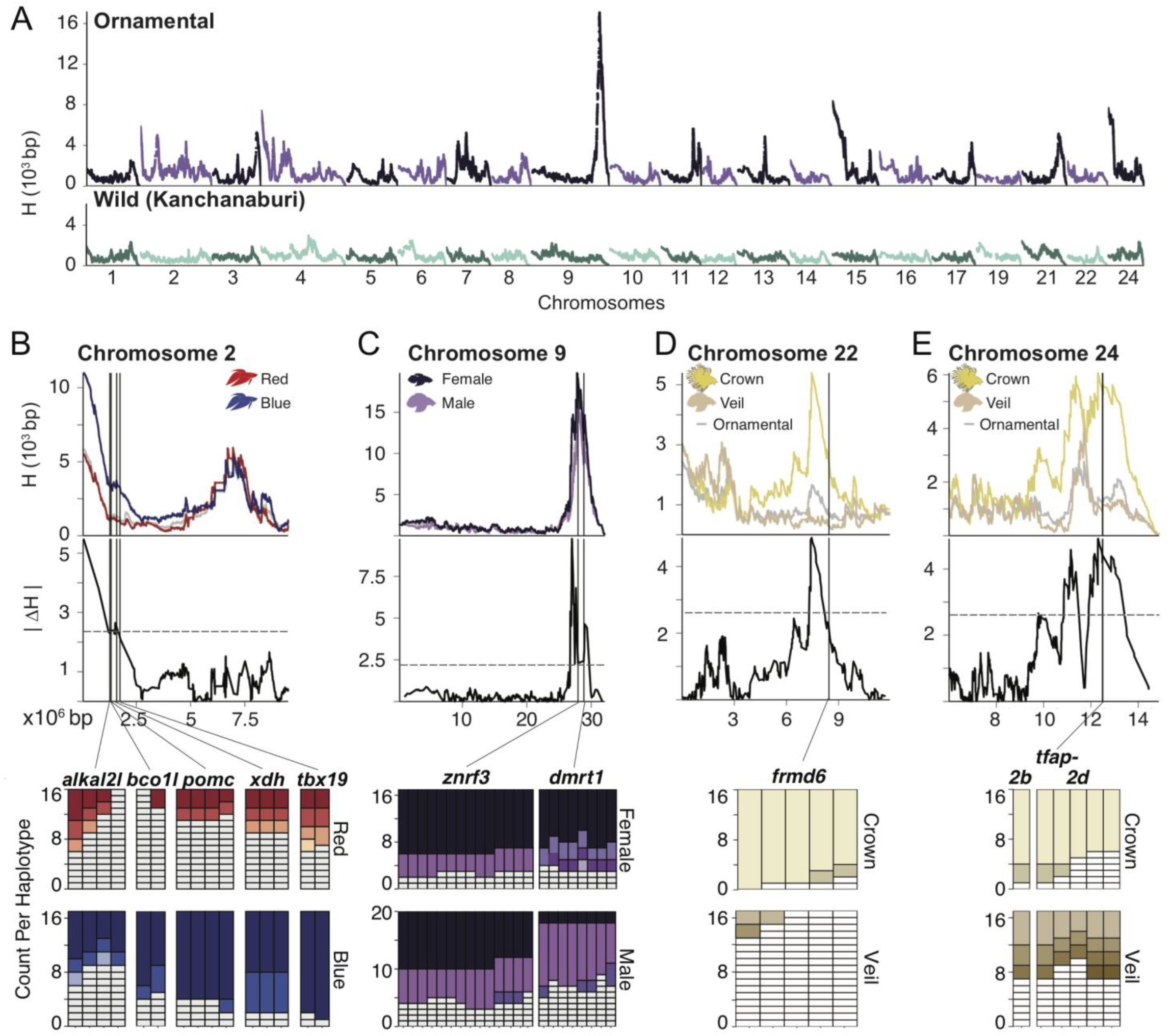
Genomic signals of selection in ornamental betta. **(A)** Genome-wide H-scan within ornamental betta (n=37) and within wild *B. splendens* from Kanchanaburi (n=24). **(B-E)** Top: H-scan close-up of ornamentals (n=37, grey line). Middle: ornamentals separated by **(B)** color (red: 17, blue: 17), **(C)** sex (female: 17, male: 20), and **(D,E)** fin morphology (crown: 16, veil: 18). Grey dashed lines denote genome-wide threshold of significance (α=0.05) for the absolute difference in H-scan (|ΔH|) between ornamentals separated by color, sex, and fin type. **(B-E)**Bottom: Distribution of haplotypes in genes identified as outliers in both H-scan and G12 and statistically different between ornamentals separated by color, sex, and fin morphology which are further discussed in Figs. 3-5. White indicates a haplotype observed only in a single fish.

The most prominent selection peak shared across ornamentals but absent in wild *B. splendens* falls on chromosome 9 and is centered on *zinc and ring finger 3* (*znrf3*). In zebrafish, *znrf3* is required for the formation of fin rays, and in mammals it is required for limb formation and testis development (*21*–*23*). All the ornamental fish we sequenced have large fins compared to wild *B. splendens*. Therefore, we hypothesize that *znrf3* has contributed to either sexual development or the expansion of fins during betta domestication.

The majority (34/37) of the ornamental betta we sequenced represented four of the most popular varieties along two phenotypic dimensions: color and fin morphology. The fish were royal blue (n=17), solid red (n=17), veiltail (n=18) and crowntail (n=16), represented by males (n=20) and females (n=17). Veiltails are characterized by large, flowing caudal fins, and crowntails have fins that are webbed between the rays (Fig. 1B). To determine whether the footprints of selection we detected were driven by fish of a particular variety or sex, we compared H-scan and haplotype frequencies across subsets of fish representing these traits (Fig. 2B-E).

A peak close to *znrf3*, centered on *double-sex and mab-3 related transcription factor 1* (*dmrt1*), became apparent when comparing males to females (Fig. 2C). *dmrt1* is critical for gonad development in vertebrates, and functions as the sex determination gene in several fish species (*24*–*26*), in *Xenopus laevis* frogs (*27*), and in birds (*28*), suggesting *dmrt1* has a role in sex determination in betta.

A strong sweep in blue fish on chromosome 2 harbors multiple genes involved in pigmentation (Fig. 2B): *proopiomelanocortin* (*pomc*), which encodes alpha and beta melanocyte stimulating hormones) (*29*); *T-box transcription factor 19* (*tbx19*), which encodes a transcription factor expressed specifically in pituitary cells that will express *pomc (30)*; *xanthine dehydrogenase* (*xdh*), which encodes an enzyme whose homologs synthesize yellow-red pteridine pigments (*31*, *32*); *ALK and LTK-ligand 2-like* (*alkal2l*), which encodes a cell-signaling molecule important for the development of iridophores (*33*–*36*), and *beta-carotene oxygenase like-1* (*bco1l*), which encodes an enzyme whose homologs metabolize orange-red carotenoid pigments (*37*, *38*). These results suggest that one or more of these pigmentation genes were a target of selection by betta breeders.

Two selection peaks, one on chromosome 22 and another on chromosome 24, were not detected when all ornamental fish were combined or in an analysis including only veiltail fish, but were significant in the subset of crowntail fish (Fig. 2D,E), suggesting their importance to crowntail fin morphology.

### The evolution of sex determination

To test whether the loci containing *znrf3* and *dmrt1*, which had evidence of a selective sweep in ornamental betta, are involved in sex determination, we performed a genomewide association study (GWAS) using sex as the phenotype. We focused on ornamental betta, since we had a large enough sample size (20 males and 17 females) to detect variants with large effect on sex. A ~30-kb region overlapping *dmrt1* but not *znrf3* was strongly associated with sex, with 16/17 females being homozygous at the most strongly associated SNPs, while 16/20 males were heterozygous (Fig. 3A,C). We call “Y” the male-specific allele of *dmrt1* and “X” the allele present in both males and females. These results strongly implicate *dmrt1* as the sex determination gene in ornamental betta and indicate that males are the heterogametic sex.

**Figure 3.**
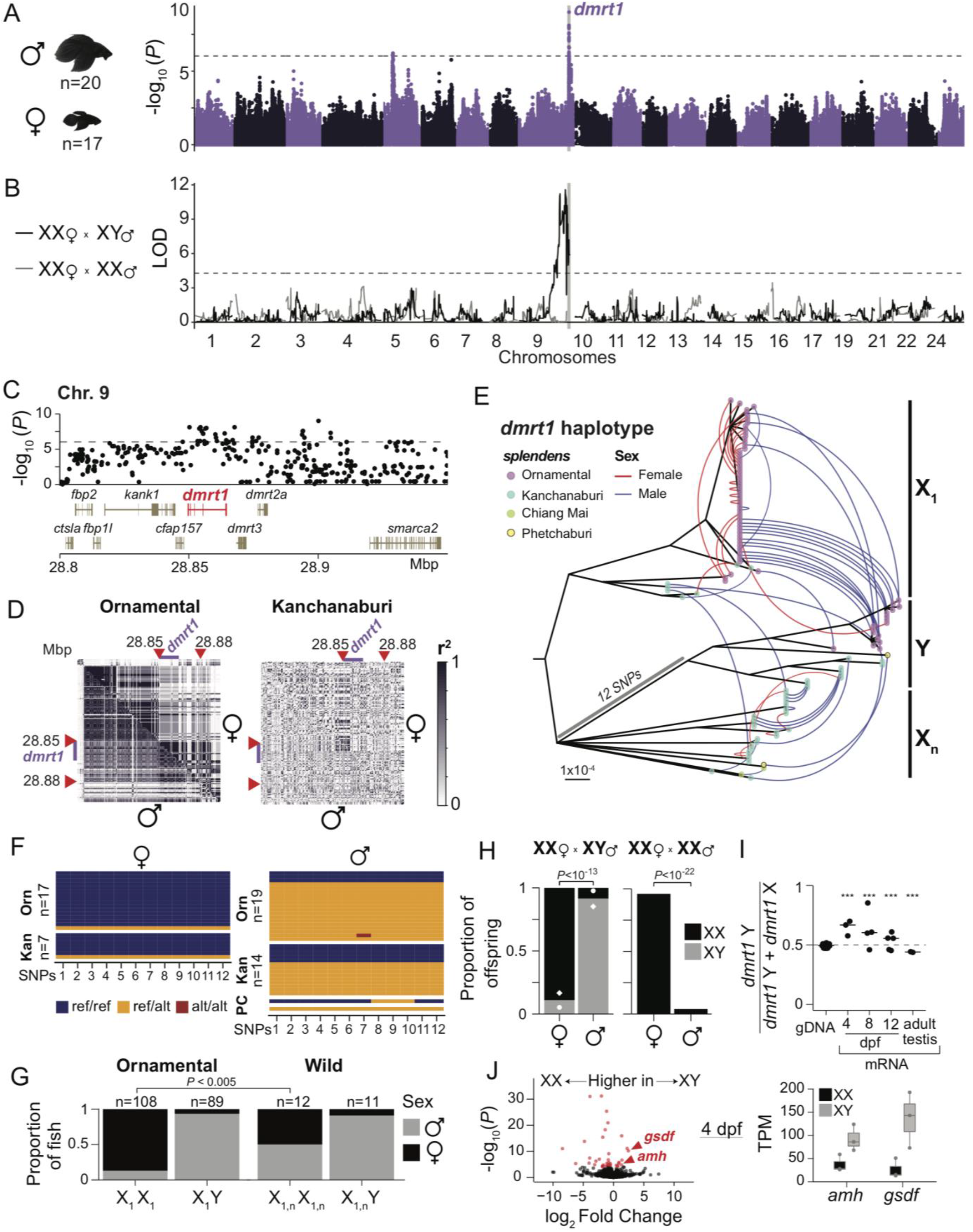
*dmrt1* is a sex determination gene in ornamental betta. **(A)** Manhattan plot for GWAS of sex in ornamental betta. Dashed line denotes the genome-wide significance threshold; see Methods. **(B)** QTL mapping of two F2 intercrosses: *dmrt1*_X_1_X_1_ female × *dmrt1*_X_1_Y male cross with 211 F2 hybrids (black line); *dmrt1*_X_1_X_1_ female × *dmrt1*_X_1_ X_1_ male cross with 100 F2 hybrids (grey line). Dashed line denotes the genome-wide significance threshold at α=0.05. **(C)** GWAS zoom-in of the *dmrt1* locus with gene annotations. **(D)** Linkage disequilibrium plot for chromosome 9 surrounding the *dmrt1* locus for ornamental betta and wild *B. splendens* from Kanchanaburi. Upper triangle, females; lower triangle, males. Red arrowheads denote the region of the *dmrt1* haplotype in (E). Purple bar denotes *dmrt1*. **(E)** Maximum likelihood phylogeny with at least 80% bootstrap support for the *dmrt1* haplotype across *B. splendens* rooted by *B. siamorientalis.* Each tip represents one of the two alleles of a sample, colored by population. Arches link the alleles of each sample and are colored by gonadal sex. **(F)** Genotypes of males and females from ornamental betta and wild *B. splendens* across 12 SNPs present in all samples within the *dmrt1_*Y group. P=Phetchaburi; C=Chiang Mai. **(G)** Sex ratios for *dmrt1* haplotypes observed in ornamental betta and wild *B. splendens*. *P* denotes the result of Fisher’s exact test. **(H)** Average haplotype ratios across sex for offspring from two *dmrt1*_X_1_X_1_ female × dmrt1_X_1_Y male crosses (n=101 labeled with diamond,112 labeled with circle) and *dmrt1*_X_1_X_1_ male × *dmrt1*_X_1_X_1_ female cross (n=100). ***, *P*<0.001 by Fisher’s exact test. **(I)** Allele-specific expression of *dmrt1* across days post fertilization (dpf) for *dmrt1*_X_1_Y larvae from a *dmrt1*_X_1_X_1_ × *dmrt1*_X_1_Y cross. Each dot represents a sample. ***, *P*<0.001 by binomial test. **(J)** Differential mRNA expression between 3 *dmrt1*_X_1_Y and 3 *dmrt1*_X_1_X_1_ 4-dpf larvae. Red denotes expression differences where *P*<10^−6^. Left: Differential mRNA expression between 3 *dmrt1*_X_1_Y and 3 *dmrt1*_X_1_X_1_ 4-dpf larvae. Red denotes expression differences where *P*<10^−6^. Right: Transcripts per million (TPM) of two genes important for male sex development: *anti-mullerian hormone* (*amh*) and *gonadal soma-derived factor* (*gsdf*).

The reference genome was generated from a wild male *B. splendens*, so genomic sequences present only in females or only in ornamental betta would not be represented in the SNPs that we used for GWAS. Only 0.5% of sequencing reads of individuals from both sexes could not be mapped to the reference genome (male vs female *P*=0.64), indicating there are no major sex-specific regions that are absent from the reference (Suppl. Fig 13A). To test if smaller-scale sequence differences were associated with sex, we performed a GWAS independent of the reference genome using k-mers from the sequencing reads. We found k-mers significantly associated with sex, and when we assembled those k-mers into contigs, they corresponded to *dmrt1*, consistent with the results from SNP-based GWAS (Suppl. Fig. 13B). Smaller copy number variations (CNVs) not captured by genome size estimation or by k-mers could be associated with sex but not be tagged by linked SNPs. To test for this possibility, we compared the frequency of individual CNVs genomewide between the sexes, but none were significantly associated (Suppl. Fig 13C). Although sex chromosomes often carry chromosomal rearrangements, we found no evidence of an inversion in X or Y (Fig. 3D and Methods). These results indicate that, at this level of detection, only a small genomic region <30 kb within otherwise non-sexually differentiated chromosomes (autosomes) distinguish female and male ornamental betta.

Because *dmrt1* had a strong signal of a selective sweep in ornamental betta, we hypothesized that *dmrt1*’s role in sex determination evolved rapidly during domestication. To explore the relationship between *dmrt1* and sex in wild and ornamental betta, we first built a phylogenetic tree of the *dmrt1* locus defined as a ~30 kb linkage-disequilibrium block (Fig. 3E). Consistent with the selective sweep, all ornamental females, but only one wild female, had a particular haplotype we call X_1_. In wild *B. splendens*, 50% (6/12) of XX individuals were female and 91% (10/11; binomial *P*=0.00048) of XY individuals were male (Fig. 3F,G). While this evidence suggests *dmrt1*_Y promotes maleness in wild *B. splendens*, it is possible that multiple sex determination systems segregate in the wild, similar to what is seen in African cichlids (*39*). In contrast, in ornamental betta, 87% (94/108; the 17 fish in the GWAS plus 91 independent samples; binomial *P*<10^−12^) of XX individuals were female and 93% (83/89; binomial *P*<10^−12^) of XY individuals were male (Fig. 3F,G). These results are consistent with a higher penetrance of XX in promoting female development in ornamental betta than in wild *B. splendens* (Fisher’s exact test two-tailed *P*=0.005) and suggest this effect contributed to the selective sweep around *dmrt1.* In line with selection at the *dmrt1* locus occurring preferentially on the X, the ornamental X_1_ haplotype had 33% lower nucleotide diversity than the ornamental Y haplotype. Assuming no sex differences in mutation rates, X has ~3× more opportunity to accumulate mutations than Y, since it is present as two alleles in most females (XX) but only as one in most males (XY). Adjusting by this 3:1 ratio of X to Y, X_1_ has 78% lower diversity than Y. The more marked decrease in diversity on X_1_ supports the hypothesis that selection in ornamental betta has preferentially occurred in the X_1_ haplotype.

Since *dmrt1* XX-XY status was not perfectly related to gonadal sex, we searched for additional sex-linked loci that may have been missed by GWAS. To do so, we performed two quantitative trait locus (QTL) mapping experiments, one in a cross between an XX female and an XY male, and another between an XX female and an XX male. In the XX × XY cross, 52% of the offspring were female and we detected a single sex-linked locus encompassing *dmrt1* (Fig 3B,H). In the XX × XX cross, 90% of the offspring were female and no locus was linked to sex (Fig 3B,H). In the XX × XY cross, 85% of the XX offspring were female and 90% of the XY offspring were male. However, in the XX × XX cross all offspring were XX yet 10% of these fish developed as males, confirming the incomplete penetrance of the XX-XY locus in sex determination, as has been observed in *Oryzias latipes* (medaka fish) that also bear a *dmrt1* XX-XY sex determination system (*40*). In sum, these results confirm that the *dmrt1* locus is strongly linked to sex in ornamental betta, but that XX and XY are neither necessary nor sufficient to determine a particular sex.

To determine whether the X and Y transcripts of *dmrt1* are differentially expressed during sex determination, we performed allele-specific expression analyses in XY ornamental larvae at several time points after fertilization. The results indicated that the *dmrt1* Y allele constitutes 65% of the *dmrt1* mRNA molecules at 4 days post fertilization (dpf) and that this allelic bias progressively decreases at 8 and 12 dpf, until it reverses in adult testis, where only 45% of the *dmrt1* transcripts originate from the Y allele (Fig. 3I). This timing of *dmrt1* XY allele-specific expression is consistent with that of sex determination, since we found that by 4 dpf, XX and XY larvae have started the process of sex differentiation: XY larvae express higher levels of *gonadal soma derived factor* (*gsdf*), a teleost-specific gene essential for testis development (*41*, *42*), and higher levels of *antimullerian hormone* (*amh*), a gene that promotes vertebrate male development (Fig 3J). Each of these genes are the sex determination locus in other fish species (*43*, *44*) and are in separate chromosomes from *dmrt1* in betta, indicating that their sex-specific expression is a response in *trans* to *dmrt1*. Thus, the variants that distinguish *dmrt1* X from Y are associated with higher expression of the *dmrt1* Y allele in a manner that is temporally linked to sex differentiation, further implicating *dmrt1* as the major sex determination gene in ornamental betta.

### Genetic bases of coloration in ornamental betta

Ornamental betta breeders have generated a vast array of fish varieties (e.g. “royal blue”) that differ along multiple axes of coloration: hue, brightness, saturation, and the anatomical distribution of these features. To determine if any of the genes we found to be under strong selection, as well as any others, contribute to coloration in ornamental betta, we performed a GWAS of the red (n=17) and blue (n=17) fish that were used for the selection scans (Fig. 4A; Suppl. Fig. 14A). Red and blue fish lie at opposite ends of the betta hue spectrum and also differ in their brightness and saturation (Fig. 4A,C; Suppl. Fig. 15). However, association mapping alone between pure red and pure blue fish, which are largely fixed for all these color features, cannot establish which of these features are affected by significant loci. Therefore, we also performed a QTL mapping experiment by generating a second-generation (F_2_) hybrid population of red-blue fish in which individual coloration components could segregate (Fig. 4B). In these 211 F_2_ hybrids, we measured the proportion of the anal, caudal and dorsal fins, of the side of the body, and of the head, that was red, blue, or very dark (which we refer to as black). We also measured the hue, brightness, and saturation of the red and blue areas on each body part and used these phenotypes for QTL mapping.

**Figure 4.**
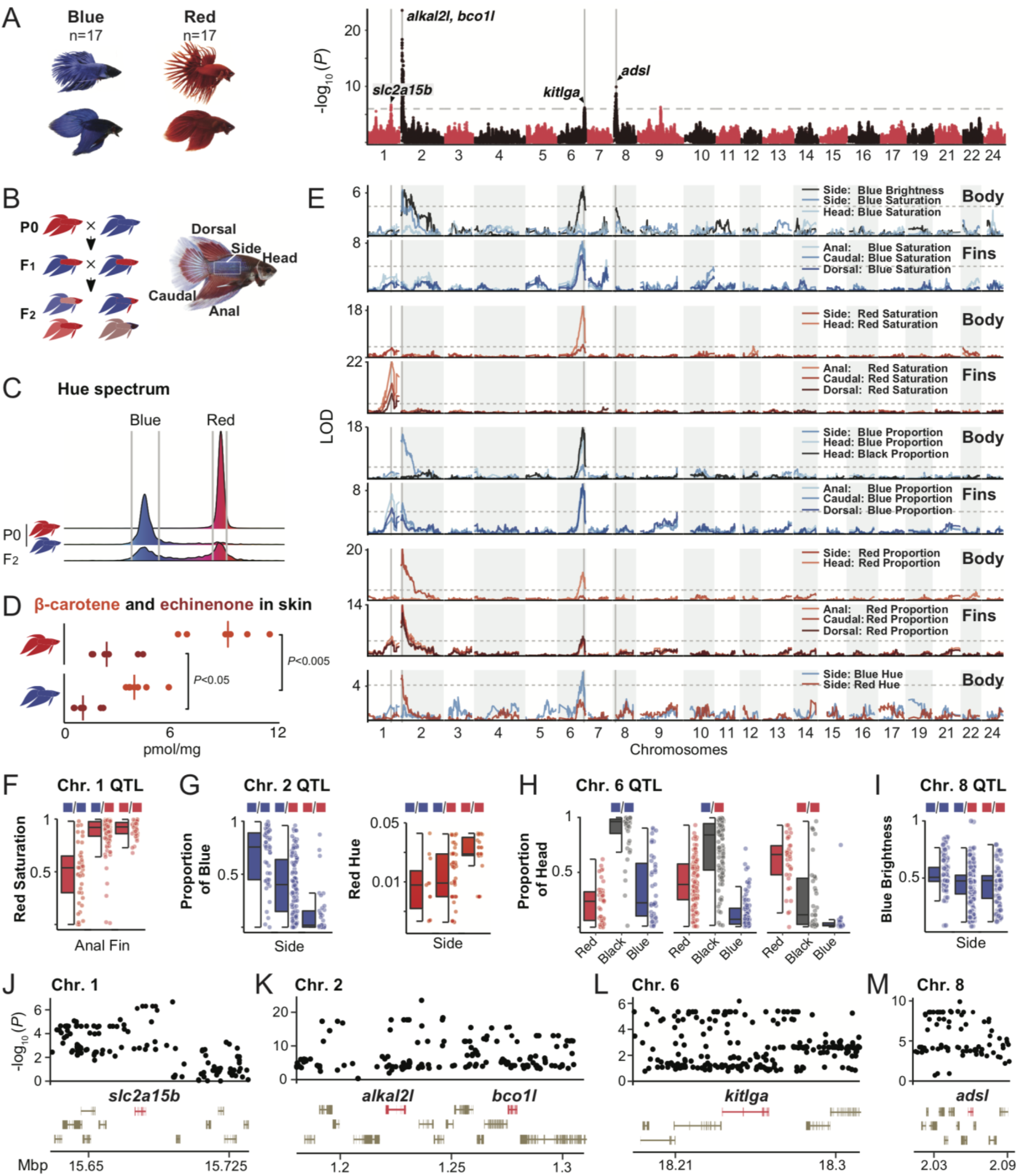
Genomic loci regulating coloration in ornamental betta. **(A)** Manhattan plot for GWAS of color between 17 blue and 17 red ornamental betta. Dashed line denotes the genome-wide significance threshold (see Methods). **(B)** Schematic of the red and blue F2 intercross and photo of a hybrid annotated with the body parts analyzed. **(C)** Distribution of hues observed in the red and blue founder (P0) populations (red, n=13; blue, n=11), and F2 hybrids (n=202). Grey vertical lines depict the hue intervals for red and blue assessed in the F2 population. **(D)** Concentration of β-carotene and echinenone in skin of red and blue ornamental betta. *P* denotes the result of a Mann-Whitney U test. **(E)** QTL mapping of color features across different body parts. Dashed lines denote the genome-wide significance threshold at α=0.05. **(F,G,H,I)** Phenotypic distribution for QTL genotypes across the F2s (red and blue squares denote alleles inherited from red and blue P0’s, respectively). **(J,K,L,M)** GWAS zoom-in with gene annotations across significant loci.

The strongest GWAS signal occurred between *augmentator-α2* (*alkal2l*) and *beta-carotene oxygenase 1-like* (*bco1l*) on chromosome 2 (Fig. 4A,K), a region with a large difference in selection sweep signal between blue and red fish (Fig. 2B). This GWAS peak was aligned with a QTL at which the allele swept in blue fish increased the proportion of blue and decreased the proportion of red on fins and body in the hybrids (Fig 4E,G). Interestingly, this locus modulates blue saturation only on the body and not on the fins or the head (Fig. 4E). *alkal2l* encodes a ligand of Leukocyte Tyrosine Kinase (*33*, *36*) which is expressed in the precursors of iridophores, the chromatophores that generate refractive colors such as blue (*34*, *35*). In zebrafish, *alkal2l* is necessary for iridophore development (*33*). Altogether, this suggests that the large number of iridophores in blue ornamental betta, compared to red fish, is caused by genetic variation affecting this developmental cell-signaling ligand. *alkal2l* likely corresponds to the gene referred to by betta breeders as the *spread iridocyte* gene, hypothesized to increase the prevalence of iridescence throughout the body (*45*).

Notably, the *alkal2l–bco1l* locus also modulated the red hue of the red parts of the body (Fig. 4E,G), suggesting that *bco1l,* which encodes a protein predicted to metabolize orange-red carotenoids, could also be involved in differences between red and blue fish. Through biochemical assays, we found that, as predicted by its sequence homology to other BCO1 proteins, BCO1L has 15,15′-dioxygenase activity that cleaves β-carotene into two molecules of all-trans retinal (Suppl. Fig. 16A-D). Consistent with the QTL effect on red hue and BCO1L biochemical activity, we found that red fish have more β-carotene and echinenone in their skin than blue fish (Fig. 4D). One of the *bco1l* variants most strongly associated with red and blue coloration results in a change from threonine in red fish to isoleucine in blue fish (Suppl. Fig. 14B and Suppl. Fig. 16E,F). We did not detect differential biochemical activity of the two alleles in vitro, but it is possible that their activity, stability, or gene expression differs in vivo (Suppl. Fig. 14C). Therefore, variation in the locus containing *alkal2l* and *bco1l* likely affects both blue and red coloration through these two genes located only ~50 kb apart. The tight linkage might explain why breeders struggle to make the “perfect” red fish without any iridescence.

The second strongest GWAS peak, on chromosome 8, mapped to *adenylosuccinate lyase* (*adsl*), and the strongest QTL at this locus was for the brightness of blue areas on the body (Fig. 4E,I,M). *adsl* encodes an enzyme involved in the de novo synthesis of purines (*46*). Purines are the major components of the reflective platelets in fish skin iridophores that underlie iridescence (*47*), and these platelets differ in structure between blue and red betta fish (*48*). While the homologs of *adsl* have not been previously implicated in animal coloration, mutations in other genes in the de novo purine synthesis pathway cause iridophore defects in zebrafish (*49*). *adsl* likely corresponds to the gene betta breeders refer to as *blue* (*48, 50–52*).

The third strongest GWAS peak, on chromosome 1, mapped to *solute carrier family 2, member 15b* (*slc2a15b*), a gene necessary for the development of larval yellow xanthophores in medaka (*53*), but whose role in adult pigmentation was previously not described (Fig. 4A,J). We found a QTL that overlaps *slc2a15b* that strongly affected the saturation of red areas in the fins, but not of the body or the head (Fig. 4E,F). Intense coloration on the fins relative to the body is a phenotype referred to by breeders as the “Cambodian” variety, and our results suggest *slc2a15b* contributes to this phenotype.

The fourth strongest GWAS peak, on chromosome 6, mapped to *kit ligand* (*kitlga*), whose orthologues affect melanin pigmentation in other fish and in mammals (*54*, *55*) (Fig. 4A,L). A QTL overlapping *kitlga* strongly modulated the proportion of black, blue, and red on the head and fins, but less so on the body (Fig. 4E,H). A black head, a phenotype we found is linked to *kitlga,* is referred to by breeders as the *mask* trait (*7*). This QTL also modified the saturation of blue areas on the fins but not on the body, and had minor effects on red saturation outside the head. Its comparatively stronger impact on blue saturation may be related to the tight histological association of iridophores and melanophores as a unit in betta skin (*48*).

Altogether, we discovered that red-blue variation in ornamental betta is linked to genetic polymorphisms near two genes encoding cell-signaling ligands (*alkal2l* and *kitlga*), two enzymes (*bco1l*, which metabolizes pigments, and *adsl*, which produces material for reflective structures), and a membrane solute transporter (*slc2a15b*). Genes we identified likely correspond to those inferred, but not molecularly identified, by betta geneticists beginning in the 1930s (*52*, *56*). Notably, all of these genes had anatomical specificity, and all but two were on separate chromosomes (Fig. 4A,D).

### Genetic bases of tail morphology in ornamental betta

We found strong signals of selective sweeps in crowntail fish on chromosomes 22 and 24, suggesting these regions could harbor variants associated with crown morphology (Fig. 2D,E). To identify such variants within selective peaks, and elsewhere throughout the genome, we performed a GWAS with the 18 veiltail and 16 crowntail fish used for the selection scans. We found two significant peaks, one on chromosome 22 and another on chromosome 24, overlapping the selection peaks (Fig. 5A), indicating that these regions are not only under selection but are the main loci contributing to differences between veiltail and crowntail fish.

**Figure 5.**
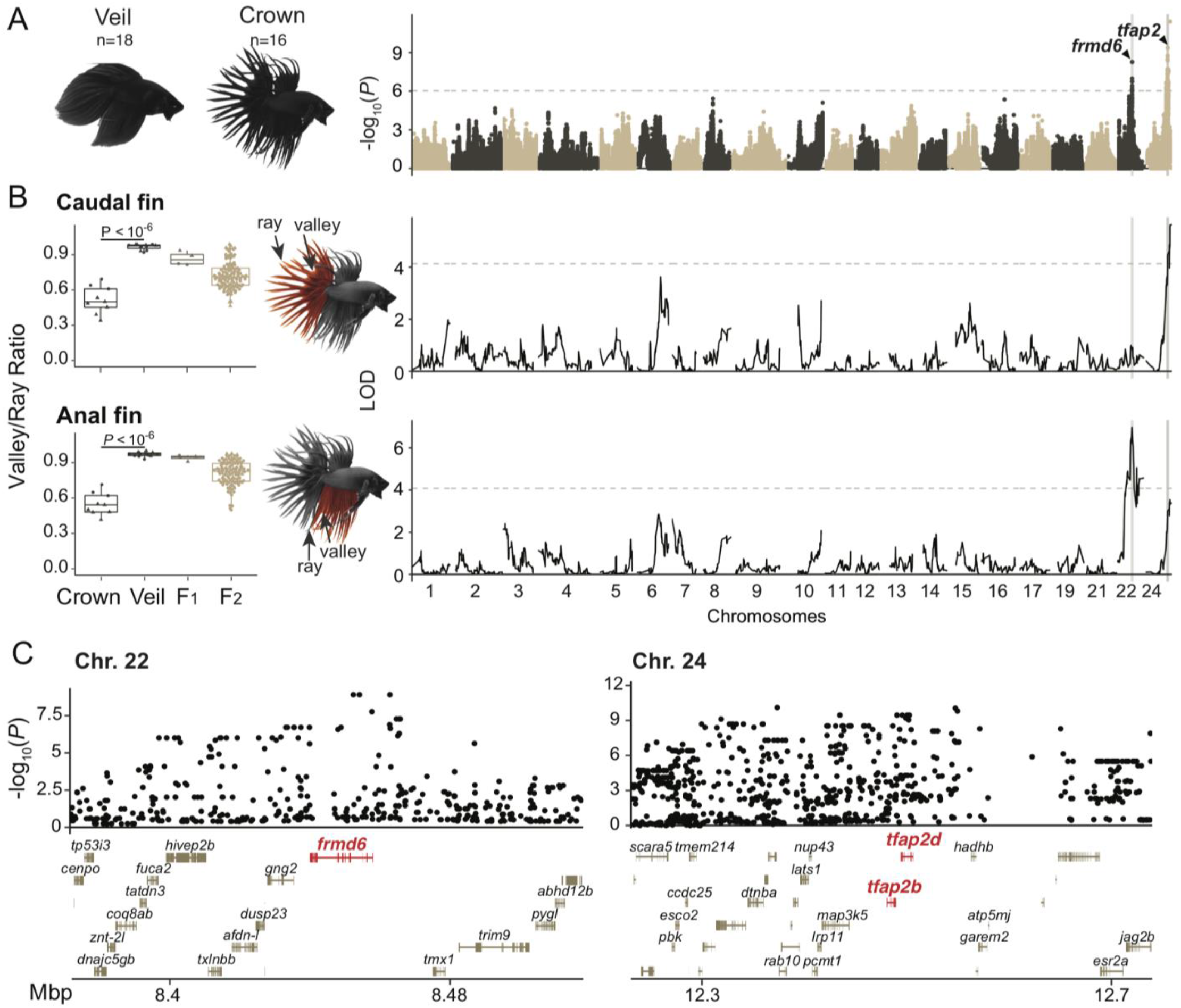
Genomic loci regulating fin morphology in ornamental betta. **(A)** Manhattan plot for GWAS of fin type between 18 veiltail and 16 crowntail ornamental betta. Dashed line denotes the genome-wide significance threshold (see Methods). **(B)** Left: valley/ray ratio in anal and caudal fins across crown (n=9) and veil (n=13) founder populations, F1 (n=4), and F2 hybrids (n=139). *P* denotes the result of a Mann-Whitney U test. Right: QTL mapping of valley/ray ratios for caudal and anal fins. Dashed line denotes the genome-wide significance threshold at α=0.05. **(C)** GWAS zoom-in with gene annotations across significant loci.

To confirm the involvement of the GWAS loci in fin morphology, we performed a QTL mapping experiment in an F_2_-hybrid population from a cross between veil and crowntail fish (Fig. 5B). In agreement with the GWAS results, we found two significant QTLs, one on chromosome 22 and another on chromosome 24, that overlap the GWAS peaks. Surprisingly, we found that the chromosome 22 locus is significantly linked only to anal fin webbing and not caudal fin webbing, whereas the chromosome 24 locus is linked to caudal fin webbing but not significantly linked to anal fin webbing (Fig. 5B). These complementary association and quantitative mapping experiments demonstrate that two loci are the primary determinants of veil–crown morphology and that webbing of different fins is under separate genetic control.

Examining the genes at the crown-veil GWAS peaks identified promising causal genes. The strongest association signal on chromosome 22 maps to *frmd6*, which encodes the protein willin that regulates tissue growth as part of the hippo pathway (*57*) (Fig. 5C,D; Suppl. Fig. 17). The region of strongest association on chromosome 24 is larger and encompasses 22 genes. Of these, *tfap2b* and *tfap2d*, which have evolutionarily ancient roles in ectodermal development (*58*), are prominent candidate genes (Fig. 5C,D; Suppl. Fig. 17). Interestingly, as with the variants that affect coloration, we also find evidence of anatomical modularity for the variants that affect fin morphology. These results also demonstrate that there is no single “crowntail gene”, as had been speculated by ornamental betta breeders (*7*).

## Discussion

Using whole genome sequencing of multiple *Betta* species, populations, and individuals, we take an important first step in unraveling the domestication history of betta fish. Our results suggest that betta were domesticated more than 1,000 years ago —at least three centuries earlier than previously suggested. While domesticated betta are largely derived from *Betta splendens*, they carry genetic contributions from two other species that are also endemic to the Malay Peninsula: *B. imbellis* and *B. mahachaiensis*. None of the alleles derived from these other *Betta species* are present in all ornamental individuals, nor do they contribute to the regions under selection driving sex determination, coloration, or fin morphologies (Supp. Fig 12). These introgressed alleles, however, might contribute to other traits of domesticated betta or represent historical attempts by breeders to introduce new phenotypes into ornamental fish through hybridization.

The strongest genetic evidence of selection during domestication involves *dmrt1*, which we discover is the sex determination gene in ornamental betta. Most females are XX and most males are XY, settling a long-standing question in the field (*59*). A selective sweep of a *dmrt1_*X allele with increased penetrance may have been selected by breeders, since it would lead to more predictable sex ratios in spawns. The lower penetrance of *dmrt1* on sex determination in wild *B. splendens* suggests that additional sex determination loci that operate in the wild are not present in domesticated betta, similar to what is seen in zebrafish (*60*). In contrast to domesticated zebrafish, where sex is not determined by a single locus, in ornamental betta sex is predominantly determined by a single large-effect locus that maps to *dmrt1*.

In poeciliid fishes such as guppies and swordtails, the sex determination locus is linked to multiple color genes that contribute to sexually dimorphic coloration and shape the genetics of female preference for male color traits (*61*). In contrast, the betta sex determination locus is only ~30 kb in size and is not linked to genes known to affect color. These results are consistent with coloration not being particularly sexually dimorphic in betta. Instead, we find that color and fin morphology in betta has a lego-like logic, in which major-effect genes located on different chromosomes modulate color and fin morphology with surprising anatomical specificity (Suppl. Table 2). Betta breeders are keenly aware of the mix-and-match possibilities of betta, and leverage this feature to breed new fish varieties by combining different body, head, and fin colors with various fin morphologies.

Our results provide molecular entry points for further study of the developmental and evolutionary bases of change in morphology and sex determination. The genomic resources we generated will also enable genetic studies into how centuries of artificial selection of betta for fighting purposes have shaped their aggression and other fighting-related traits. Altogether, our work elucidates the genomic consequences of the domestication of ornamental betta and helps establish this fish as a modern system for evolutionary genetic interrogation.

## Acknowledgments

The DNA pipelines staff at the Wellcome Sanger Institute generated sequencing data. Debbie Leung and Hiroki Tomida photographed fish. Ronny Kyller as well as members of the International Betta Congress including Liz Hahn, Sieg Illig, Karen MacAuley, and Holly Rutan provided samples. Leo Buss, Darcy Kelley, Carol Mason, and Molly Przeworski provided comments on the manuscript.

## Funding

Searle Scholarship and Sloan Foundation Fellowship (AB). Wellcome grant WT206194 (IB, JW, SM, WC, KH, RD). Wellcome grant WT207492 (SM, RD). Flemish University Research Fund (JC-G, HS). FWO Research Foundation Flanders Ph.D. fellowship (NV). National Institutes of Health grant EY020551 (JvL). National Institutes of Health grant EY028121 (JvL).

## Author contributions

Conceptualization: AB, RD, YMK, HS. Formal analysis: AB, IB, WC, JC-G, KH, YMK, SM, HS, NV, JW. Funding acquisition: AB, RD, JvL, HS. Investigation: AB, SB, KXF, CH, YMK, MRL, HS. Project administration: AB, RD, HS. Resources: HHT, LR. Software: JC-G, YMK, HS, NV. Supervision: AB, RD, JvL, HS. Visualization: SB, YMK, HS, NV. Writing – original draft: AB, YMK, HS. Writing – review & editing: AB, RD, CH, KH, YMK, JvL, MRL, LR, HS.

## Competing interests

Authors declare that they have no competing interests.

## Data and materials availability

Data used in the analysis are available in NCBI GenBank GCA_900634795.3 and BioProject PRJNA486171.

## Notes

### Competing Interest Statement

The authors have declared no competing interest.

## References and Notes

1. L. A. F. Frantz, D. G. Bradley, G. Larson, L. Orlando, Animal domestication in the era of ancient genomics. Nat. Rev. Genet. 21, 449–460 (2020).

2. C.-J. Rubin, M. C. Zody, J. Eriksson, J. R. S. Meadows, E. Sherwood, M. T. Webster, L. Jiang, M. Ingman, T. Sharpe, S. Ka, F. Hallböök, F. Besnier, O. Carlborg, B. Bed’hom, M. Tixier-Boichard, P. Jensen, P. Siegel, K. Lindblad-Toh, L. Andersson, Whole-genome resequencing reveals loci under selection during chicken domestication. Nature. 464, 587–591 (2010).

3. P. Xu, X. Zhang, X. Wang, J. Li, G. Liu, Y. Kuang, J. Xu, X. Zheng, L. Ren, G. Wang, Y. Zhang, L. Huo, Z. Zhao, D. Cao, C. Lu, C. Li, Y. Zhou, Z. Liu, Z. Fan, G. Shan, X. Li, S. Wu, L. Song, G. Hou, Y. Jiang, Z. Jeney, D. Yu, L. Wang, C. Shao, L. Song, J. Sun, P. Ji, J. Wang, Q. Li, L. Xu, F. Sun, J. Feng, C. Wang, S. Wang, B. Wang, Y. Li, Y. Zhu, W. Xue, L. Zhao, J. Wang, Y. Gu, W. Lv, K. Wu, J. Xiao, J. Wu, Z. Zhang, J. Yu, X. Sun, Genome sequence and genetic diversity of the common carp, *Cyprinus carpio*. Nat. Genet. 46, 1212–1219 (2014).

4. Z. Chen, Y. Omori, S. Koren, T. Shirokiya, T. Kuroda, A. Miyamoto, H. Wada, A. Fujiyama, A. Toyoda, S. Zhang, T. G. Wolfsberg, K. Kawakami, A. M. Phillippy, NISC Comparative Sequencing Program, J. C. Mullikin, S. M. Burgess, De novo assembly of the goldfish (*Carassius auratus*) genome and the evolution of genes after whole-genome duplication. Sci Adv. 5, eaav0547 (2019).

5. D. Chen, Q. Zhang, W. Tang, Z. Huang, G. Wang, Y. Wang, J. Shi, H. Xu, L. Lin, Z. Li, W. Chi, L. Huang, J. Xia, X. Zhang, L. Guo, Y. Wang, P. Ma, J. Tang, G. Zhou, M. Liu, F. Liu, X. Hua, B. Wang, Q. Shen, Q. Jiang, J. Lin, X. Chen, H. Wang, M. Dou, L. Liu, H. Pan, Y. Qi, B. Wu, J. Fang, Y. Zhou, W. Cen, W. He, Q. Zhang, T. Xue, G. Lin, W. Zhang, Z. Liu, L. Qu, A. Wang, Q. Ye, J. Chen, Y. Zhang, R. Ming, M. Van Montagu, H. Tang, Y. Van de Peer, Y. Chen, J. Zhang, The evolutionary origin and domestication history of goldfish (*Carassius auratus*). Proc. Natl. Acad. Sci. U. S. A. 117, 29775–29785 (2020).

6. M. Bekoff, Encyclopedia of Human-Animal Relationships (Greenwood Press, 2007).

7. M. Brammah, The Betta Bible (CreateSpace Independent Publishing Platform, 2015).

8. L. Rüber, R. Britz, R. Zardoya, Molecular phylogenetics and evolutionary diversification of labyrinth fishes (Perciformes: *Anabantoidei*). Syst. Biol. 55, 374–397 (2006).

9. N. Sriwattanarothai, D. Steinke, P. Ruenwongsa, R. Hanner, B. Panijpan, Molecular and morphological evidence supports the species status of the Mahachai fighter *Betta* sp. Mahachai and reveals new species of *Betta* from Thailand. J. Fish Biol. 77, 414–424 (2010).

10. International Betta Congress, About *Betta splendens*. IBC (2020), (available at https://www.ibcbettas.org/about-betta-splendens/).

11. G. Fan, J. Chan, K. Ma, B. Yang, H. Zhang, X. Yang, C. Shi, H. Chun-Hin Law, Z. Ren, Q. Xu, Q. Liu, J. Wang, W. Chen, L. Shao, D. Gonçalves, A. Ramos, S. D. Cardoso, M. Guo, J. Cai, X. Xu, J. Wang, H. Yang, X. Liu, Y. Wang, Chromosome-level reference genome of the Siamese fighting fish *Betta splendens*, a model species for the study of aggression. Gigascience. 7 (2018), doi:10.1093/gigascience/giy087.

12. S. Prost, M. Petersen, M. Grethlein, S. J. Hahn, N. Kuschik-Maczollek, M. E. Olesiuk, J.-O. Reschke, T. E. Schmey, C. Zimmer, D. K. Gupta, T. Schell, R. Coimbra, J. De Raad, F. Lammers, S. Winter, A. Janke, Improving the chromosome-level genome assembly of the Siamese fighting fish (*Betta splendens*) in a university master’s course. G3 Genes|Genomes|Genetics. 10, 2179–2183 (2020).

13. L. Wang, F. Sun, Z. Y. Wan, B. Ye, Y. Wen, H. Liu, Z. Yang, H. Pang, Z. Meng, B. Fan, Y. Alfiko, Y. Shen, B. Bai, M. S. Q. Lee, F. Piferrer, M. Schartl, A. Meyer, G. H. Yue, Genomic basis of striking fin shapes and colours in the fighting fish. Mol. Biol. Evol. (2021), doi:10.1093/molbev/msab110.

14. N. L. Bennington, Germ cell origin and spermatogenesis in the Siamese fighting fish, *Betta splendens*. J. Morphol. 60, 103–125 (1936).

15. A. Rhie, S. A. McCarthy, O. Fedrigo, J. Damas, G. Formenti, S. Koren, M. Uliano-Silva, W. Chow, A. Fungtammasan, G. L. Gedman, L. J. Cantin, F. Thibaud-Nissen, L. Haggerty, C. Lee, B. J. Ko, J. Kim, I. Bista, M. Smith, B. Haase, J. Mountcastle, S. Winkler, S. Paez, J. Howard, S. C. Vernes, T. M. Lama, F. Grutzner, W. C. Warren, C. Balakrishnan, D. Burt, J. M. George, M. Biegler, D. Iorns, A. Digby, D. Eason, T. Edwards, M. Wilkinson, G. Turner, A. Meyer, A. F. Kautt, P. Franchini, H. William Detrich, H. Svardal, M. Wagner, G. J. P. Naylor, M. Pippel, M. Malinsky, M. Mooney, M. Simbirsky, B. T. Hannigan, T. Pesout, M. Houck, A. Misuraca, S. B. Kingan, R. Hall, Z. Kronenberg, J. Korlach, I. Sović, C. Dunn, Z. Ning, A. Hastie, J. Lee, S. Selvaraj, R. E. Green, N. H. Putnam, J. Ghurye, E. Garrison, Y. Sims, J. Collins, S. Pelan, J. Torrance, A. Tracey, J. Wood, D. Guan, S. E. London, D. F. Clayton, C. V. Mello, S. R. Friedrich, P. V. Lovell, E. Osipova, F. O. Al-Ajli, S. Secomandi, H. Kim, C. Theofanopoulou, Y. Zhou, R. S. Harris, K. D. Makova, P. Medvedev, J. Hoffman, P. Masterson, K. Clark, F. Martin, K. Howe, P. Flicek, B. P. Walenz, W. Kwak, H. Clawson, M. Diekhans, L. Nassar, B. Paten, R. H. S. Kraus, H. Lewin, A. J. Crawford, M. T. P. Gilbert, G. Zhang, B. Venkatesh, R. W. Murphy, K.-P. Koepfli, B. Shapiro, W. E. Johnson, F. Di Palma, T. Margues-Bonet, E. C. Teeling, T. Warnow, J. M. Graves, O. A. Ryder, D. Hausler, S. J. O’Brien, K. Howe, E. W. Myers, R. Durbin, A. M. Phillippy, E. D. Jarvis, Towards complete and error-free genome assemblies of all vertebrate species. Nature, In press. (2021), doi:10.1101/2020.05.22.110833

16. N. Patterson, P. Moorjani, Y. Luo, S. Mallick, N. Rohland, Y. Zhan, T. Genschoreck, T. Webster, D. Reich, Ancient admixture in human history. Genetics. 192, 1065–1093 (2012).

17. M. Malinsky, H. Svardal, A. M. Tyers, E. A. Miska, M. J. Genner, G. F. Turner, R. Durbin, Whole-genome sequences of Malawi cichlids reveal multiple radiations interconnected by gene flow. Nat Ecol Evol. 2, 1940–1955 (2018).

18. F. L. Wu, A. I. Strand, L. A. Cox, C. Ober, J. D. Wall, P. Moorjani, M. Przeworski, A comparison of humans and baboons suggests germline mutation rates do not track cell divisions. PLoS Biol. 18, e3000838 (2020).

19. F. Schlamp, J. van der Made, R. Stambler, L. Chesebrough, A. R. Boyko, P. W. Messer, Evaluating the performance of selection scans to detect selective sweeps in domestic dogs. Mol. Ecol. 25, 342–356 (2016).

20. A. M. Harris, N. R. Garud, M. DeGiorgio, Detection and classification of hard and soft sweeps from unphased genotypes by multilocus genotype identity. Genetics. 210, 1429–1452 (2018).

21. A. Harris, P. Siggers, S. Corrochano, N. Warr, D. Sagar, D. T. Grimes, M. Suzuki, R. D. Burdine, F. Cong, B.-K. Koo, H. Clevers, I. Stévant, S. Nef, S. Wells, R. Brauner, B. Ben Rhouma, N. Belguith, C. Eozenou, J. Bignon-Topalovic, A. Bashamboo, K. McElreavey, A. Greenfield, ZNRF3 functions in mammalian sex determination by inhibiting canonical WNT signaling. Proc. Natl. Acad. Sci. U. S. A. 115, 5474–5479 (2018).

22. E. Szenker-Ravi, U. Altunoglu, M. Leushacke, C. Bosso-Lefèvre, M. Khatoo, H. Thi Tran, T. Naert, R. Noelanders, A. Hajamohideen, C. Beneteau, S. B. de Sousa, B. Karaman, X. Latypova, S. Başaran, E. B. Yücel, T. T. Tan, L. Vlaminck, S. S. Nayak, A. Shukla, K. M. Girisha, C. Le Caignec, N. Soshnikova, Z. O. Uyguner, K. Vleminckx, N. Barker, H. Kayserili, B. Reversade, RSPO2 inhibition of RNF43 and ZNRF3 governs limb development independently of LGR4/5/6. Nature. 557, 564–569 (2018).

23. Y. Tatsumi, M. Takeda, M. Matsuda, T. Suzuki, H. Yokoi, TALEN-mediated mutagenesis in zebrafish reveals a role for r-spondin 2 in fin ray and vertebral development. FEBS Lett. 588, 4543–4550 (2014).

24. U. F. Mustapha, D.-N. Jiang, Z.-H. Liang, H.-T. Gu, W. Yang, H.-P. Chen, S.-P. Deng, T.-L. Wu, C.-X. Tian, C.-H. Zhu, G.-L. Li, Male-specific Dmrt1 is a candidate sex determination gene in spotted scat (*Scatophagus argus*). Aquaculture. 495, 351–358 (2018).

25. I. Nanda, M. Kondo, U. Hornung, S. Asakawa, C. Winkler, A. Shimizu, Z. Shan, T. Haaf, N. Shimizu, A. Shima, M. Schmid, M. Schartl, A duplicated copy of DMRT1 in the sex-determining region of the Y chromosome of the medaka, *Oryzias latipes*. Proc. Natl. Acad. Sci. U. S. A. 99, 11778–11783 (2002).

26. Z. Cui, Y. Liu, W. Wang, Q. Wang, N. Zhang, F. Lin, N. Wang, C. Shao, Z. Dong, Y. Li, Y. Yang, M. Hu, H. Li, F. Gao, Z. Wei, L. Meng, Y. Liu, M. Wei, Y. Zhu, H. Guo, C. H. K. Cheng, M. Schartl, S. Chen, Genome editing reveals *dmrt1* as an essential male sex-determining gene in Chinese tongue sole (*Cynoglossus semilaevis*). Sci. Rep. 7, 42213 (2017).

27. S. Yoshimoto, N. Ikeda, Y. Izutsu, T. Shiba, N. Takamatsu, M. Ito, Opposite roles of DMRT1 and its W-linked paralogue, DM-W, in sexual dimorphism of *Xenopus laevis*: implications of a ZZ/ZW-type sex-determining system. Development. 137, 2519–2526 (2010).

28. C. A. Smith, K. N. Roeszler, T. Ohnesorg, D. M. Cummins, P. G. Farlie, T. J. Doran, A. H. Sinclair, The avian Z-linked gene DMRT1 is required for male sex determination in the chicken. Nature. 461, 267–271 (2009).

29. L. Cal, P. Suarez-Bregua, J. M. Cerdá-Reverter, I. Braasch, J. Rotllant, Fish pigmentation and the melanocortin system. Comp. Biochem. Physiol. A Mol. Integr. Physiol. 211, 26–33 (2017).

30. J. Liu, C. Lin, A. Gleiberman, K. A. Ohgi, T. Herman, H. P. Huang, M. J. Tsai, M. G. Rosenfeld, Tbx19, a tissue-selective regulator of POMC gene expression. Proc. Natl. Acad. Sci. U. S. A. 98, 8674–8679 (2001).

31. I. Ziegler, The pteridine pathway in zebrafish: regulation and specification during the determination of neural crest cell-fate. Pigment Cell Res. 16, 172–182 (2003).

32. A. G. Reaume, D. A. Knecht, A. Chovnick, The rosy locus in *Drosophila melanogaster*: xanthine dehydrogenase and eye pigments. Genetics. 129, 1099–1109 (1991).

33. E. S. Mo, Q. Cheng, A. V. Reshetnyak, J. Schlessinger, S. Nicoli, Alk and Ltk ligands are essential for iridophore development in zebrafish mediated by the receptor tyrosine kinase Ltk. Proc. Natl. Acad. Sci. U. S. A. 114, 12027–12032 (2017).

34. A. Fadeev, P. Mendoza-Garcia, U. Irion, J. Guan, K. Pfeifer, S. Wiessner, F. Serluca, A. P. Singh, C. Nüsslein-Volhard, R. H. Palmer, ALKALs are in vivo ligands for ALK family receptor tyrosine kinases in the neural crest and derived cells. Proc. Natl. Acad. Sci. U. S. A. 115, E630–E638 (2018).

35. S. S. Lopes, X. Yang, J. Müller, T. J. Carney, A. R. McAdow, G.-J. Rauch, A. S. Jacoby, L. D. Hurst, M. Delfino-Machín, P. Haffter, R. Geisler, S. L. Johnson, A. Ward, R. N. Kelsh, Leukocyte tyrosine kinase functions in pigment cell development. PLoS Genet. 4, e1000026 (2008).

36. A. V. Reshetnyak, P. B. Murray, X. Shi, E. S. Mo, J. Mohanty, F. Tome, H. Bai, M. Gunel, I. Lax, J. Schlessinger, Augmentor α and β (FAM150) are ligands of the receptor tyrosine kinases ALK and LTK: Hierarchy and specificity of ligand-receptor interactions. Proc. Natl. Acad. Sci. U. S. A. 112, 15862–15867 (2015).

37. H. Helgeland, M. Sodeland, N. Zoric, J. S. Torgersen, F. Grammes, J. von Lintig, T. Moen, S. Kjøglum, S. Lien, D. I. Våge, Genomic and functional gene studies suggest a key role of beta-carotene oxygenase 1 like (*bco1l*) gene in salmon flesh color. Sci. Rep. 9, 20061 (2019).

38. M. A. Gazda, P. M. Araújo, R. J. Lopes, M. B. Toomey, P. Andrade, S. Afonso, C. Marques, L. Nunes, P. Pereira, S. Trigo, G. E. Hill, J. C. Corbo, M. Carneiro, A genetic mechanism for sexual dichromatism in birds. Science. 368, 1270–1274 (2020).

39. W. J. Gammerdinger, T. D. Kocher, Unusual diversity of sex chromosomes in African cichlid fishes. Genes. 9 (2018), doi:10.3390/genes9100480.

40. I. Nanda, U. Hornung, M. Kondo, M. Schmid, M. Schartl, Common spontaneous sex-reversed XX males of the medaka *Oryzias latipes*. Genetics. 163, 245–251 (2003).

41. T. Myosho, H. Otake, H. Masuyama, M. Matsuda, Y. Kuroki, A. Fujiyama, K. Naruse, S. Hamaguchi, M. Sakaizumi, Tracing the emergence of a novel sex-determining gene in medaka, *Oryzias luzonensis*. Genetics. 191, 163–170 (2012).

42. T. Imai, K. Saino, M. Matsuda, Mutation of gonadal soma-derived factor induces medaka XY gonads to undergo ovarian development. Biochem. Biophys. Res. Commun. 467, 109–114 (2015).

43. H. Kaneko, S. Ijiri, T. Kobayashi, H. Izumi, Y. Kuramochi, D.-S. Wang, S. Mizuno, Y. Nagahama, Gonadal soma-derived factor (*gsdf*), a TGF-beta superfamily gene, induces testis differentiation in the teleost fish *Oreochromis niloticus*. Mol. Cell. Endocrinol. 415, 87–99 (2015).

44. R. S. Hattori, Y. Murai, M. Oura, S. Masuda, S. K. Majhi, T. Sakamoto, J. I. Fernandino, G. M. Somoza, M. Yokota, C. A. Strüssmann, A Y-linked anti-Müllerian hormone duplication takes over a critical role in sex determination. Proc. Natl. Acad. Sci. U. S. A. 109, 2955–2959 (2012).

45. G. A. Lucas, A study of variation in the Siamese fighting fish, *Betta splendens*, with emphasis on color mutants and the problem of sex determination (1968) (available at https://lib.dr.iastate.edu/cgi/viewcontent.cgi?article=4488&context=rtd).

46. S. Ratner, in The Enzymes, P. D. Boyer, Ed. (Academic Press, 1972), vol. 7, pp. 167–197.

47. N. M. LeDouarin, C. Kalcheim, The Neural Crest (Cambridge University Press, 1999).

48. G. Khoo, T. M. Lim, V. P. E. Phang, Cellular basis of metallic iridescence in the Siamese fighting fish, *Betta splendens*. The Israeli Journal of Aquaculture-Bamidgeh (2014) (available at https://evols.library.manoa.hawaii.edu/handle/10524/49087).

49. A. Ng, R. A. Uribe, L. Yieh, R. Nuckels, J. M. Gross, Zebrafish mutations in *gart* and *paics* identify crucial roles for de novo purine synthesis in vertebrate pigmentation and ocular development. Development. 136, 2601–2611 (2009).

50. K. Umrath, Über die vererbung der farben und des geschlechts beim schleierkampffisch, Betta splendens. Z. Indukt. Abstamm. Vererbungsl. 77, 450–454 (1939).

51. K. Eberhardt, Die vererbung der farben bei *Betta splendens* Regan. Z. Indukt. Abstamm. Vererbungsl. 79, 548–560 (1941).

52. H. M. Wallbrunn, Genetics of the Siamese fighting fish, *Betta splendens*. Genetics. 43, 289–298 (1958).

53. T. Kimura, Y. Nagao, H. Hashimoto, Y.-I. Yamamoto-Shiraishi, S. Yamamoto, T. Yabe, S. Takada, M. Kinoshita, A. Kuroiwa, K. Naruse, Leucophores are similar to xanthophores in their specification and differentiation processes in medaka. Proc. Natl. Acad. Sci. U. S. A. 111, 7343–7348 (2014).

54. A. Slominski, D. J. Tobin, S. Shibahara, J. Wortsman, Melanin pigmentation in mammalian skin and its hormonal regulation. Physiol. Rev. 84, 1155–1228 (2004).

55. K. A. Hultman, E. H. Budi, D. C. Teasley, A. Y. Gottlieb, D. M. Parichy, S. L. Johnson, Defects in ErbB-dependent establishment of adult melanocyte stem cells reveal independent origins for embryonic and regeneration melanocytes. PLoS Genet. 5, e1000544 (2009).

56. H. B. Goodrich, R. N. Mercer, Genetics and colors of the Siamese fighting fish, *Betta splendens*. Science. 79, 318–319 (1934).

57. M. Sudol, K. F. Harvey, Modularity in the Hippo signaling pathway. Trends Biochem. Sci. 35, 627–633 (2010).

58. T. L. Hoffman, A. L. Javier, S. A. Campeau, R. D. Knight, T. F. Schilling, Tfap2 transcription factors in zebrafish neural crest development and ectodermal evolution. J. Exp. Zool. B Mol. Dev. Evol. 308, 679–691 (2007).

59. T. P. Lowe, J. R. Larkin, Sex reversal in *Betta splendens* Regan with emphasis on the problem of sex determination. J. Exp. Zool. 191, 25–32 (1975).

60. C. A. Wilson, S. K. High, B. M. McCluskey, A. Amores, Y.-L. Yan, T. A. Titus, J. L. Anderson, P. Batzel, M. J. Carvan 3rd, M. Schartl, J. H. Postlethwait, Wild sex in zebrafish: loss of the natural sex determinant in domesticated strains. Genetics. 198, 1291–1308 (2014).

61. A. Lindholm, F. Breden, Sex chromosomes and sexual selection in poeciliid fishes. Am. Nat. 160 Suppl 6, S214–24 (2002).

62. M. Malinsky, R. J. Challis, A. M. Tyers, S. Schiffels, Y. Terai, B. P. Ngatunga, E. A. Miska, R. Durbin, M. J. Genner, G. F. Turner, Genomic islands of speciation separate cichlid ecomorphs in an East African crater lake. Science. 350, 1493–1498 (2015).

63. A. M. Bolger, M. Lohse, B. Usadel, Trimmomatic: a flexible trimmer for Illumina sequence data. Bioinformatics. 30, 2114–2120 (2014).

64. C. Kowasupat, B. Panijpan, P. Ruenwongsa, T. Jeenthong, *Betta siamorientalis*, a new species of bubble-nest building fighting fish (Teleostei: Osphronemidae) from eastern Thailand. Vertebr. Zool. 62, 387–397 (2012).

65. A. Monvises, B. Nuangsaeng, N. Sriwattanarothai, B. Panijpan, The Siamese fighting fish: well-known generally but little-known scientifically. Sci. Asia. 35, 8–16 (2009).

66. C.-S. Chin, P. Peluso, F. J. Sedlazeck, M. Nattestad, G. T. Concepcion, A. Clum, C. Dunn, R. O’Malley, R. Figueroa-Balderas, A. Morales-Cruz, G. R. Cramer, M. Delledonne, C. Luo, J. R. Ecker, D. Cantu, D. R. Rank, M. C. Schatz, Phased diploid genome assembly with single-molecule real-time sequencing. Nat. Methods. 13, 1050–1054 (2016).

67. A. C. English, S. Richards, Y. Han, M. Wang, V. Vee, J. Qu, X. Qin, D. M. Muzny, J. G. Reid, K. C. Worley, R. A. Gibbs, Mind the gap: upgrading genomes with Pacific Biosciences RS long-read sequencing technology. PLoS One. 7, e47768 (2012).

68. E. Garrison, G. Marth, Haplotype-based variant detection from short-read sequencing. arXiv (2012), (available at http://arxiv.org/abs/1207.3907).

69. J. M. Flynn, R. Hubley, C. Goubert, J. Rosen, A. G. Clark, C. Feschotte, A. F. Smit, RepeatModeler2 for automated genomic discovery of transposable element families. Proc. Natl. Acad. Sci. U. S. A. 117, 9451–9457 (2020).

70. A. L. Price, N. C. Jones, P. A. Pevzner, De novo identification of repeat families in large genomes. Bioinformatics. 21 Suppl 1, i351–8 (2005).

71. Z. Bao, S. R. Eddy, Automated de novo identification of repeat sequence families in sequenced genomes. Genome Res. 12, 1269–1276 (2002).

72. S. Ou, N. Jiang, LTR_retriever: A highly accurate and sensitive program for identification of long terminal repeat retrotransposons. Plant Physiology. 176 (2018), pp. 1410–1422.

73. D. Ellinghaus, S. Kurtz, U. Willhoeft, LTRharvest, an efficient and flexible software for de novo detection of LTR retrotransposons. BMC Bioinformatics. 9, 18 (2008).

74. Smit, F. A. A., Repeat-Masker Open-3.0. http://www.repeatmasker.org (2004) (available at https://ci.nii.ac.jp/naid/10029514778/).

75. G. Marçais, A. L. Delcher, A. M. Phillippy, R. Coston, S. L. Salzberg, A. Zimin, MUMmer4: A fast and versatile genome alignment system. PLoS Comput. Biol. 14, e1005944 (2018).

76. M. Chakraborty, N. W. VanKuren, R. Zhao, X. Zhang, S. Kalsow, J. J. Emerson, Hidden genetic variation shapes the structure of functional elements in *Drosophila*. Nat. Genet. 50, 20–25 (2018).

77. H. Li, Aligning sequence reads, clone sequences and assembly contigs with BWA-MEM. arXiv (2013), (available at http://arxiv.org/abs/1303.3997).

78. Broad Institute, “Picard Toolkit”, Broad institute, GitHub repository. Picard Toolkit (2019), (available at http://broadinstitute.github.io/picard/).

79. H. Li, A statistical framework for SNP calling, mutation discovery, association mapping and population genetical parameter estimation from sequencing data. Bioinformatics. 27, 2987–2993 (2011).

80. A. McKenna, M. Hanna, E. Banks, A. Sivachenko, K. Cibulskis, A. Kernytsky, K. Garimella, D. Altshuler, S. Gabriel, M. Daly, M. A. DePristo, The Genome Analysis Toolkit: a MapReduce framework for analyzing next-generation DNA sequencing data. Genome Res. 20, 1297–1303 (2010).

81. J. T. Robinson, H. Thorvaldsdóttir, W. Winckler, M. Guttman, E. S. Lander, G. Getz, J. P. Mesirov, Integrative genomics viewer. Nat. Biotechnol. 29, 24–26 (2011).

82. M. Martin, M. Patterson, S. Garg, S. O. Fischer, N. Pisanti, G. W. Klau, A. Schöenhuth, T. Marschall, WhatsHap: fast and accurate read-based phasing. Cold Spring Harbor Laboratory (2016), p. 085050.

83. O. Delaneau, J.-F. Zagury, M. R. Robinson, J. L. Marchini, E. T. Dermitzakis, Accurate, scalable and integrative haplotype estimation. Nat. Commun. 10, 5436 (2019).

84. Y. M. Kwon, K. Gori, N. Park, N. Potts, K. Swift, J. Wang, M. R. Stammnitz, N. Cannell, A. Baez-Ortega, S. Comte, S. Fox, C. Harmsen, S. Huxtable, M. Jones, A. Kreiss, C. Lawrence, B. Lazenby, S. Peck, R. Pye, G. Woods, M. Zimmermann, D. C. Wedge, D. Pemberton, M. R. Stratton, R. Hamede, E. P. Murchison, Evolution and lineage dynamics of a transmissible cancer in Tasmanian devils. PLoS Biol. 18, e3000926 (2020).

85. S. Purcell, B. Neale, K. Todd-Brown, L. Thomas, M. A. R. Ferreira, D. Bender, J. Maller, P. Sklar, P. I. W. de Bakker, M. J. Daly, P. C. Sham, PLINK: a tool set for whole-genome association and population-based linkage analyses. Am. J. Hum. Genet. 81, 559–575 (2007).

86. B. Q. Minh, H. A. Schmidt, O. Chernomor, D. Schrempf, M. D. Woodhams, A. von Haeseler, R. Lanfear, IQ-TREE 2: new models and efficient methods for phylogenetic inference in the genomic era. Mol. Biol. Evol. 37, 1530–1534 (2020).

87. S. Kalyaanamoorthy, B. Q. Minh, T. K. F. Wong, A. von Haeseler, L. S. Jermiin, ModelFinder: fast model selection for accurate phylogenetic estimates. Nat. Methods. 14, 587–589 (2017).

88. M. Malinsky, M. Matschiner, H. Svardal, Dsuite - Fast D-statistics and related admixture evidence from VCF files. Mol. Ecol. Resour. 21, 584–595 (2021).

89. G. Nilsen, K. Liestøl, P. Van Loo, H. K. Moen Vollan, M. B. Eide, O. M. Rueda, S.-F. Chin, R. Russell, L. O. Baumbusch, C. Caldas, A.-L. Børresen-Dale, O. C. Lingjaerde, Copynumber: efficient algorithms for single- and multi-track copy number segmentation. BMC Genomics. 13, 591 (2012).

90. A. Bergström, S. A. McCarthy, R. Hui, M. A. Almarri, Q. Ayub, P. Danecek, Y. Chen, S. Felkel, P. Hallast, J. Kamm, H. Blanché, J.-F. Deleuze, H. Cann, S. Mallick, D. Reich, M. S. Sandhu, P. Skoglund, A. Scally, Y. Xue, R. Durbin, C. Tyler-Smith, Insights into human genetic variation and population history from 929 diverse genomes. Science. 367 (2020), doi:10.1126/science.aay5012.

91. K. Ulm, A simple method to calculate the confidence interval of a standardized mortality ratio (SMR). Am. J. Epidemiol. 131, 373–375 (1990).

92. A. J. Dobson, K. Kuulasmaa, E. Eberle, J. Scherer, Confidence intervals for weighted sums of Poisson parameters. Stat. Med. 10, 457–462 (1991).

93. N. R. Garud, P. W. Messer, E. O. Buzbas, D. A. Petrov, Recent selective sweeps in North American *Drosophila melanogaster* show signatures of soft sweeps. PLoS Genet. 11, e1005004 (2015).

94. L. Speidel, M. Forest, S. Shi, S. R. Myers, A method for genome-wide genealogy estimation for thousands of samples. Nat. Genet. 51, 1321–1329 (2019).

95. J. Terhorst, J. A. Kamm, Y. S. Song, Robust and scalable inference of population history from hundreds of unphased whole genomes. Nat. Genet. 49, 303–309 (2017).

96. X. Zhou, M. Stephens, Genome-wide efficient mixed-model analysis for association studies. Nat. Genet. 44, 821–824 (2012).

97. A. Rahman, I. Hallgrímsdóttir, M. Eisen, L. Pachter, Association mapping from sequencing reads using k-mers. Elife. 7 (2018), doi:10.7554/eLife.32920.

98. S. D. Jackman, B. P. Vandervalk, H. Mohamadi, J. Chu, S. Yeo, S. A. Hammond, G. Jahesh, H. Khan, L. Coombe, R. L. Warren, I. Birol, ABySS 2.0: resource-efficient assembly of large genomes using a Bloom filter. Genome Res. 27, 768–777 (2017).

99. S. Picelli, A. K. Björklund, B. Reinius, S. Sagasser, G. Winberg, R. Sandberg, Tn5 transposase and tagmentation procedures for massively scaled sequencing projects. Genome Res. 24, 2033–2040 (2014).

100. R. Corbett-Detig, R. Nielsen, A hidden markov model approach for simultaneously estimating local ancestry and admixture time using next generation sequence data in samples of arbitrary ploidy. PLoS Genet. 13, e1006529 (2017).

101. K. W. Broman, H. Wu, S. Sen, G. A. Churchill, R/qtl: QTL mapping in experimental crosses. Bioinformatics. 19, 889–890 (2003).

102. D. T. Hoang, O. Chernomor, A. von Haeseler, B. Q. Minh, L. S. Vinh, UFBoot2: improving the ultrafast bootstrap approximation. Mol. Biol. Evol. 35, 518–522 (2018).

103. A. Dobin, C. A. Davis, F. Schlesinger, J. Drenkow, C. Zaleski, S. Jha, P. Batut, M. Chaisson, T. R. Gingeras, STAR: ultrafast universal RNA-seq aligner. Bioinformatics. 29, 15–21 (2013).

104. B. Li, V. Ruotti, R. M. Stewart, J. A. Thomson, C. N. Dewey, RNA-Seq gene expression estimation with read mapping uncertainty. Bioinformatics. 26, 493–500 (2010).

105. M. I. Love, W. Huber, S. Anders, Moderated estimation of fold change and dispersion for RNA-seq data with DESeq2. Genome Biol. 15, 550 (2014).

106. L. D. Thomas, S. Bandara, V. M. Parmar, R. Srinivasagan, N. Khadka, M. Golczak, P. D. Kiser, J. von Lintig, The human mitochondrial enzyme BCO2 exhibits catalytic activity toward carotenoids and apocarotenoids. J. Biol. Chem. 295, 15553–15565 (2020).

107. J. von Lintig, K. Vogt, Filling the gap in vitamin A research. Molecular identification of an enzyme cleaving beta-carotene to retinal. J. Biol. Chem. 275, 11915–11920 (2000).

108. M. E. Kelly, S. Ramkumar, W. Sun, C. Colon Ortiz, P. D. Kiser, M. Golczak, J. von Lintig, The biochemical basis of Vitamin A production from the asymmetric carotenoid β-cryptoxanthin. ACS Chem. Biol. 13, 2121–2129 (2018).

109. A. Daruwalla, J. Zhang, H. J. Lee, N. Khadka, E. R. Farquhar, W. Shi, J. von Lintig, P. D. Kiser, Structural basis for carotenoid cleavage by an archaeal carotenoid dioxygenase. Proc. Natl. Acad. Sci. U. S. A. 117, 19914–19925 (2020).

