## Supplementary Materials for "Genomic consequences of domestication of the Siamese fighting fish"

**This PDF file includes:**

Materials and Methods
Supplementary Text
Figs. S1 to S18
Tables S1 to S4

### Materials and Methods

Sample collection and re-sequencing

Sequencing libraries were generated from genomic DNA (gDNA) extracted from ethanol-preserved fin clips of wild *Betta splendens* species complex and *Betta compuncta* samples and were sequenced on an Illumina platform, following the Illumina's PCR-based library protocols. For resequencing of ornamental betta, gDNA was extracted from fin clips using the Mag-Bind® Blood & Tissue DNA HDQ Kit (Cat. M6399, Omega Bio-Tek). Whole genome sequencing libraries for the ornamental samples were generated and indexed using QIASeq FX DNA Library Kit (Cat. 180473, Qiagen) with an average insert size of 550 base pairs (bp). All libraries were PCR-free with the exception of Orn30, Orn27, Orn37 which were PCR-amplified for 6 cycles. Through receiver operating characteristic (ROC) curves of total variants called and number of Mendelian errors in the trios and quartet, we determined that 15× coverage was enough for our sample set, and sequenced our ornamental splendens panel to ~15×; a trio and quartet were sequenced to >30× coverage. We demultiplexed using bcl2fastq allowing for 0 mismatches and trimmed our reads using trimmomatic version 0.36 (*63*).

For RNA sequencing for genome annotation, organs (fin, gonad, brain, liver and gill) from a wild *B. splendens* adult male from Chiang Mai and a wild *B. splendens* adult female from Phetchaburi were flash frozen and stored in RNAlater ICE (Cat. AM7030, Invitrogen) for transport until extraction. They were mechanically disrupted in TRIzol Reagent (Cat. 15596026, Invitrogen) using a homogenizer and RNA purified using Zymo Direct-zol RNA MiniPrep Plus (Cat. R2070, Zymo Research), which includes a DNAse step to remove DNA contamination. RNA sequencing libraries were generated using NEBNext Ultra Directional RNA Library Prep Kit for Illumina (Cat. E7420, New England Biolabs (NEB)). Libraries of two different insert sizes (~350 bp and ~515 bp) were made for each tissue and sequenced to an average coverage of $72\times10^6$ reads (range: $33\times10^6$-$145\times10^6$) with 2x150 bp reads in a NextSeq 500.

RNAseq reads were submitted to NCBI Sequence Read Archive as BioProject PRJNA486171 and used by NCBI to annotate the *B. splendens* genome in Annotation Release 100.

Samples and the associated metadata included in the study are listed in Supplementary Table 1.

<u>Sampling sites and distribution</u>

Approximate distribution ranges are based on IUCN red list, except for *B. siamorientalis*, where an approximate range was drawn from ref. (*64*). The range for *B. splendens* was extended to the north based on records in ref. (*65*) and our own sampling locations.

<u>Genome assembly and info</u>

The fBetSpl5 assembly was created by assembling 48× PacBio CLR reads from SplBan0 with Falcon-unzip (*66*). Haplotigs representing retained duplication were removed using purge_haplotigs before scaffolding the assembly using Scaff10x (https://github.com/wtsi-hpag/Scaff10X) with 183× Illumina HiSeqX, 10X Chromium reads from a different individual (SplPhe0). Following scaffolding the assembly was gap-filled with PBJelly (*67*) and polished with Arrow (https://github.com/PacificBiosciences/pbbioconda) using the long reads data, and then polished further using Freebayes (*68*) with 83× Illumina HiSeqX reads from SplBan0.

After assembly the genome was improved with manual curation, a gEVAL database was used to correct mis-scaffolding along with aligned Bionano consensus optical maps generated from SplPhe0. Additional scaffolding was possible utilizing haplotypic contig overlaps. The assembly was separated into chromosomes through alignment with the existing *Betta splendens* assembly (GCA_003650155.1) and named by synteny to the medaka (*Oryzias latipes*) assembly (GCA_002234675.1). Curation resulted in 67 manual breaks, 92 manual joins and the removal of 113 regions representing false duplications. The final assembly consists of 69 scaffolds totaling

441Mb, with a scaffold N50 of 20.1Mb. 98.62% of the assembly has been assigned to the 21 chromosomes.

We used the genomes of fAnaTes1.2 and fBetSpl5.3 to generate de novo TE consensus libraries using RepeatModeler2 (*69*), which uses the RepeatScout (*70*) and RECON (*71*) algorithms for discovery of transposable elements. To enhance detection of LTR retrotransposons we used LTR_retriever (*72*) and LTR_harvest (*73*) options on RepeatModeler2. This combined new library of de novo elements was used to annotate the genomes of fBetSpl5.3, Bspl.v1.2018.1, and fAnaTes1.2, with RepeatMasker version open-4.0.9 (*74*). For further analysis, TE families were parsed from RepeatMasker output file (Suppl. Figs. 1B,C).

<u>Animal husbandry</u>

Adult fish were housed individually in 1-1.4 L tanks in a system with recirculating water with automated dosing using Instant Ocean Sea Salt and sodium bicarbonate to maintain a conductivity of 1.0 µS and a pH of 7.0. 10% of the volume of the system is replaced daily with reverse osmosis purified water. Water and room temperature are maintained at 28 ºC with 14:10 hours light:dark cycle.

Larvae were grown from 5-15 days post fertilization (dpf) in 2.8 L tanks without recirculation at a density of 30 larvae per tank, and were fed *Brachionus rotundiformis* rotifers from 5-14 dpf. At 15 dpf, fry were moved to 37 L grow-out tanks containing biofilters and fed 2-3 times daily with decapsulated newly-hatched *Artemia nauplii*. Water quality in grow-out tanks was monitored regularly for presence of nitrates, nitrites, and ammonia and water exchanged with system water as needed. Animal experimentation protocols were approved by the Columbia University Institutional Animal Care and Use Committee.

Comparisons across reference assemblies

For synteny analyses between genome references, we used nucmer 4.0.0beta2 (*75*) on the following reference assemblies: male wild *B. splendens*: GCF_900634795.3 (fBetSpl5.3); female ornamental *B. splendens* based on Oxford Nanopore (*dmrt1_XX*): GCA_013403625.1, Bspl.v1.2018 (*12*); male ornamental *B. splendens* based on HiC (*dmrt1_XY*): GCA_003650155.1, ASM365015v1 (*11*);  ornamental *B. splendens* based on Illumina short reads: Betta_splendens_chrs.v1 (*13*). We used mplotter (*76*) with '-filter' to generate dotplots of the nucmer output to compare the reference.

Since sex chromosomes often involve chromosomal rearrangements, we searched for evidence of such rearrangements involving *dmrt1* in males and in females. The long-molecule Oxford Nanopore reads from the ornamental male (*12*) did not contain any "chimeric" reads that would indicate an inversion.

Alignment, variant detection, filtering, and phasing

We aligned all samples to the fBetSpl5.3 NCBI genome using bwa-mem2 (*77*) (https://github.com/bwa-mem2/bwa-mem2) and marked duplicate reads using picard version 2.0.1 (http://broadinstitute.github.io/picard/) (*78*).

We identified filter criteria for variants based on Mendelian errors from a trio and quartet, using variant calls across the ornamental samples using two variant callers: BCFtools mpileup version 1.11 (*79*)  and GATK4 version 4.1.4 (*80*). We assumed that de novo mutations are rare, and that Mendelian errors are likely sequencing artifacts, and thus represent false positives. Based on ROC analysis, BCFtools appeared to call more variants at similar false positive rates compared to GATK. We determined that filtering out variants with Mapping Quality (MQ) < 50, and genotypes with Genotype Quality (GQ) < 30, Depth (DP) < 4, and where the allele depth (AD) support for genotype was less than 2 reads provided the least number of Mendelian errors with the highest number of non-Mendelian errors through ROC of mpileup variant calls. We also excluded variants that are adjacent to each other within 3 bp. To increase confidence of the heterozygous genotypes, we further filtered out heterozygous genotypes in which the variant allele fraction (alt depth/total depth, VAF) showed an allelic imbalance: VAF<0.25 and VAF>0.75.

For short (<100 bp) indel analysis, we used GATK4 for calling indels as it uses local realignment around indels and filtered with more stringent criteria including requiring indels to have GQ>50, fraction missing <0.2, and are present in more than 1 individual. These variants must also be concordant with Mendelian expectation if present in either the trio or quartet, leaving 228,066 indels. For SV analysis (structural rearrangements, indels > 100 bp), we used smoove (https://github.com/brentp/smoove), filtering out genotypes with GQ<50 and where deletions carried a depth more than 0.7 relative to the flanking region depth (DHFFC<0.7). We additionally filtered out duplications that were less than 1.3× fold-change relative to other similar GC-content bins across the genome (DHBFC>1.3), and inversions/translocations with DP<4. We performed chi-square analysis corrected for false discovery rate to assess association of variants near regions identified by genome-wide association and quantitative trait loci analysis. We manually inspected these variants through Integrated Genomics Viewer (IGV) (*81*).

*Region filter*

We calculated read counts for each bam file across the fBetSpl5.3 which was segmented into 1000 bp windows with a 500 bp slide. For each sample, we $\log_2$-normalized the read counts of each window by the median read count across the genome. For pop-genome analysis, we removed any region where the read counts were either greater than 2.5 standard deviations (SD) from the median within a sample, or where the variance of the log-normalized read counts across the bam files was greater than 2.5 SD from the median variance across the genome across all bam files. For association mapping analyses in ornamental betta, we applied the latter filter only to the ornamental betta samples to prevent removal of regions that were lost or gained in other species/populations, but are stably diploid in ornamental betta.

To further filter out regions with poor mappability, we used SNPable (https://github.com/lh3/misc) with –l 200 and –r 0.5, and converted the masked genome to a bed file using a custom python script. Although initially designed for single-end read mappability estimations, we observed that 98.0% of regions that were masked through SNPable were also removed based on our read-coverage filter.

| | **Filters** | **Genome (bp) removed (Chromosomes only)** | **% Genome removed (Chromosomes only: 435314815bp)** |
|---|---|---|---|
| *B. splendens* species complex | SNPable filter | 31564934 | 7.3% |
| | Transposable Element (TE) filter | 89388187 | 20.5% |
| | Read-coverage filter | 99305522 | 22.8% |
| | Read-coverage + SNPable filter + TE filter | 127909927 | 29.4%; 98% overlap of SNPable to read-coverage; 74.1% overlap of TE filter to read-coverage |
| *B. splendens* | Read-coverage + SNPable filter + TE filter | 122306715 | 28.1%; 93.5% overlap of SNPable to read-coverage; 47.6% overlap of TE filter to read-coverage |
| Ornamental betta | Read-coverage + SNPable filter + TE filter | 120969387 | 27.8%; 86.3% overlap of SNPable to read-coverage; 39.1% overlap of TE filter to read-coverage |
| Wild *B. splendens* | Read-coverage + SNPable filter | 120059452 | 27.5%; 91.6% overlap of SNPable to read-coverage; 44.6% overlap of TE filter to read-coverage |

*Phasing*

Using the filters described above, along with a fraction missing threshold of 0.2, we phased the genotypes of the remaining variants in a two-step method. We generated initial phase-sets of our quartet and trio using WhatsHap version 0.18 (*82*), which performs pedigree-informed, read-backed haplotype assembly. We included the phase-set of the families generated by WhatsHap into ShapeIt4 version 4.1.3 (*83*) to phase all samples with the recommended error rate (--use-PS 0.0001) and settings for sequencing data (--sequencing). To increase phasing accuracy, we increased the number of mcmc iterations "--mcmc-iterations 10b,1p,1b,1p,1b,1p,1b,1p,10m" and the number of conditioning haplotypes "--pbwt-depth 8".

Copy number calling and segmentation

We calculated and $log_2$-normalized read counts ($log_2R$) in 1000 bp intervals with a 500 bp slide across the nuclear genome. We called copy number (CN) states (CN0-CN8) implementing a Poisson logit-linked model (*84*). To assess more focal duplications such as gene-level duplications, we calculated the average read depth across each gene normalized to the length of the gene taken from the fBetSpl5.3 NCBI genome annotation.

Species phylogenies

Based on the filtered phased biallelic SNP callset, we calculated pairwise differences across individuals using plink v1.90p (*85*) with the option --distance square both for the whole genome and in 100 kb windows. NJ trees were calculated with the nj function of Python's skbio package version 0.5.6 (http://scikit-bio.org/). Block bootstrap support was determined by randomly subsampling window-pairwise differences with replacement and testing how often a node of the whole genome tree was supported in 1000 random replicates. Maximum likelihood trees of the biallelic SNP call set were generated using IQTree v2.0.3 (*86*, *87*) with recommended model based on ModelFinder and then bootstrapped (-B).

Principal component analysis

We used plink v1.90p (*85*) with the options --double-id --allow-extra-chr to convert the filtered phased biallelic SNP vcf file to binary plink format. We used this file for PCA analyses in plink (option --pca) on (i) all samples except the outgroup individual Com0 and (ii) only *B. splendens* and ornamental betta samples in each case using a minor allele frequency cutoff of 1% (--maf 0.01). The included samples were specified with the option --keep.

ABBA-BABA analysis

We computed ABBA-BABA tests for all triplets of samples with Dsuite Version 0.4 r38 (*88*) defining each individual as a separate population and running Dsuite Dtrios with default parameters. Unless otherwise stated, we used the *B. compuncta* sample as outgroup. All shown f4 ratio plots show f4(p1,p2,p3,outgroup) for f4(p1,p2,p3,outgroup) >0 and -f4(p2,p1,p3,outgroup) otherwise.

$f_{DM}$ analysis in ornamental betta

Windowed $f_{dM}$ was computed across the genome with Dsuite Dinvestigate using a window size of 100 SNPs with a step of 25 SNPs for each of the triplets of samples that showed signs of introgression with the ABBA-BABA tests. For each triplets of samples, we smoothed the $f_{DM}$ across the chromosome using a MAD winsorization as described by ref. (*89*), so that a single window outlier inconsistent with the $f_{DM}$ values observed in adjacent windows would not affect downstream segmentation. We then applied a piecewise segmentation across the chromosomes, and calculated the mean $f_{DM}$ of each segment. Across all triplets of samples tested, we observed a clear bi-modal distribution in $f_{DM}$ values of these segments with one distribution centered at 0, and another that was higher than $f_{DM}$ of 0.2. We thus filtered for segments where $f_{DM}$ greater than 0.2, and merged adjacent remaining segments that were within 5 kbp of each other to estimate the start and end points of the introgressed regions. Local gene trees across these genomic regions were then constructed using IQ-Tree across the haplotypes of the ornamental bettas and other *Betta splendens* complex samples that did not show recent introgression through the ABBA-BABA tests. Across each region, we determined the ancestry of each haplotype for the ornamental sample (p2) with high $f_{DM}$ by calculating the average branch length distance of the samples haplotype to the haplotypes of other samples for each species. We also calculated the average distance of each *B. splendens* haplotype to other *B. splendens* haplotypes and calculated the standard deviation. If the average distance of the p2 sample's haplotype to the haplotypes of non-*splendens* species was less than one standard deviation away from the average distance between *B. splendens* haplotypes, we determined that the haplotype likely shared ancestry with that non-splendens species. We further validated ancestry assignments by visual inspection of the trees. Proportion of genome carrying introgression accounts for the diploid genome.

Estimation of the de novo mutation rate

To estimate the per generation mutation rate, we sequenced an ornamental trio and an ornamental quartet (two offspring to one set of parents) family at >30× coverage and 37 ornamentals to ~15× coverage. We used variants called with bcftools mpileup as described above and more stringent filtering criteria (similar to ref. (*17*)) to identify putative de novo mutations while reducing false positives. Across all ornamental samples, we excluded regions with poor mappability as described in the regions filter. We removed any non-single nucleotide variants, and filtered out variants within 3 bp of an indel. We further removed sites where the average depth of coverage (DP) across samples was below DP/1.65 or above DP×1.65 (*90*), MQ<50, MQ0F>0.15, or had a homozygous genotype for the non-reference allele. We excluded any site where any of the ornamentals that were not offspring of the trio or quartet were heterozygous. We further excluded any site where individuals of the trio or quartet had GQ<30 or DP<10. We also removed sites where either parent had at least one read supporting the non-reference allele or where the offspring had no read support of the alternate allele in either the forward or reverse strand. We also removed any sites where the heterozygous genotypes in the offspring had a VAF<0.25 or >0.75.

Genotypes that were heterozygous in the offspring but homozygous for the reference allele in the parents were considered putative de novo mutations. The offspring of the trio had 5 de novo mutations, while the offspring of the quartet had 2 and 0 de novo mutations. We visualized each of these putative mutations in IGV for potential errors in alignment such as inclusion in reads near an indel, or with suspiciously high or low coverage compared to neighboring regions. We then calculated the accessible genome to use as the denominator in the mutation rate calculation by counting the number of bases remaining after the filters above, which yielded 311.9, 310.5, and 311.3 Mb for each trio. We multiplied the accessible genome by two because the mutation could have occurred in the paternal or maternal line and obtained a de novo mutation rate of $3.75\times10^{-9}$ mutations per base pair per generation by dividing the 7 observed mutations by the total accessible genome. We estimated the 95% confidence interval using the exact method (*91*, *92*) which assumes the number of de novo mutations as a Poisson single value with a chi-square distribution (*17*).

<u>Diversity statistics</u>

We calculated genome-wide diversity statistics including average nucleotide diversity (pi), Tajima's D for ornamental and wild *B. splendens*, accounting for accessible sites after filtering using the *B. splendens* regions filter. We removed the following samples from the wild populations because they had signs of introgression: SplKan8, SplPhe2, SplBan0, and SplBan1 from the analysis. Nucleotide diversity was calculated per site using the python package scikit-allel (https://github.com/cggh/scikit-allel), and the sum is divided by the total accessible (invariant and variant) sites remaining after filtering. Confidence intervals were calculated based on jack-knife resampling of chromosomes: Ornamental pi=0.00113, CI=$\pm1.2\times10^{-10}$; Kanchanaburi pi = 0.00137, CI=$\pm6.9\times10^{-11}$. We calculated windowed nucleotide diversity and Tajima's D in 10 kbp windows with 100 bp slide across the genome (Suppl. Fig. 12). We removed windows where 3 kbp or less of the window was accessible.

Linkage disequilibrium analysis

Across both ornamental betta and wild *B. splendens*, we assessed linkage disequilibrium (LD) using plink version1.90 (*85*), calculating $r^2$ with a MAF of 0.2 and --ld-window-kb 999 across each chromosome. To assess LD decay, we generated bins that incrementally increased by 3 bp from 0 to 999 kbp and calculated the mean $r^2$ of SNPs where the distance between the SNPs fell within each bin. For ornamentals, decay at halfmax (0.46 $r^2$) was at 6097 bp; for Kanchanaburi, it was 0.44 $r^2$ at 257 bp. Calculations of interchromosomal $r^2$ served as the baseline: Ornamental, 0.038; Kanchanaburi, 0.063 using --r2 interchrom and --ld-window-r2 0 with a MAF of 0.2 and --ld-window-kb 999.

Selection scans

We performed two types of genome-wide scans to identify any signatures of selection that may have occurred during domestication. We performed H-scan (*19*) to assess run lengths of pairwise haplotype homozygosity across the genome. We additionally performed G12 and G2/G1 scans with 200 SNP windows (*93*) to assess haplotype frequency spectra across the genome and to identify hard and soft sweeps across populations. To determine whether signature differences between groups of fish were greater than expected by chance while addressing linkage, we clumped SNPs that were in linkage using plink --clump, keeping the highest signature value to represent each clump. Across 1000 permutations, we randomly sampled signature values across the genome equaling the number of clumps across both populations, and calculated the absolute differences, keeping only those in the 95th percentile. We then calculated the threshold where alpha = 0.05 on the permuted data, to serve as the genome-wide threshold of significance.

<u>Divergence time estimations using Relate on phased genomes</u>

We used Relate version 1.1.4 (*94*) to infer the demographic histories of the wild *B. splendens* populations and ornamental betta. A *B. compuncta* (Com0) sample served as the ancestral sequence and we created a genetic recombination map with a constant recombination rate. We ran Relate using our point estimate for the de novo mutation rate and the upper and lower confidence interval (CI) estimates: $3.75\times10^{-9}$ mutations per bp per generation [Lower CI: $9.05\times10^{-10}$, Upper CI: $9.39\times10^{-9}$]. We assumed a generation time of 6 months (*7*) and with a starting effective population size of 50,000. We excluded the following samples with inferred signs of recent introgression and that are the offspring of the quartet or trio: Orn29, Orn21, Orn5, Orn38, Orn39, SplPhe2, SplBan0, SplBan1. We ran the Relate EstimatePopulationSize algorithm for one iteration to avoid introducing biases generated from the assumption that our samples belong to one panmictic population.

<u>Effective size histories using SMC++ on unphased genomes</u>

We used smc++ v1.15.4 (*95*) to estimate the effective size histories of the ornamental and wild splendens populations using their unphased genomes. We excluded the following samples from the analysis due to signs of gene-flow or due to being the offspring of the trio or quartet: SplKan8, Orn5, Orn29, Orn38, and Orn39. We separately estimated the population size histories using the mask that includes regions filtered out by the filters implemented for analysis as described above. For each population, we used 12 individuals to serve as the distinguished lineages on the basis of not being ancestral outliers or low coverage, and combined these inputs to generate a composite likelihood for each population. We used the de novo mutation rate ($3.75\times10^{-9}$ mutations per bp) estimated from the trio and quartet, using default settings with piecewise estimation. We performed jack-knife resampling of chromosomes to assess variability in the population size curves.

Genome-wide association analysis

Genome-wide association analysis for sex, color, and fin morphology across ornamental betta was performed using Gemma version 0.98.1 (*96*). We removed variants with minor allele frequency (MAF)<0.05, fraction missing <0.2, and $r^2$>0.8. We performed genome-wide association using bi-allelic variants for sex across 37 ornamentals (females:17; males:20). For color, we performed genome-wide association across 34 ornamentals (red:17; blue:17). For veil, we performed genome-wide association across 34 ornamentals (veil:18; crown:16).

We considered haplotype blocks to represent the number of independent statistical tests. We calculated the number of haplotype blocks using plink --blocks with a max kbp window of 999 and MAF 0.05. We identified 53,844 haploblocks in ornamental betta. Thus, for a Bonferroni-corrected *P* value of 0.05, we set the significance threshold in GWAS as $-\log_{10}(0.05/53844) = 6.03$.

To determine whether there are regions highly associated with sex that were not captured by reference-based association analysis, we performed a k-mer association analysis using HAWK version 1.3 (*97*) for sex in ornamental betta. We assembled the k-mers into contigs using ABYSS version 2.0 (*98*) and mapped these sequences to the fBetSpl5.3 genome using bwa-mem. All significantly associated k-mers could be successfully mapped to the reference. We identify the positions of the k-mers within these contigs and determine their genomic positions, preserving the k-mers and their associated *P* values.

Quantitative Trait Locus analysis of sex

We performed an $F_2$ intercross between a male (*dmrt1_XY*) and a female (*dmrt1_XX*) ornamental betta. We crossed the $F_1$ (*dmrt1_XY*) males and $F_1$ (*dmrt1_XX*) females to generate 211 $F_2$

progeny. We sexed each F2 progeny based on the presence of an ovipositor and of ovaries or testis. We generated Tn5-tagmented whole genome sequencing libraries (*99*) with an average insert size of 400 bp and sequenced the samples to a depth of ~0.05× using NextSeq 500/550 High Output Kit v2.5 (75 Cycles). We generated a panel of SNPs where the two founders differed in their genotype. In addition to the filters described in “Alignment, variant detection, filtering, and phasing” section, we filtered out variants with read-depth greater than or less than 1.5 SD of the median depth on a per sample basis, and removed variants that were within 150 bp from each other (read length). We excluded SNPs that were private to a founder, and were not present across the ornamentals in the GWAS cohort. We also excluded variants with non-Mendelian segregation in our trio and quartet (i.e. likely genotyping errors). For the F2s, we counted the allele read depth for each SNP using alleleCount (https://github.com/cancerit/alleleCount) with base quality >20 and mapping quality >35.

We performed stringent pruning of the SNPs used for imputation. We excluded SNPs where fewer than 10% of individuals had read-coverage. On the filtered SNP panel, we performed imputation using AncestryHMM version 0.94 (*100*) for an F2 intercross design. We removed SNPs where the genotype likelihood was less than 90% across more than 50% of the samples. We additionally removed SNPs where the difference of the posterior probability of the SNP relative to the preceding SNP was less than 10%. Using the remaining 15,572 SNPs, we performed QTL analysis for sex using R/qtl (*101*). We performed non-parametric interval mapping and determined significance thresholds by 1,000 permutations of the data using scanone.

To investigate if a secondary sex determination system exists for males carrying the *dmrt1_XX* haplotype, we also performed an F2 intercross between a male (*dmrt1_XX*) and female (*dmrt1_XX*) ornamental splendens, resulting in 100 F2s. Following the same filtering criteria, we performed QTL analysis for sex on 8,561 SNPs.

To minimize biases through environmental effects, we raised the *dmrt1_XY* male × *dmrt1_XX* female and *dmrt1_XX* male × *dmrt1_XX* female crosses at the same time using the same housing,water, and temperature conditions.

<u>XY haplotype analysis</u>

To generate a haplotype tree of the *dmrt1* locus, we used the phased variants filtered based on the criteria of "Alignment, variant detection, filtering, and phasing" section with the *B. splendens* regions filter within the LD block of chromosome 9 spanning 28850248-28884157 bp. We included both invariant and variant sites within the region, but masked out the regions that were excluded for downstream analysis by our filters. We excluded the following samples due to signs of recent gene-flow, due to being the offspring of the quartet or trio, or due to being sexually immature: Orn29, Orn5, Orn38, Orn39, SplPhe1, SplPhe2, SplBan0, SplBan1, SplChi1, SplChi2, and SplKan6. To generate the initial maximum likelihood tree, we used IQTree v2.0.3 (*86*, *87*) with ModelFinder. We then re-ran IQTree with the recommended model 'HKY+F+R2' with 1000 bootstraps, optimizing the bootstrap tree with nearest-neighbor interchange (-bnni) (*102*). From the maximum likelihood tree, we then collapsed all branches with bootstrap support <80%. We used the collapsed bootstrap tree (-te) to infer ancestral sequence at each node and tip of the tree (--asr). We selected the mutations that supported the basal branches of the *dmrt1_X* and *dmrt1_Y* haplotypes for re-genotyping.

Genotyping

<u>Sequencing based genotyping</u>

To validate the sex GWAS results, we generated an independent panel unrelated to the GWAS sample set consisting of 161 unrelated ornamentals (females:83; males:78). We performed a local chi-square analysis of the regions [+/-1kbp] carrying the highest associated GWAS SNPs for sex in the ornamentals to select SNPs that segregated the most between sex and present in the Y branch (Suppl. Table 2.1). We designed primers flanking the SNPs. We amplified and Illumina indexed (1× 0.9× SPRI-cleaned) products using OneTaq polymerase, (Cat. No. M0482 (conditions detailed in Suppl. Table 3,4), and sequenced the indexed (2× 0.9× SPRI-cleaned) amplicons using NextSeq 500/550 High Output Kit v2.5 (75 Cycles) (Illumina).

We removed aligned reads which had a mapping quality less than 10 and then performed bcftools mpileup with -C 50. We removed genotypes with DP<50, GQ<30, and where the SNP VAF was between 0.2-0.3 or 0.7-0.8 from further analysis.

Restriction fragment length polymorphism (RFLP) based genotyping for sex

We developed a quick diagnostic RFLP-based assay for genotyping *dmrt1_XX* vs *dmrt1_XY*. We identified a SNP with a $r^2>0.8$ with the top 0.5% of highest associated GWAS SNPs for sex that overlapped the restriction enzyme site for MluCI and designed primers flanking the SNP and the restriction enzyme site (Suppl. Table 3). We PCR-amplified the region and over-night digested the amplicons with MluCI (Cat. R0538L, NEB) (Suppl. Table 4). The PCR fragments were then visualized on a 2.5% agarose gel (Suppl. Fig. 18). To confirm the accuracy of the RFLP assay, we performed capillary (Sanger) sequencing of the undigested PCR amplicons of 5 *dmrt1_XY* samples and 5 *dmrt1_XX* samples and there was perfect concordance.

Larvae RNA sequencing and allele-specific expression

We crossed a *dmrt1_XY* male to a *dmrt1_XX* female, genotyped by both RFLP and sequencing. We extracted genomic DNA and RNA of the fry at 4, 8, and 12 days post fertilization (dpf) using Zymo Microprep Kit. We generated RNAseq libraries for 3 *dmrt1_XY* and 3 *dmrt1_XX* 4-dpf larvae using NEBNext Ultra Directional II RNA Library Prep Kit for Illumina (Cat. E7760, NEB). We sequenced the RNASeq libraries using NextSeq 500/550 High Output Kit v2.5 (75

Cycles). We aligned the reads using STAR2 (*103*) and counted reads with RSEM (*104*). Analysis of differential expression was done using DeSeq2 (*105*) implemented in R.

For targeted allele-specific expression analysis, we designed primers that spanned between exon 4 and exon 5 of *dmrt1* that included Tn5-ME adaptors (B: 5'-GTCTCGTGGGCTCGGAGATGTGTATAAGAGACAG-3; A: 5'-TCGTCGGCAGCGTCAGATGTGTATAAGAGACAG-3'), and flanked a X-Y variant on exon 5 (chromosome_9: 28864161) (Suppl. Table 3). We generated cDNA using LunaScript RT (Cat. E3010, NEB), and amplified the target region using NEB Q5® High-Fidelity DNA Polymerase (Cat. M0491, NEB). We 0.9× SPRI-cleaned and Illumina-indexed the product. Post-indexing, we SPRI-cleaned (2× 0.9×) the product and sequenced using NextSeq 500/550 High Output Kit v2.5 (75 Cycles). We additionally performed sequence-based genotyping of heterozygous SNPs including this site across the 4, 8, and 12 dpf larvae (Suppl. Table 3). The number of alternate allele reads to total reads across the heterozygous sites fell within the variance of a normal binomial distribution for all sequence-based genotyping SNPs, suggesting that technical noise was minimal. We performed a binomial test to assess potential allele-specific expression (ASE) at each dpf across larvae, implemented in R.

<u>QTL analysis of color</u>

We generated an intercross between a red (*dmrt1_XY*) male and a blue (*dmrt1_XX*) female ornamental betta, crossed 2 pairs of $F_1$ hybrids, and generated 211 $F_2$ progeny. We genotyped $F_2$s as described in "Quantitative Trait Locus of sex". We then performed Haley-Knott regression in R/qtl with 17,155 markers.

<u>Imaging</u>

We photographed the $F_2$s after they reached sexual maturity, which was confirmed by the presence of ovipositors in females and bubble nests in the males. We further confirmed sexual maturation by presence of ovaries or testis after photos were taken. We used a Canon EOS RP camera with a Canon Macro lens EF 100 mm on manual focus set on a mini tripod. Settings for the camera were the following: ISO auto, F 4.0, exposure 0, and stabilizer activated. The fish was placed in a plastic tank (2.5" x 3.5" x 4.4") which showed minimal light distortions when assessed with polarized lens, and pictures were taken in a white table top photo studio tent (14” x 16” x 14”) lined with white LEDs on all sides of the booth. For our analysis, we observed minimal difference between the use of a halogen light source and LEDs after calibration with a color checker card, and all analysis was performed with the LEDs. At the start of each photo session, we calibrated the camera’s white balance using the white balance (CT24-23-1424) provided on the 24ColorCard from CameraTrax.com. Each image included the color checker to calibrate each image to the reference colors provided on the card. We took pictures of both sides and top of each fish.

Color Processing analysis

To calibrate each image to the reference color card, we imported each .RAW image using rawpy version 0.15.0 (https://pypi.org/project/rawpy/). Using a custom python script, we automatically identified the color-matrix, and calibrated each picture based on the reference color values on the color card from CameraTrax.com with a pipeline built on python packages, rawpy, colour, and PlantCV. We segmented the fish using a Recursive-Convolutional Neural Net model across 122 images that were annotated and trained in Matlab 2019b after adjusting contrast thresholds on masks of the image. Images of fish that were poorly segmented with the automated segmentation were manually segmented with ImageJ for downstream analysis. Both automated and manually segmented images were exported to a .tiff format, and downstream analysis of color were performed on the segmented images using a custom R script based on imager (https://dahtah.github.io/imager/). Color analysis was based on the Hue [0-360º]-Saturation[0-1]-

Value[0-1] (HSV) color space. We selected the hue range for red to ([0,0.045], [0.98,1]), and for blue ([0.6,0.728]), whose ranges maximized the difference between the red (n=17) and blue (n=17) ornamental betta through iterative subtractive binning. We similarly determined the range [0-0.3] to assess blackness based on the value axis along the color space.

Carotenoids extraction

The frozen fish skin samples were grinded with a mortar and a pestle precooled on dry ice. After weighing, the powder was transferred to an Eppendorf tube and was homogenized in 250 µL phosphate-buffered saline. Then, 250 µL of methanol was added and the suspension was vortexed. After addition of 500 µL acetone, the suspension was vortexed and kept for 5 minutes on ice. 200 µL of diethyl ether and 400 µL hexane were added to extract carotenoids. Phase separation was achieved by centrifugation at 3,000 x g for one minute. The upper organic layer was transferred to a new Eppendorf tube. The extraction was repeated once with 500 µL hexane. The combined organic phases were vacuum dried in a Speedvac (Eppendorf, Hauppauge, NY) and the debris were dissolved in 200 µL hexane: ethyl acetate (70:30 v/v) for High-performance liquid chromatography (HPLC) analysis.

HPLC analysis

HPLC performed on a 1200 Agilent HPLC series equipped with a diode array detector and normal-phase Agilent Zorbax silica column (4.6 mm ID x 150 mm with 5 µm packing; Agilent, Santa Clara, CA, USA) as previously described ref. (*106*). Chromatographic separation was achieved with isocratic flow of hexane: ethyl acetate (90:10 v/v) for β-carotene and retinoid separation and of hexane: ethyl acetate (70:30 v/v) for carotenoids from fish skin, respectively. The flow rate was 1.4 mL/min. The system was scaled with known amounts of authentic standards.

Identification of echinenone

We compared the retention time and spectral characteristics of beta-beta-carotene-4,4'-dione (canthaxanthin) with an unknown peak (X) in the fish extracts. The absorption maximum of peak X at 458 nm showed a bathochromic shift of ~9 nm when compared with the absorption maximum (467 nm) of canthaxanthin (Suppl. Fig. 16A). The addition of the second ketone group in canthaxanthin increases the carotenoid's polyene chain conjugation length, accompanied by a spectral shift towards the absorption of longer wavelength light. The absorption maximum agrees with the reported absorption maximum of echinenone in hexane as reported in the carotenoid database (http://carotenoiddb.jp). Further, peak X's retention time was shifted towards an earlier time point, being consistent with the loss of an oxo-group in one of the terminal β-ionone rings of canthaxanthin. Together with the characteristic shape of the absorption curve of peak X, this information strongly indicates that peak X is most likely echinenone.

Biochemical activity of β-carotene oxygenase I-like

We analyzed the enzymatic activity of the red-allele and blue-allele β-carotene oxygenase 1-like (BCO1L) protein by co-expressing the recombinant MBP-BCO1L fusion proteins in a β-carotene producing *E. coli* strain. This method has been previously used to characterize carotenoid cleavage enzymes from several species (*107*). The presence of β-carotene and various retinal-oxime diastereomers in the lipid extracts of these bacteria that express BCO1L display β-carotene 15-15'-dioxygenase activity (Supp. Fig. 16B). We confirmed enzymatic activity with protein crude extracts after lysing the bacteria (Supp. Fig. 16C). The red-allele BCO1L displayed activity in these assays (*108*). The blue-allele BCO1L did not display catalytic activity under the applied conditions. We enriched for the red-allele BCO1L by affinity chromatography using an established protocol for MBP-fused carotenoid cleavage dioxygenases(*106*). The purified protein maintained its catalytic activity and converted β-carotene into all-*trans*-retinal similar to the control (human BCO1 protein).

We modelled the structures of red and blue allele BCO1L proteins using the recently determined *Candidatus Nitrosotalea devanaterra* (*Nd*CCD) structure as template (*109*) (Suppl. Fig. 16G). The Thr>Ile mutation is located at the surface of the protein and far away from the enzymes' active site (non-polar substrate tunnel). Therefore, the mutation most likely does not affect the substrate binding and/or catalytic activity of the enzyme. Notably, the recombinant enzymes expressed in *E.coli*, particularly the blue allele of BCOL1, showed relatively poor solubility when expressed in *E coli* and the majority of the protein was found in the insoluble protein fraction (inclusion bodies). We cannot predict whether this behavior in *E. coli* reflects in the natural cellular environment of the proteins. However, our analyses suggest that the polymorphism affects protein solubility and stability rather than catalytic activity of the enzymes.

#### QTL analysis of fin morphology

We generated an $F_2$ intercross consisting of 139 $F_2$ progeny from a veiltail (*dmrt1_XX*) male with a crowntail (*dmrt1_XX*) female ornamental betta. We photographed the $F_2$s after they reached sexual maturity, which was confirmed by the presence of ovipositors in females and bubble nests in the males. We further confirmed sexual maturation by presence of ovaries or testis after photos were taken. We euthanized each fish, and pinned the fish to a sylgard-coated petri dish with the fins expanded. We photographed the fish in the lighting setup described in the "QTL analysis of color" Imaging section. Using Fiji, we annotated the length of the ray and webbings between the primary rays measured from the base of the tail scaled to centimeters using a ruler included within each image. We normalized the length of the webbing to the length of the ray. We genotyped $F_2$s as described in "Quantitative trait locus of sex". We performed Haley-Knott regression with 8,561 markers using R/qtl.

**Supplementary Text**

1. Synteny between wild *B. splendens* and ornamental betta

To discover structural rearrangements that may have arisen during domestication, we performed whole genome alignments using three ornamental betta references (*11–13*) and our wild *B. splendens* reference. An ornamental genome assembled using HiC is largely syntenic with wild *B. splendens (11)* (Suppl. Fig. 1D). The largest differences are a translocation of a portion of chromosome 14 in the wild genome to chromosome 12 in ornamental and a large intrachromosomal rearrangement of chromosome 16 (Suppl. Fig. 1D). The translocation of wild chromosome 14 was not observed in the comparison to the other two ornamental references, which were assembled using Oxford Nanopore sequencing (*12*) (Suppl. Fig. 1E) or Illumina short reads (*13*) (Suppl. Fig. 1F), suggesting this translocation is not present in all ornamental betta, or reflects an incorrect assembly of the ornamental HiC reference. In contrast, the rearrangement of chromosome 16 was also observed in the ornamental Nanopore reference and possibly in the Illumina reference (Suppl. Fig. 1E,F). Because this rearrangement is present in at least two separate ornamental betta, it may have occurred in the lineage that gave rise to these domesticated betta, or is specific to the wild *B. splendens* used for the fBetSpl5.3 reference or its lineage. To distinguish between these possibilities, we aligned the wild *B. splendens* genome to *Anabas testudineus (15)*, an anabantoid fish that is an outgroup to the *Betta* genus, and found that their chromosomes 16 were largely syntenic (Suppl. Fig. 1G). Thus, the most parsimonious explanation is that the chromosome 16 rearrangement occurred during domestication. The Nanopore assembly has many putative rearrangements compared to wild *B. splendens*, but these are also observed when aligning the Nanopore reference to the HiC ornamental reference, suggesting that the Nanopore assembly is not as well assembled. However, in addition to the large-scale chromosome 16 rearrangement, chromosomes 11 and 19 have small intrachromosomal rearrangements consistent between the three ornamental betta genomes and the wild *B. splendens*. Together, these results indicate that the genomes of ornamental and wild *B. splendens* are largely syntenic, except for a large rearrangement of a single chromosome and smaller changes in two other chromosomes.

<u>2. Evolutionary relationships between species of the *B. splendens* complex</u>

There are strong and highly significant signals of excess allele sharing among most species of the *B. splendens* species complex (Suppl. Fig. 3A). These signals are particularly striking for *B. mahachaiensis*, where excess allele sharing with other groups varies strongly among the three samples. For example, while Mah2 and Mah1 show very strong excess allele sharing with wild *B. splendens* samples compared to *B. siamorientalis* or *B. imbellis* (median f4-admixture ratios f4(*B. siamorientalis*, *B. splendens*; *B. mahachaiensis*, *B. compuncta*) = 28.5% and 19.8%, respectively; block jackknifing $P<10^{-300}$) (Suppl. Fig 3A), Mah0 shows the opposite pattern (median f4-admixture ratio f4(*B. splendens, B. siamorientalis*; *B. mahachaiensis*, *B. compuncta*) = 5.6%; block jackknifing $P <10^{-300}$) (Suppl. Fig. 3A). This pattern is highly suggestive of varying levels of introgression into these *B. mahachaiensis* samples.

For *B. imbellis*, sample Imb1 from the western Malay Peninsula was ~4.7% closer to *B. splendens* than was Imb0 from the eastern Malay Peninsula (median f4(Imb0, Imb1; *B. splendens, B. compuncta*)=0.047, block jackknifing $P<10^{-300}$) (Suppl. Fig. 3B). This difference between Imb0 and Imb1 was equally strong across all sampled *B. splendens* populations (Suppl. Fig. 3C), suggesting genetic introgression into *B. imbellis* populations of the western Malay Peninsula from a *B. splendens* lineage ancestral to, or separate from, the populations that we sampled.

3. Evolutionary relationships among *B. splendens* populations

Investigating the relationships between wild *B. splendens* populations, pairwise genetic differences, patterns of derived allele sharing (BBAA patterns from ABBA-BABA analysis) and phylogenetic clustering, give a consistent picture with the populations from Chiang Mai and Phetchaburi clearly closest, Kanchanaburi a sister group to them and Bang Phlat as an outgroup (Suppl. Figs. 2; 4A). Strong signals of excess allele sharing also suggest a complex divergence history. Most strikingly, while Kanchanaburi show moderate, mostly non-significant excess allele sharing with Bang Phlat  compared to two Phetchaburi samples, the pattern is reversed for the third Phetchaburi sample, SplPhe2, which shows highly significant excess allele sharing with Bang Phlat compared to Kanchanaburi (median f4(Kanchanaburi, SplPhe2, Bang Phlat, outgroup) = 4.5%, median $P$=2x10$^{-7}$, Suppl. Fig. 4B). This observation suggests significant amounts of genetic material from a population related to Bang Phlat in SplPhe2.

Furthermore, individuals from Kanchanaburi are significantly closer to Bang Phlat relative to individuals from Chiang Mai (median f4(Chiang Mai, Kanchanaburi; Bang Phlat, outgroup)= 1.9%, median $P$=0.003, Suppl. Fig. 4B). However, comparisons involving SplKan8 are outliers in this statistic, showing a much stronger signal (median f4(Chiang Mai, SplKan8; Bang Phlat, Outgroup)= 4.5%, median $P$=10$^{-8}$; Suppl. Fig. 4B).

Finally, individuals from Phetchaburi tend to be closer to individuals from Kanchanaburi compared to individuals from Chiang Mai (median f4(Chiang Mai, Phetchaburi; Kanchanaburi, outgroup)= 1.6%, median $P$=0.003), and individuals from Kanchanaburi are closer to Bangrad relative to Chiang Mai (Suppl. Fig. 4B). This trend tends to be more extreme for comparisons involving SplPhe2 (median f4(Chiang Mai, SplPhe2; Kanchanaburi, outgroup)= 2.2%, median $P$=0.0007) and clearly is most extreme for comparisons involving both SplPhe2 and SplKan8 (median f4(Chiang Mai, SplPhe2; Kanchanaburi, Outgroup)= 4.7%, median $P$=10$^{-9}$, Suppl. Fig.

4B). Together, the observations in this section suggest some shared ancestry between SplPhe2, SplKan8 and the two Bang Phlat samples. In Suppl. Note 4, we show that this pattern is explained by ornamental betta ancestry in these samples, likely due to introgression.

4. Excess allele sharing of ornamental betta with non-*splendens* species

We tested for excess allele sharing between ornamental betta and non-*splendens* with ABBA-BABA tests of the form *D*(wild *splendens*, ornamental; non-*splendens* species, outgroup). The outcome of these tests varied substantially by the population of origin of the wild *B. splendens* considered, pointing to a complex divergence history within *B. splendens*. This variation made these tests ill-suited to draw definite conclusions on non-*splendens* contributions to ornamental betta.

To address this issue, we focused on comparisons of the form *D*(ornamental except focal, focal ornamental; non-*splendens* species, outgroup) (Suppl. Figs. 4G-J). We removed *B. mahachaiensis* individuals Mah1 and Mah2 from the analysis as they carry ornamental betta introgression (Suppl. Fig. 4L), which could bias inferences about introgression into ornamental betta. The few genomic regions of excess allele sharing of ornamental betta with *B. siamorientalis* and *B. smaragdina* correspond to gene trees where a haplotype of the focal sample clusters with *B. imbellis* and *B. mahachaiensis,* respectively, suggesting that these regions are less phylogenetically distinct and suggest these signals also correspond to introgression from *B. imbellis* and *B. mahachaiensis*. Indeed, excess allele sharing with *B. siamorientalis* is expected under *B. imbellis* introgression, given their sister-species relationship, and *B. smaragdina* excess allele sharing in samples with *B. mahachaiensis* introgression can be explained by allele sharing between between *B. mahachaiensis* and *B. smaragdina* suggestive of gene flow between these species (Suppl. Fig. 3A).

To test whether the excess allele sharing of some ornamental betta with non-*splendens* species is also observed for wild *B. splendens*, we performed the same analysis as in Suppl. Figs. 4G-J for wild *B. splendens*. We computed for each focal wild *B. splendens* sample, *D*(same wild population as focal except focal, focal, non-*splendens* species, outgroup) and calculated the number of significant tests (i.e., below Bonferroni corrected $P<0.01$, corresponding to a z-score of 5.49) (Supp. Fig. 4K). We removed two wild splendens outlier individuals SplKan8 and SplPhe2 which carry ornamental betta ancestry (Suppl. Fig 5A). We found that wild *B. splendens* generally show no significant excess allele sharing with non-*splendens* species compared to other samples of the same population. The only clear exception to this are the two Bang Phlat samples. SplBan1 is consistently closer to other species than SplBan0 in all comparisons involving SplBan0 and SplBan1 as p1 and p2, respectively (Suppl. Figs. 5B,C). Since our reference assembly is based on SplBan0, we hypothesized that the observed pattern could be caused by the fact that sequencing reads for which the outgroup and the reference genome differ (corresponding to BXXA patterns) are less likely to align for relatively a divergent outgroup, leading to an excess of ABBA compared to BABA patterns. To test this we recomputed ABBA-BABA tests using the more closely related *B. smaragdina* as outgroup instead of *B. compuncta*. With *B. smaragdina* as an outgroup, there is no differential allele sharing of SplBan0 and SplBan1 with other groups (Suppl. Fig. 5C, right panel), while the excess allele sharing results of ornamental samples still hold up (Suppl. Figs. 4G-J). In conclusion, significant excess allele sharing with non-splendens species is generally not observed in wild *B. splendens* populations but it is common among ornamental betta, suggesting that many of them carry significant (but generally small amounts) of non-*splendens* introgression.

5. Ornamental betta introgression is widespread among wild *Betta*

One striking observation in the pairwise-distance based NJ trees presented above is that, despite high bootstrap support, the topology of relationships among wild *B. splendens* populations changes once ornamental samples are added (Suppl. Figs. 2A,C). In both trees, the samples from Chiang Mai and Phetchaburi are clearly closest to each other. However, without the ornamental samples, these populations form a sister-group to samples from Kanchanaburi with samples from Bang Phlat as a further outgroup. Conversely, in the tree including ornamentals, Chiang Mai and Phetchaburi cluster with ornamentals and Kanchanaburi and Bang Phlat together form a sister group to this. Furthermore, in the tree including ornamentals, one Phetchaburi sample changes its placement, now forming an outgroup to Chiang Mai and other Phetchaburi samples.

To further investigate this, we computed all ABBA-BABA statistics consistent with the phylogeny (including ornamentals) (Suppl. Figs. 2C;7). This analysis revealed that the above-mentioned Phetchaburi sample, SplPhe2, carries a substantial amount of ornamental betta ancestry (median f4(Chiang Mai, SplPhe2; ornamental, outgroup) ~22%, (Suppl. Fig. 5A). The two other Phetchaburi samples also have highly significant, but much weaker excess allele sharing with ornamental betta (median f4(Chiang Mai, other Phetchaburi; ornamental, outgroup) ~4%), suggesting that these samples might also have some ornamental betta introgression. Investigating the patterns of excess allele sharing along the chromosomes using $f_{dM}$ revealed that these patterns are widely distributed across chromosomes, without any apparent large outlier regions (Suppl. Fig. 6A). This suggests that ornamental introgression into the Phetchaburi population is not recent, leaving time for recombination to break down haplotypes.

Furthermore, Bang Phlat individuals are significantly closer to ornamental betta relative to Kanchanaburi samples (median f4 ratio = 4.0%, median $P=10^{-4}$; Suppl. Figs. 5A,B), consistent with genetic exchange between Bang Phlat individuals and a lineage related to our ornamental betta samples. Because Bang Phlat is located within Bangkok, ornamental introgression into the Bang Phlat population is plausible. An exception to the above pattern are comparisons involving SplKan8, which is closer to ornamentals compared to Bang Phlat (median f4 ratio = 3.2%, median $P$=0.004) and compared to other Kanchanaburi samples (median f4 ratio = 7.0%, median $P=10^{-12}$). The distribution of this signal along the genome is noisy, but on several large genomic regions of several 100 kilobases can be identified, pointing to more recent introgression (Suppl. Fig. 6A). Thus, in three of the four wild *B. splendens* populations sampled, we observe evidence of introgression from ornamental betta. The relatively short introgressed segments suggest this introgression happened many generations ago.

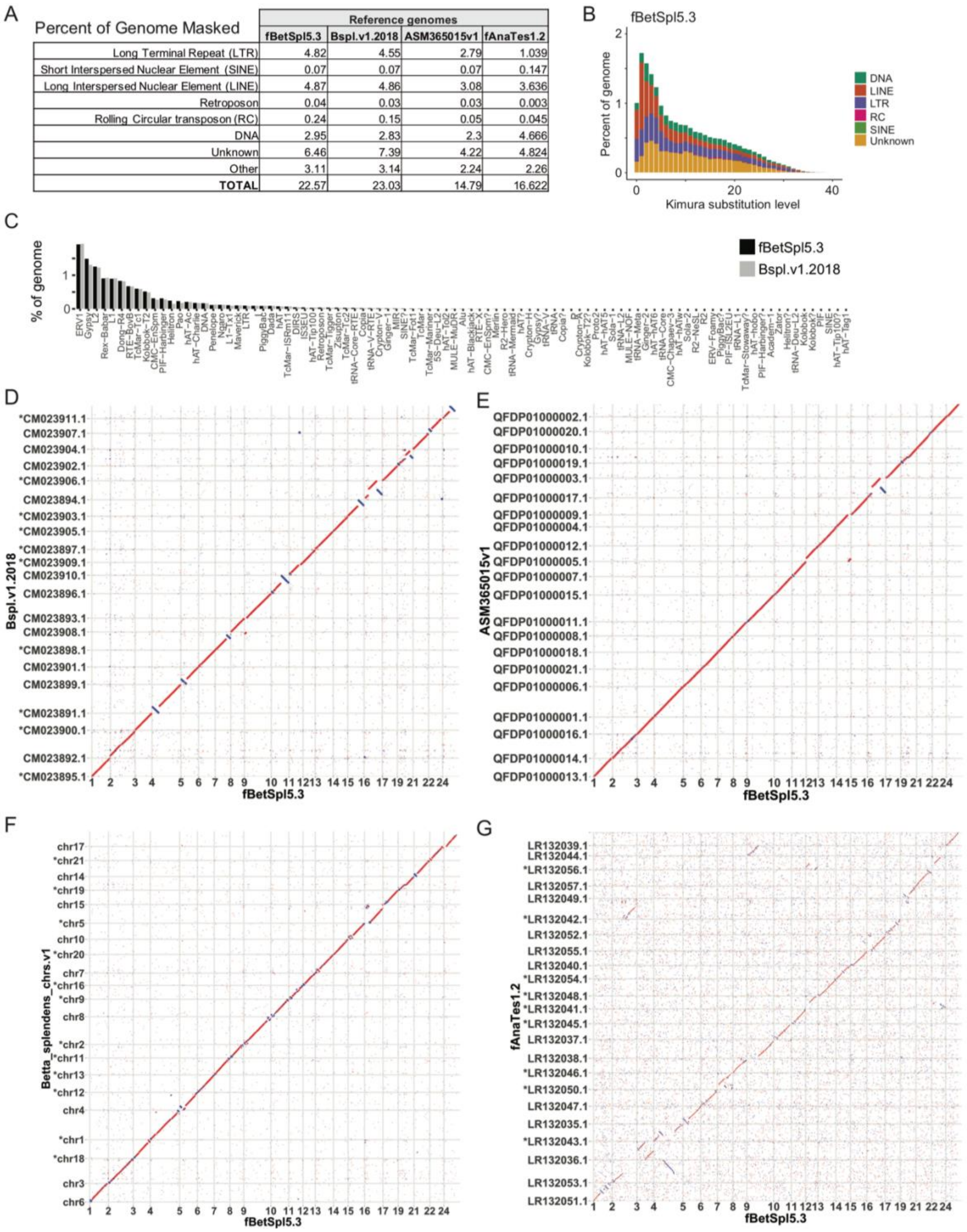


| Percent of Genome Masked | Reference genomes | | | |
|---|---|---|---|---|
| | fBetSpl5.3 | Bspl.v1.2018 | ASM365015v1 | fAnaTes1.2 |
| Long Terminal Repeat (LTR) | 4.82 | 4.55 | 2.79 | 1.039 |
| Short Interspersed Nuclear Element (SINE) | 0.07 | 0.07 | 0.07 | 0.147 |
| Long Interspersed Nuclear Element (LINE) | 4.87 | 4.86 | 3.08 | 3.636 |
| Retroposon | 0.04 | 0.03 | 0.03 | 0.003 |
| Rolling Circular transposon (RC) | 0.24 | 0.15 | 0.05 | 0.045 |
| DNA | 2.95 | 2.83 | 2.3 | 4.666 |
| Unknown | 6.46 | 7.39 | 4.22 | 4.824 |
| Other | 3.11 | 3.14 | 2.24 | 2.26 |
| TOTAL | 22.57 | 23.03 | 14.79 | 16.622 |

**Fig. S1. Cross reference genome comparisons. (A)** Genome masked across wild and domestic splendens reference genomes: fBetSpl5.3 (wild *B. splendens*; this paper); Bspl.v1.2018, Oxford nanopore read assembly polished with ASM365015v1 (ref.(*12*)). **(B)** Kimura-based sequence divergence across transposable element families in fBetSpl5.3. **(C)** Annotation of transposons across fBetSpl5.3 and Bspl.v1.2018. **(D)** Synteny plot of fBetSpl5.3 vs. ASM365015v1 reference genomes. **(E)** Synteny plot of fBetSpl5.3 vs. Bspl.v1.2018 (ref.(*11*)) reference genomes. **(F)** Synteny plot of fBetSpl5.3 vs. Betta_splendens_chrs.v1 (ref.(*13*)) reference genomes. **(G)** Synteny plot of fBetSpl5.3 vs. fAnaTes1.2 (ref.(*15*)) reference genomes. **(D-G)** Unplaced contigs were removed in the synteny plots. Asterisks indicate chromosomes with an opposite orientation from fBetSpl5.3.

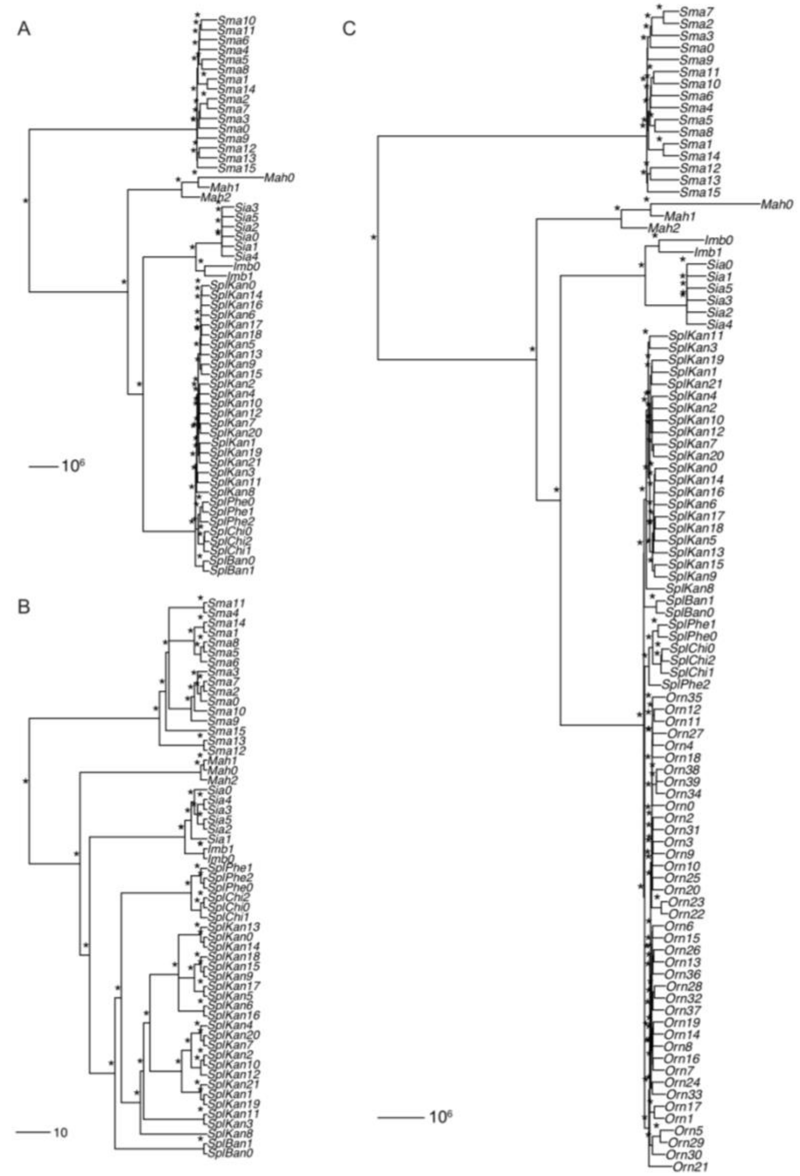


**Fig. S2.** ***Betta splendens*** **species-complex phylogeny. (A)** Neighbor-joining phylogeny based on whole-genome pairwise genetic differences of biallelic SNPs across species of the *Betta splendens* complex. Asterisks symbolize block bootstrap support values of greater than 990/1000 (99%). Block bootstrap is based on pairwise differences computed across 4361 windows of 100kbp. The tree was rooted with the outgroup *B. compuncta*. **(B)** Maximum-likelihood phylogeny based on variant SNPs across the genome excluding ornamental betta. **(C)** Neighbor-joining phylogeny based on bi-allelic SNPs across species of the *Betta splendens* complex including ornamental betta. Scale indicates differences in bp **(A,C)** and percent **(B)**.

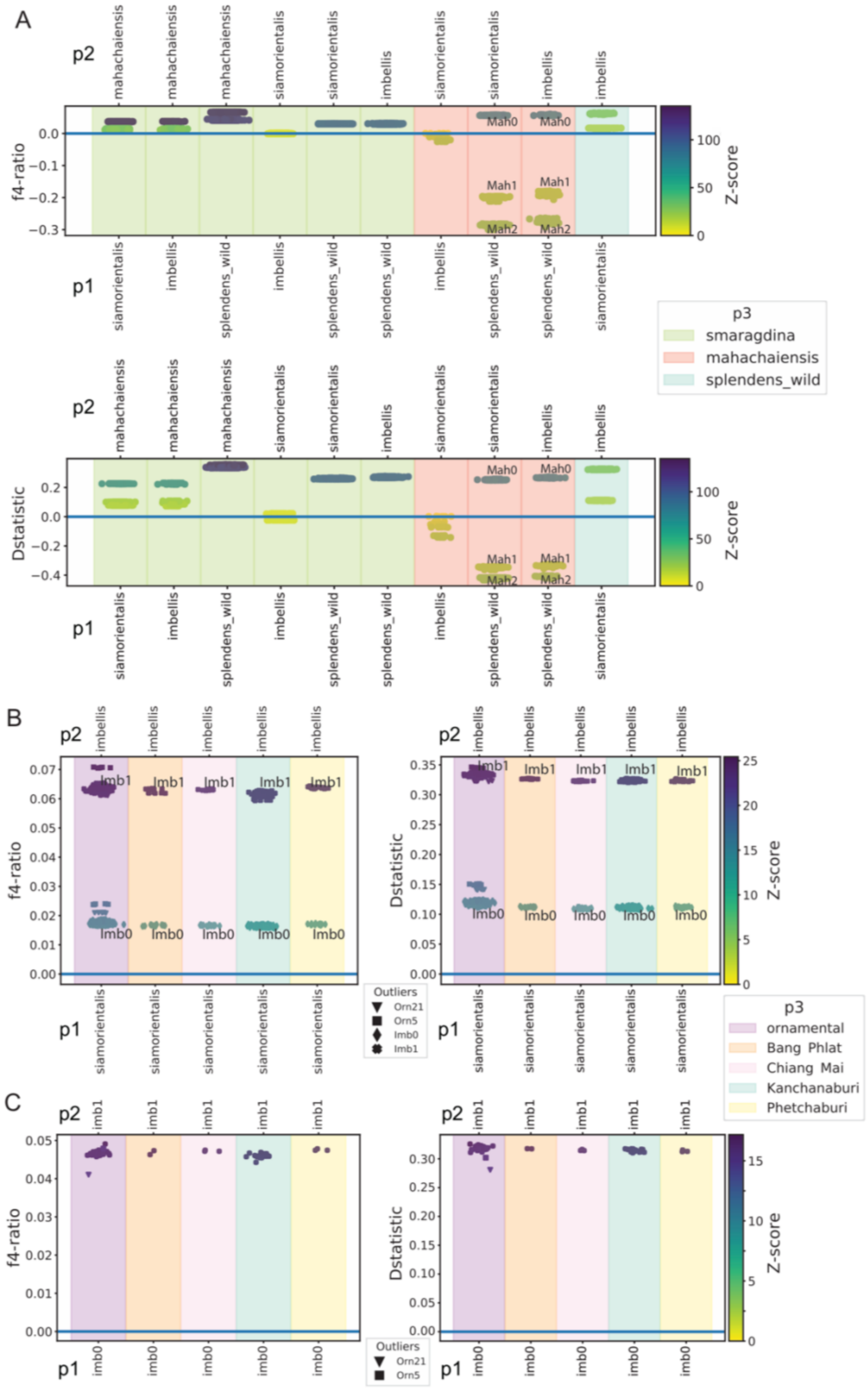


**Fig. S3. Excess allele sharing across *B. splendens* species complex and ornamental betta. (A)** Individual f4 admixture ratio tests and Patterson's *D* tests between *Betta splendens* species complex. p1, p2, p3 individuals are taken from different wild *Betta* species so that ((p1, p2), p3)

is consistent with the genome-wide phylogeny in Suppl. Fig. 2A. All f4 admixture ratio and *D* tests follow the order, f4 or *D* (p1, p2, p3, *B. compuncta*) (A-C). Color of dots indicate block-jackknife significance as Z-score; panel background colors indicate p3 (A-C). **(B)** Excess allele sharing among *B. siamorientalis* and *imbellis* with each of the wild *B. splendens* populations and with ornamental betta. f4 admixture ratio and *D* tests with the order: p1=*B. siamorientalis,* p2=*B. imbellis,* and p3=*B. splendens* population (Bang Phlat, Chiang Mai, Kanchanaburi, Phetchaburi) or ornamental betta. **(C)** Excess allele sharing among the two *B. imbellis* samples with wild *B. splendens* and ornamental betta. f4 ratio and *D* tests with the order: p1=Imb0 (*B. imbellis* individual), p2=Imb1 (*B. imbellis* individual), and p3=*B. splendens* population (Bang Phlat, Chiang Mai, Kanchanaburi, Phetchaburi) or ornamental betta.

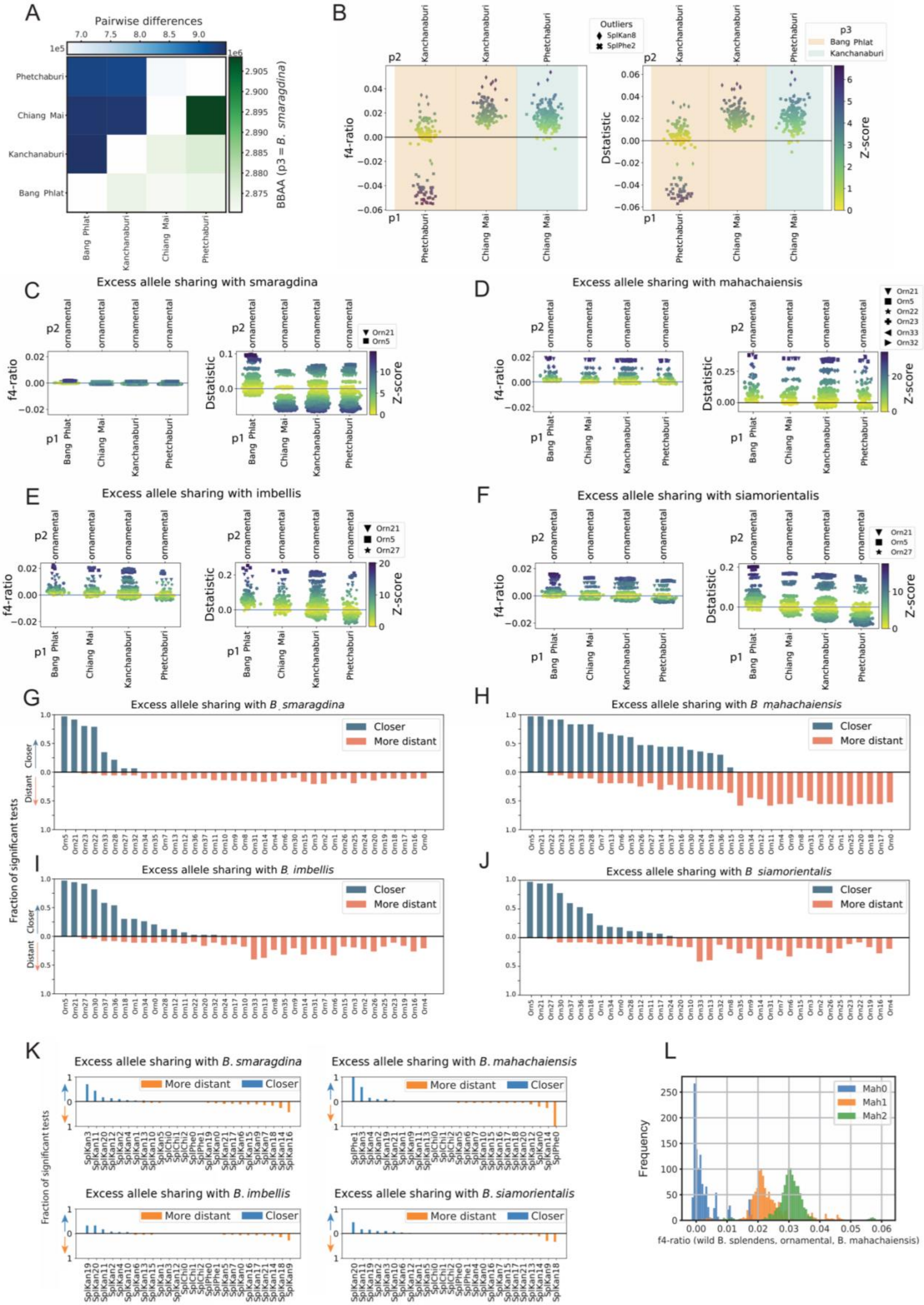
A
Pairwise differences
BBAA (p3 = B. smaragdina)
Phetchaburi
Chiang Mai
Kanchanaburi
Bang Phlat
B
Outliers
SplKan8
SplPhe2
p3
Bang Phlat
Kanchanaburi
f4-ratio
Dstatistic
Z-score
C
Excess allele sharing with smaragdina
D
Excess allele sharing with mahachaiensis
E
Excess allele sharing with imbellis
F
Excess allele sharing with siamorientalis
G
Excess allele sharing with B. smaragdina
H
Excess allele sharing with B. mahachaiensis
I
Excess allele sharing with B. imbellis
J
Excess allele sharing with B. siamorientalis
Closer
More distant
Fraction of significant tests
K
Excess allele sharing with B. smaragdina
Excess allele sharing with B. mahachaiensis
Excess allele sharing with B. imbellis
Excess allele sharing with B. siamorientalis
L
Mah0
Mah1
Mah2
Frequency
f4-ratio (wild B. splendens, ornamental, B. mahachaiensis)

**Fig. S4. Excess allele sharing between *B. splendens* and ornamental betta. (A)** Pairwise genetic differences (above diagonal) and BBAA counts (below diagonal) between wild *B. splendens* populations. BBAA counts are based on Dsuite output where p3=*B. smaragdina*. **(B)** f4-ratio and *D* tests between individuals of wild *B. splendens* populations. Color of dots indicate block-jackknife significance as z-score of each test. Panel background colors indicate p3; ornamental individuals that are outliers, with high-average f4 or *D* are shown in different shapes (C-F)**.** Individual f4-ratio and *D* tests between wild *B. splendens* populations and ornamentals relative to *smaragdina* **(C)**, *mahachaiensis* **(D)**, *imbellis* **(E)**, and *siamorientalis* **(F)**. **(G-J)** Proportion of ABBA-BABA tests *D*(ornamentals except focal, focal ornamental; non-*splendens* species, *compuncta*), where a focal ornamental individual is significantly closer to a non-splendens species compared to other ornamental individuals (D>0 significant, Closer, blue) and fraction where a focal ornamental individual is significantly more distant from a non-splendens species compared to other ornamental individuals (D<0 significant, More distant, orange). Non-*splendens* species are *smaragdina* **(G)**, *mahachaiensis* **(H)**, *imbellis* **(I)**, and *siamorientalis* **(J)**. A z-score of 5.49 is used as significance cutoff corresponding to a Bonferroni corrected *P*=0.01. **(K)** Proportion of ABBA-BABA tests *D*(wild population same as focal, wild focal; non-*splendens* species, *compuncta*) that are significantly positive (Closer, blue) and negative (More distant, orange). Non-*splendens* species are *smaragdina*, *mahachaiensis*, *imbellis*, and *siamorientalis* (ordered top left to bottom right). A z-score of 5.49 is used as significance cutoff corresponding to a Bonferroni corrected *P*=0.01. **(L)** Excess allele sharing between *B. mahachaiensis* samples (Mah0, Mah1, Mah2) and ornamental betta relative to wild *B. splendens*. Individual f4-ratio tests with p1=ornamental betta, p2=wild *B. splendens*, and p3=Mah0 (blue), Mah1 (orange), or Mah2 (green). Note that for (D), (H), and (I) only Mah0 was used in the comparisons but not Mah1 and Mah2.

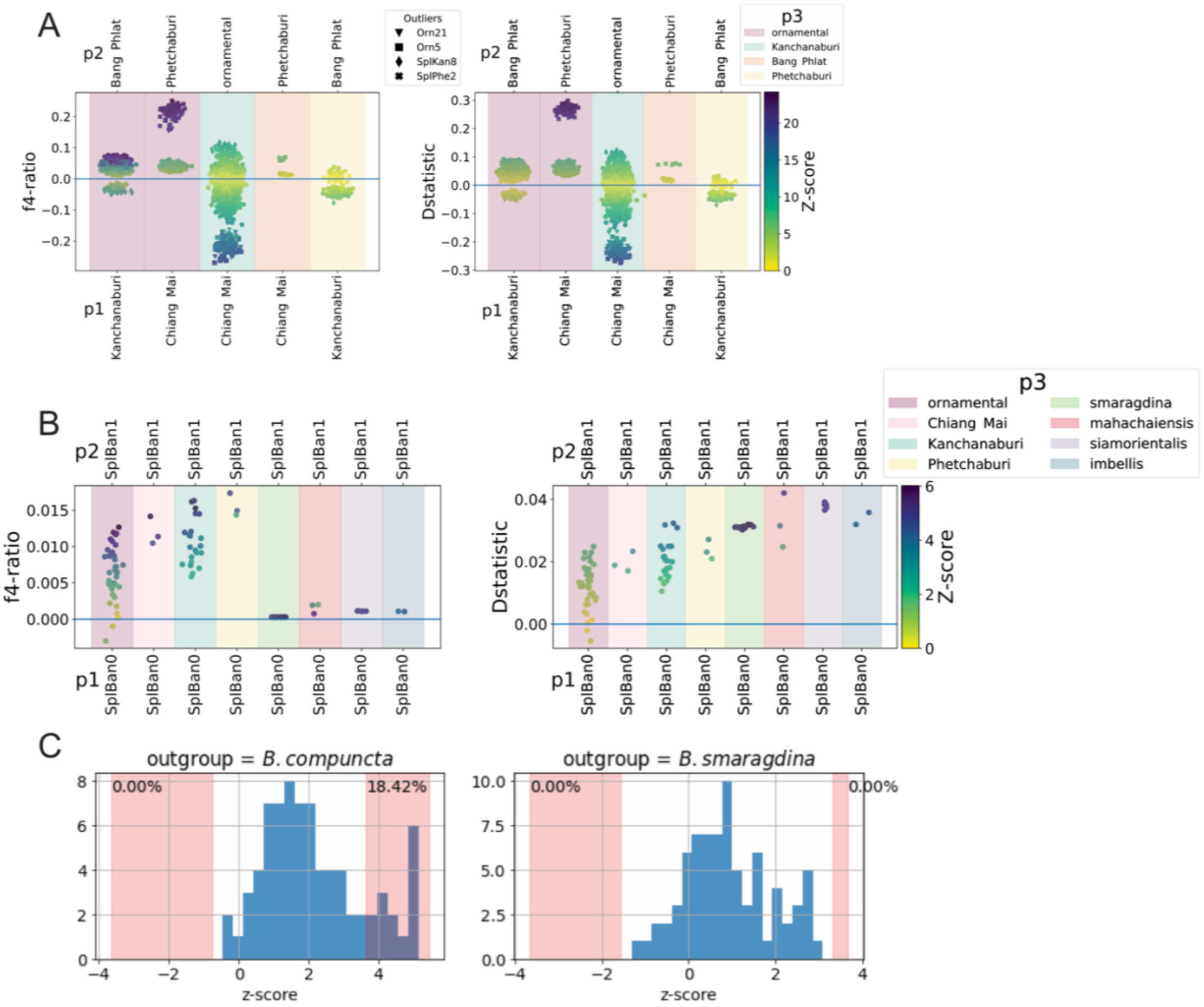


**Fig. S5. Excess allele sharing between *B. splendens* populations and ornamental betta. (A)** Excess allele sharing between wild *B. splendens* populations and ornamental betta. Individual f4 (left) or *D* (right) (p1, p2, p3, *B. compuncta*) test, where p1, p2, p3 individuals are taken from different *B. splendens* populations and ornamental betta so that ((p1, p2), p3) is consistent with the genome-wide phylogeny in Suppl. Fig. 2A. Outlier individuals are represented by different markers. Color of markers indicates block-jackknife significance as z-score; panel background colors indicate p3 **(A,B)**. **(B)** Excess allele sharing of SplBan1 relative to SplBan0 with individuals of ornamental betta, other wild *B. splendens* populations and non-*splendens* species. **(C)** Distribution of z-scores of individual Patterson's D tests of the form D(SplBan0, SplBan1; p3 , outgroup), where p3 represents each individual sample except SplBan0, SplBan1 and samples from *B. smaragdina*. The outgroup is *B. compuncta* in the left panel and *B. smaragdina* in the right panel. Pink shaded areas correspond to z-scores of Bonferroni multiple-testing corrected *P*-values < 0.01. Percentages correspond to the percentage of comparisons in which SplBan0 (left) and SplBan1(right) individuals are significantly closer to other individuals.

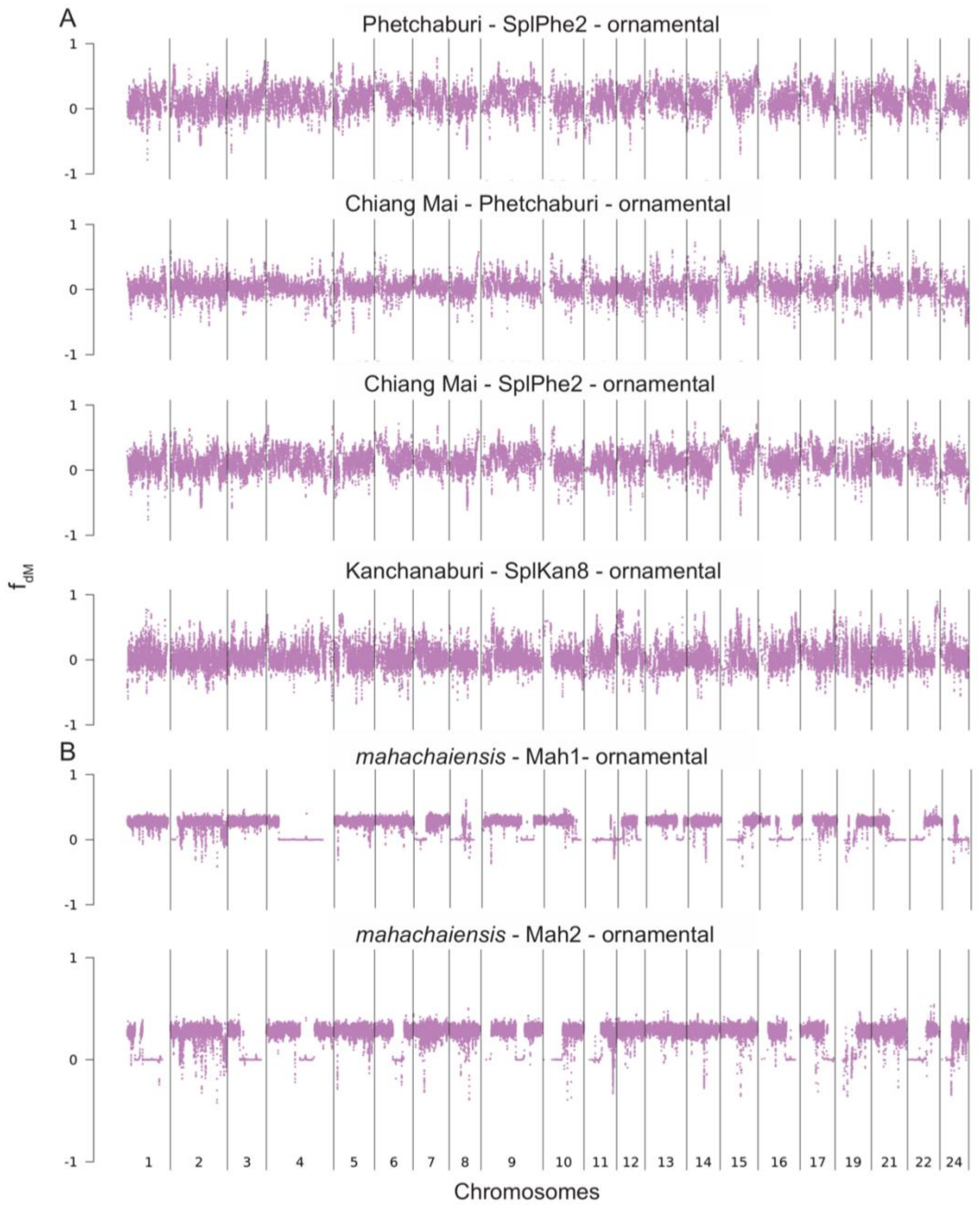


**Fig. S6. Genome-wide $f_{dM}$ for wild *B. splendens* and non-*splendens* to ornamental betta.** Genome-wide $f_{dM}$ plots with ordering specified on the title in the format (p1 - p2 - p3). The outgroup p4 is *B. compuncta*. Each point represents 100 SNPs. Positive $f_{dM}$ values measure excess allele sharing between p2 and p3 (ref(*62*)), **(A)** wild *B. splendens* individuals (p2) with ornamental introgression. **(B)** *B. mahachaiensis* samples with ornamental introgression.

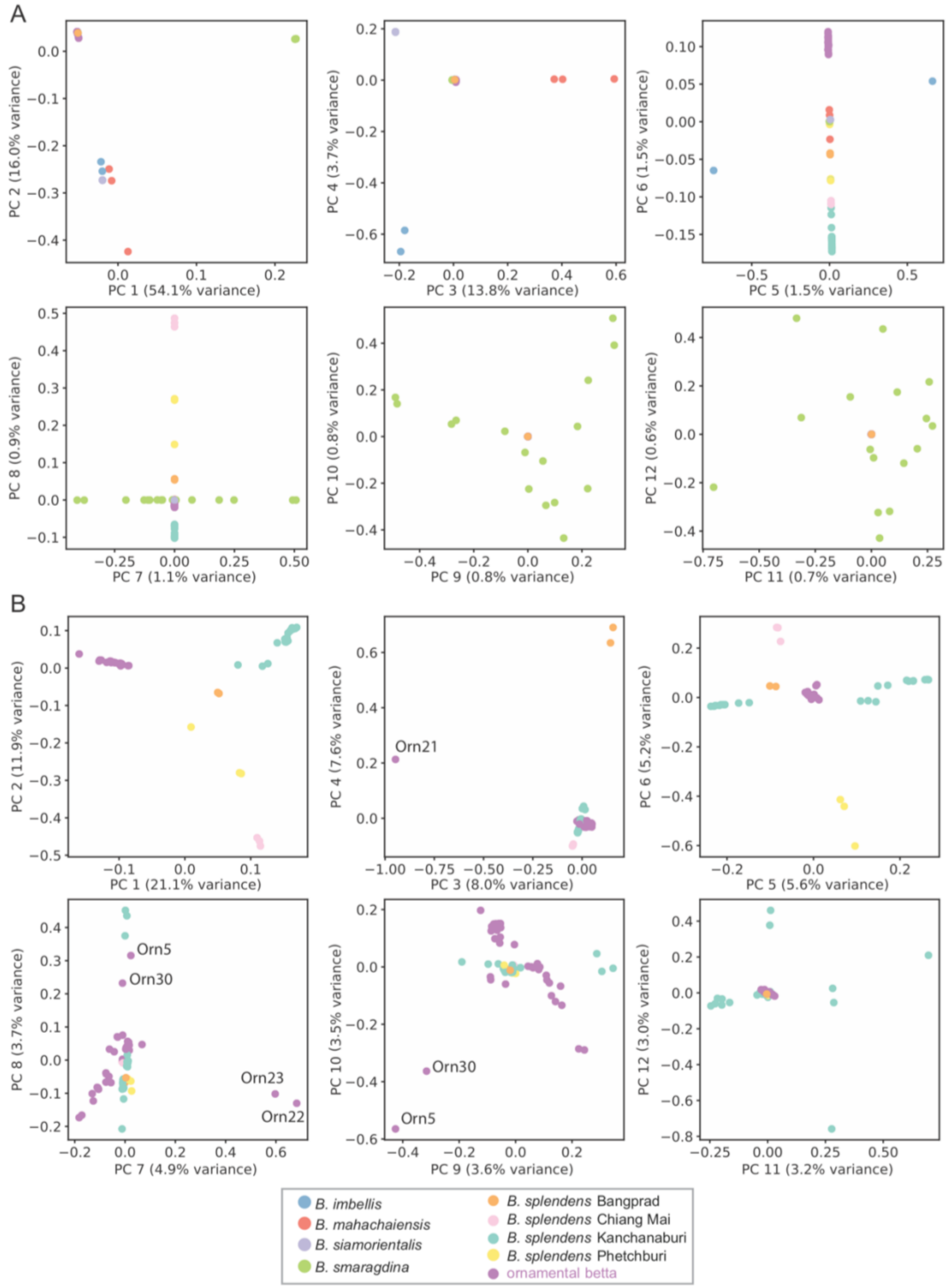


**Fig. S7. Population structure across species of *B. splendens* species complex and populations of *B. splendens*. (A)** Principal component analysis of all samples of the *B. splendens* species complex including ornamental betta. **(B)** Principal component analysis of *B.*

*splendens* samples including ornamental betta. Color codes across all PCA analysis are provided at the bottom.

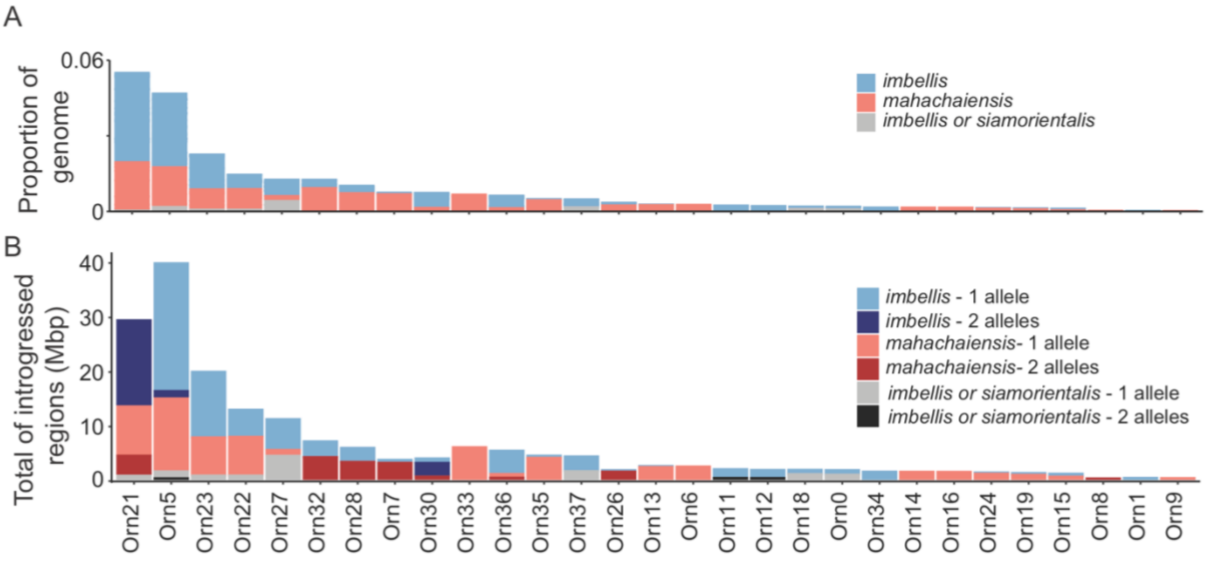


**Fig. S8. Allele-sharing from non-*splendens* into ornamental betta.** Introgression into ornamental betta as determined by $f_{dM}$ in genomic windows of 100 SNPs with 25 SNPs slide. **(A)** Proportion of genome with introgression per ornamental betta, color coded by ancestry. **(B)** Sum of introgressed regions. Lighter shading indicates regions where introgression occurs on a single allele; darker, both alleles. Grey shading indicates ancestry which could not be assigned exclusively to either *B. siamorientalis* or *B. imbellis*. Ordering of ornamental betta is based on decreasing proportion of introgressed genome, and is the same for both panels. We found significant high-$f_{dM}$ regions in one more individual than with the genome-wide D statistic. This likely reflects a higher sensitivity of $f_{dM}$ afforded by scanning for non-*splendens* ancestry in small genomic windows.

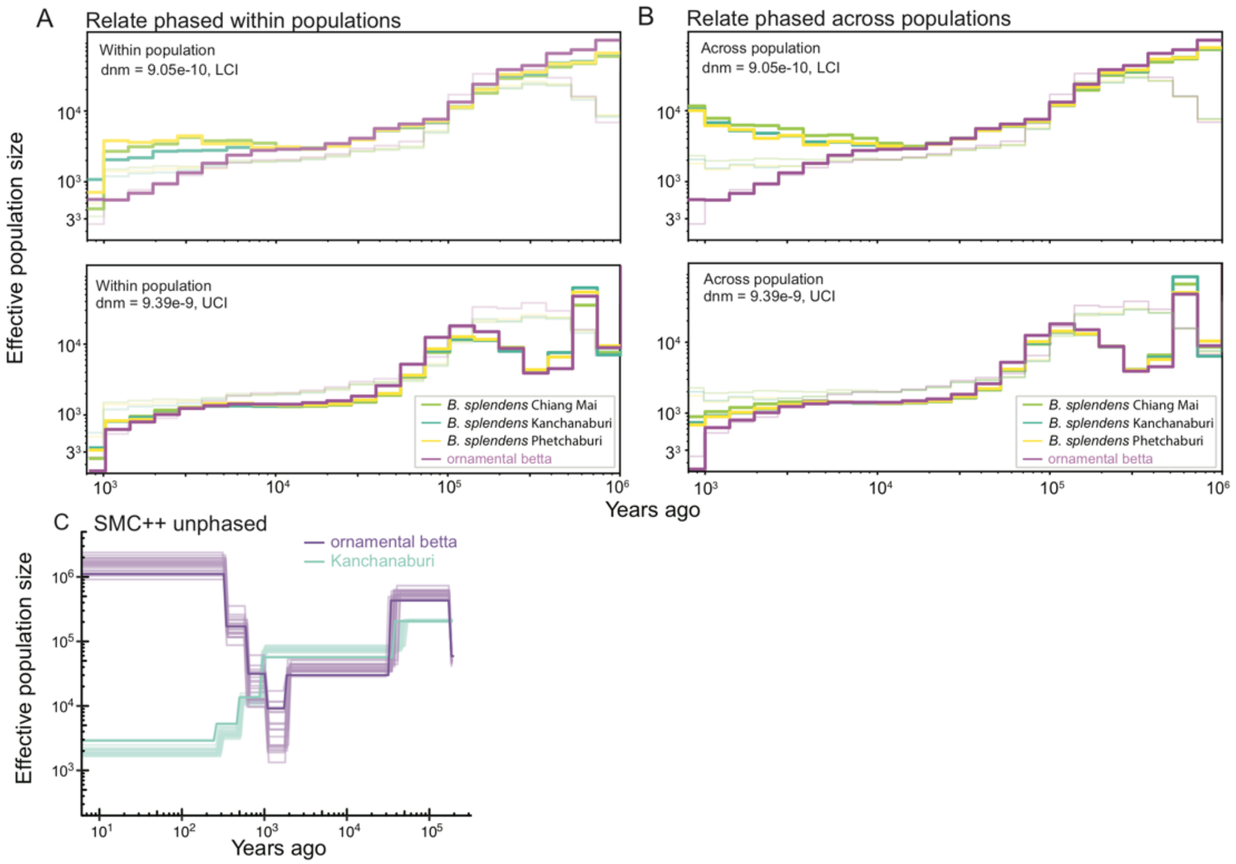


**Fig. S9. Effective population size histories. (A)** Within-population demographic estimates across ornamentals and wild *B. splendens* populations using Relate. Upper panel: demography using the upper confidence interval (UCI) for the mutation rate ($9.39 \times 10^{-9}$ mutations per bp per generation); lower panel: lower confidence interval (LCI) ($9.05 \times 10^{-10}$). Thin, lighter lines in each panel are inferences generated with the point estimate of the mutation rate ($3.75 \times 10^{-9}$ mutations per bp per generation) **(A,B)**. **(B)** Across-population demographic estimates between ornamentals and wild *B. splendens* populations using Relate. **(C)** Demographic estimation using SMC++ with unphased genomes of ornamental and wild *B. splendens* sampled from Kanchanaburi with jack-knife resampling of chromosomes. De novo mutation rate used for both demographic inferences is $3.75 \times 10^{-9}$ mutations per bp per generation. (A-C) Generation time is six months.

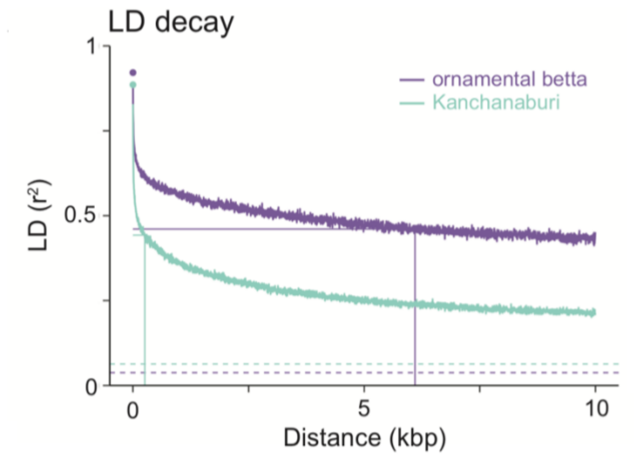


**Fig. S10. Linkage disequilibrium decay.** Linkage disequilibrium decay of ornamental and wild *B. splendens* from Kanchanaburi. Half- max for ornamentals: 6.1 kbp; for wild: 256 bp. Horizontal dashed lines denote interchromosomal $r^2$ (ornamental: 0.038; wild: 0.063).

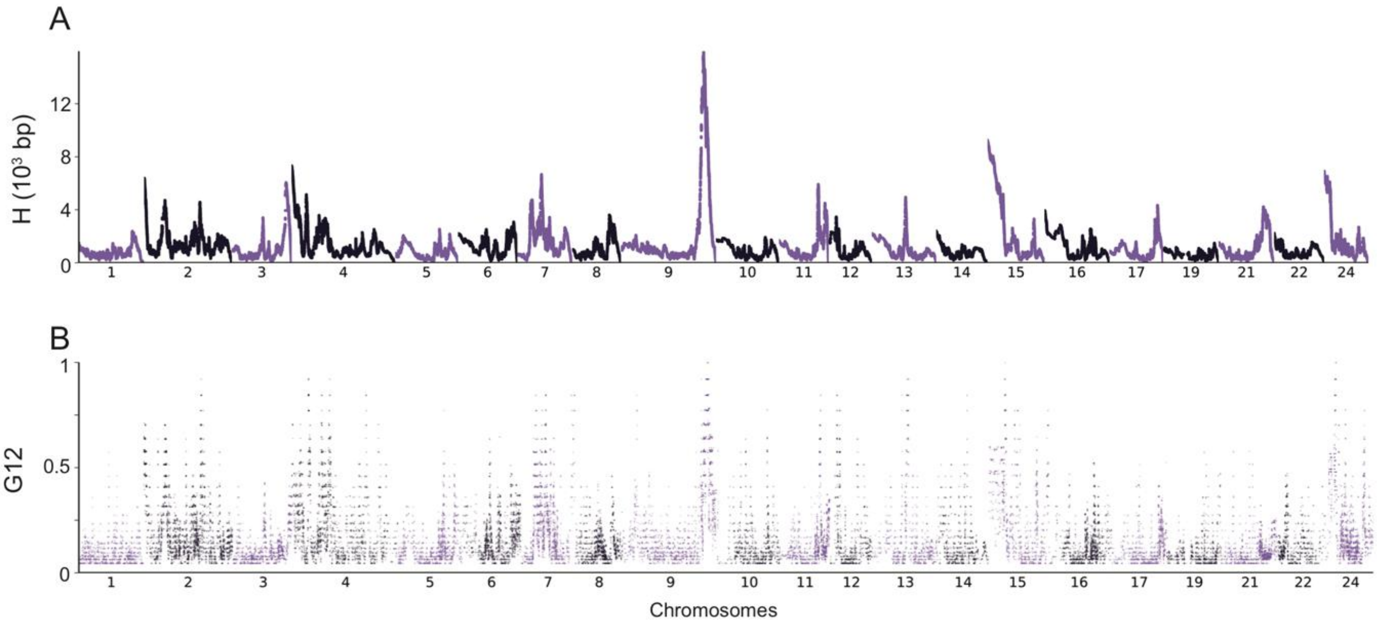


**Fig. S11. H-scan and G12 selection scans across ornamental subsets. (A)** Genome-wide H-scan of randomly sampled 24 independent ornamental bettas. **(B)** Genome-wide G12 of randomly sampled 24 independent ornamental bettas.

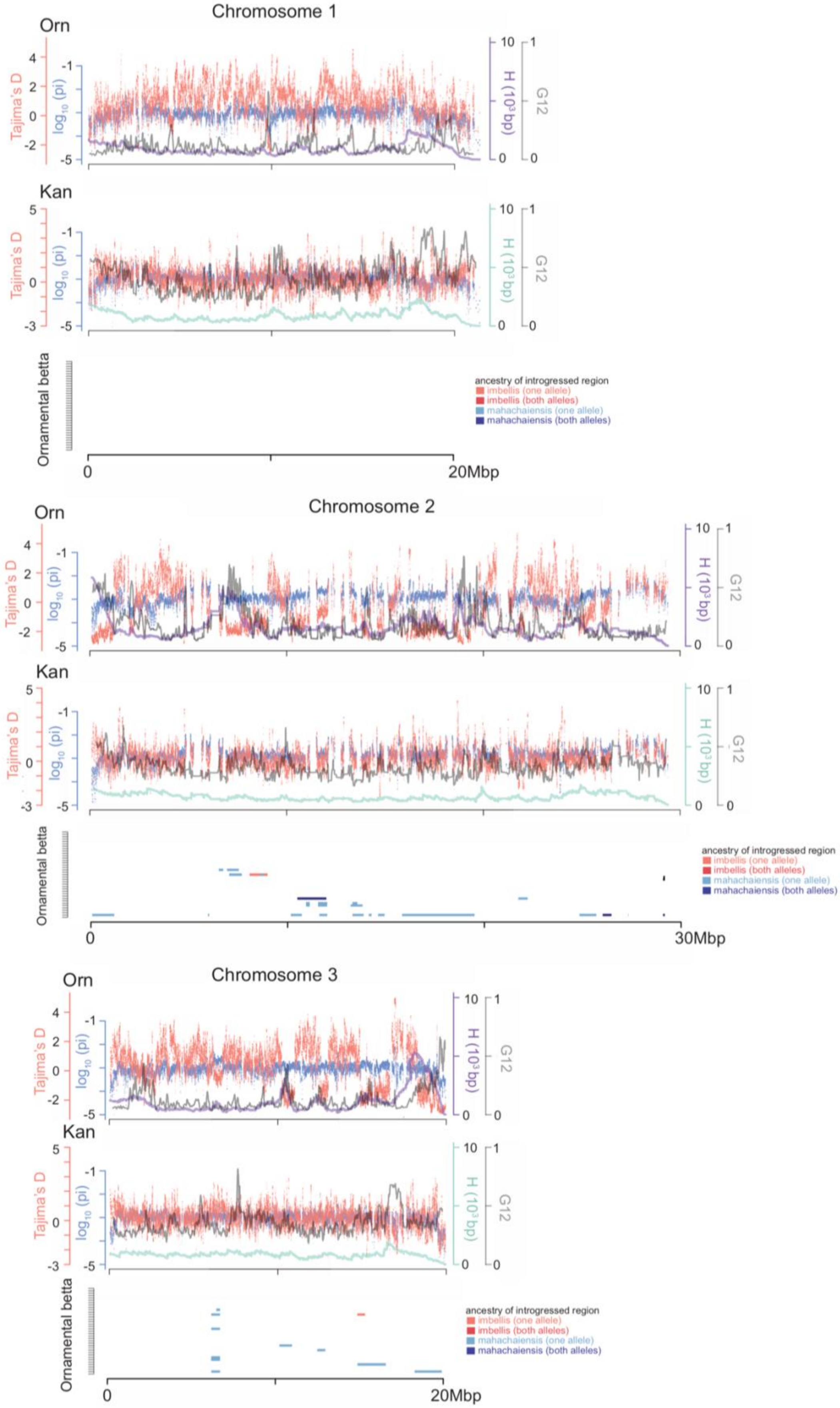
Chromosome 1
Orn
Kan
Tajima's D
log10 (pi)
H (10³bp)
G12
Ornamental betta
ancestry of introgressed region
imbellis (one allele)
imbellis (both alleles)
mahachaiensis (one allele)
mahachaiensis (both alleles)
0
20Mbp
Chromosome 2
30Mbp
Chromosome 3

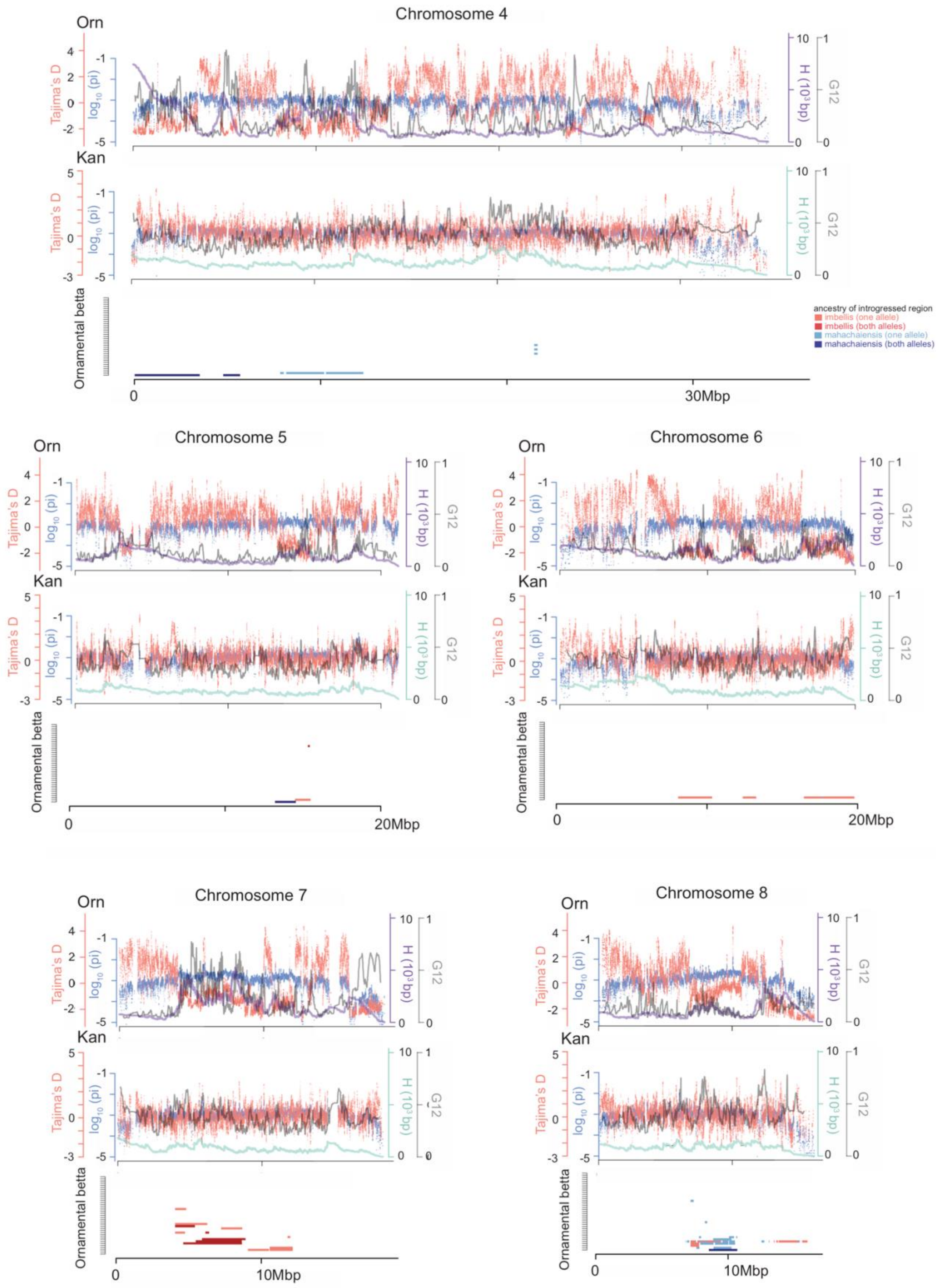
Chromosome 4
Orn
Kan
Tajima's D
log10 (pi)
H (10³bp)
G12
Ornamental betta
ancestry of introgressed region
imbellis (one allele)
imbellis (both alleles)
mahachaiensis (one allele)
mahachaiensis (both alleles)
0
30Mbp
Chromosome 5
0
20Mbp
Chromosome 6
0
20Mbp
Chromosome 7
0
10Mbp
Chromosome 8
0
10Mbp

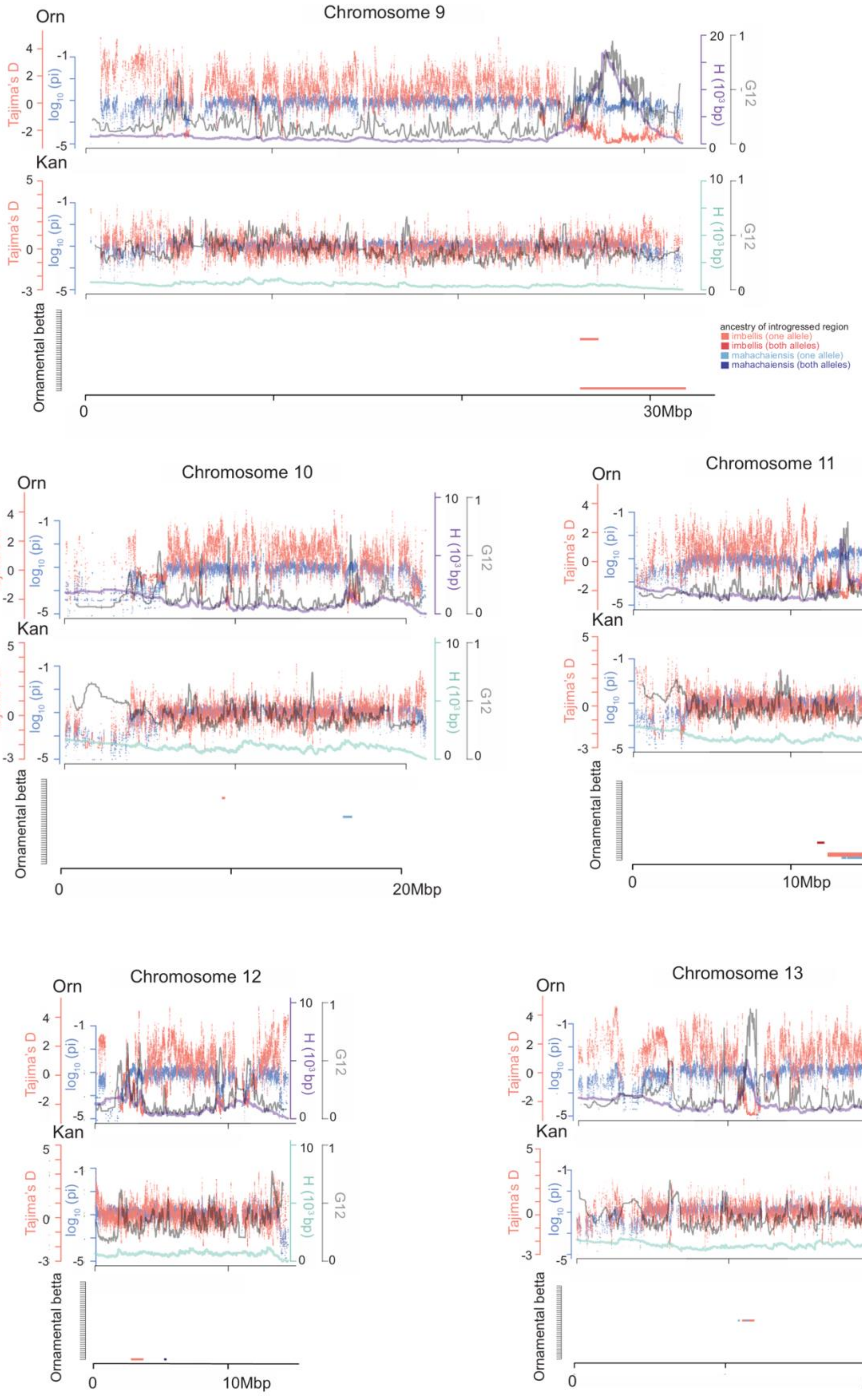
Chromosome 9
Orn
Kan
Tajima's D
log10 (pi)
H (10³bp)
G12
Ornamental betta
ancestry of introgressed region
imbellis (one allele)
imbellis (both alleles)
mahachaiensis (one allele)
mahachaiensis (both alleles)
0
30Mbp
Chromosome 10
20Mbp
Chromosome 11
10Mbp
Chromosome 12
10Mbp
Chromosome 13
20Mbp

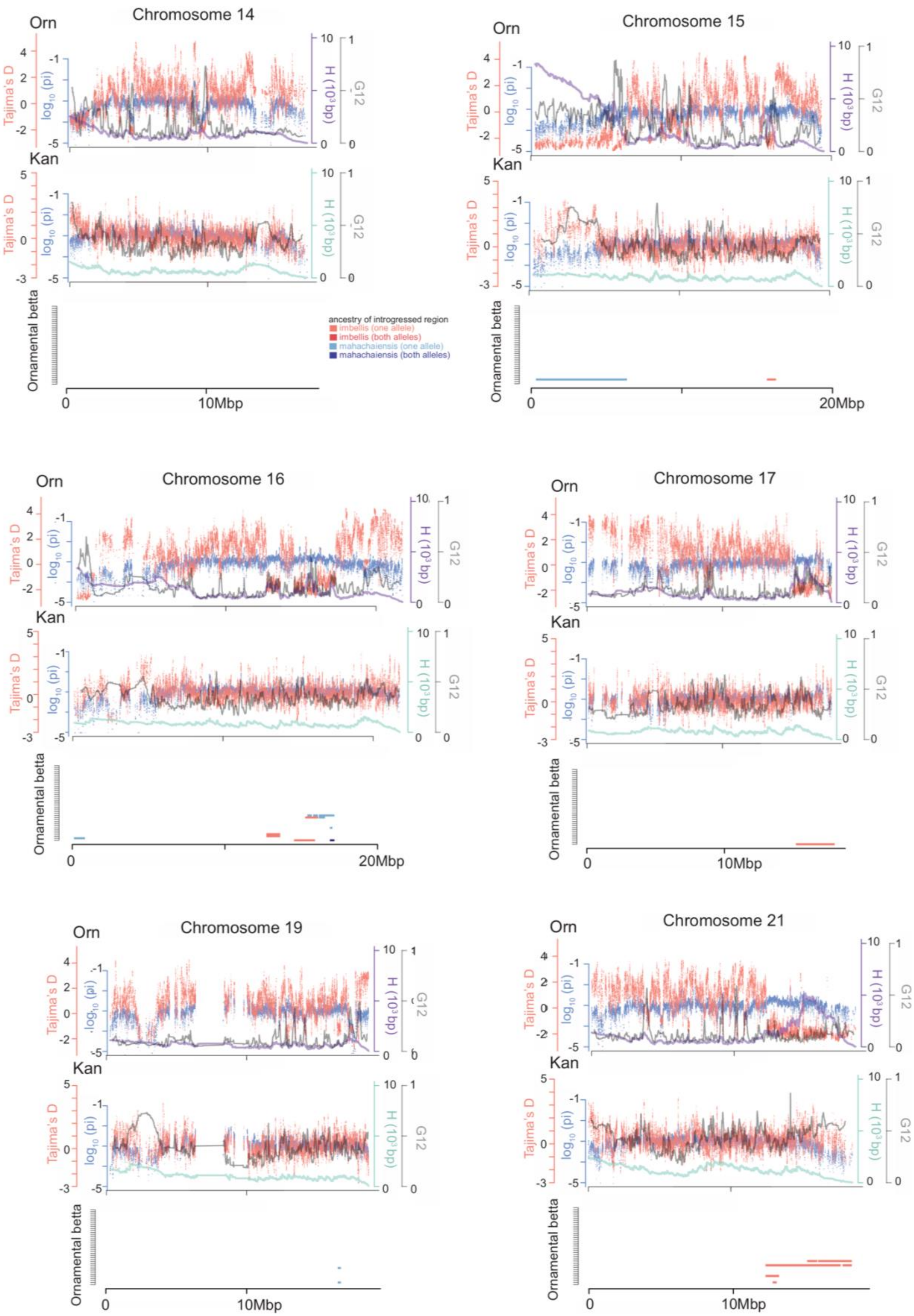
Chromosome 14
Chromosome 15
Chromosome 16
Chromosome 17
Chromosome 19
Chromosome 21
Orn
Kan
Tajima's D
log10 (pi)
H (10³bp)
G12
Ornamental betta
ancestry of introgressed region
imbellis (one allele)
imbellis (both alleles)
mahachaiensis (one allele)
mahachaiensis (both alleles)
0
10Mbp
20Mbp

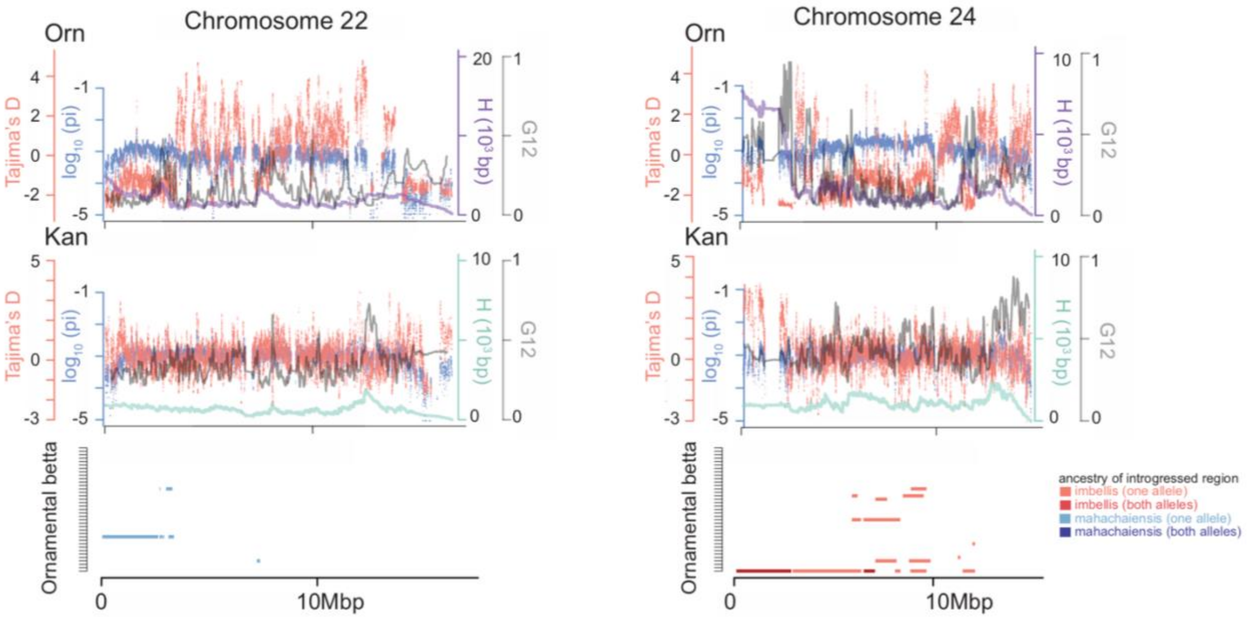


**Fig. S12. Genome diversity and selection scans of ornamental betta and wild *B. splendens*.** Per chromosome plots of nucleotide diversity (pi) and Tajima's D in 10 kb windows with 1 kb slide, as well as H-scan, and G12 in ornamental (top panel of each chromosome) and wild *B. splendens* from Kanchanaburi (middle panel). Bottom panel per chromosome: plot of regions with $f_{dM}$>0.2 per ornamental betta where p1 is the ornamental betta population and p3 is *imbellis* (red) or *mahachaiensis* (blue) populations. Lighter shaded regions indicate introgression from either *B. imbellis* or *B. mahachaiensis* on one allele. Darker shading introgressions observed in both alleles. Each row represents an ornamental individual.

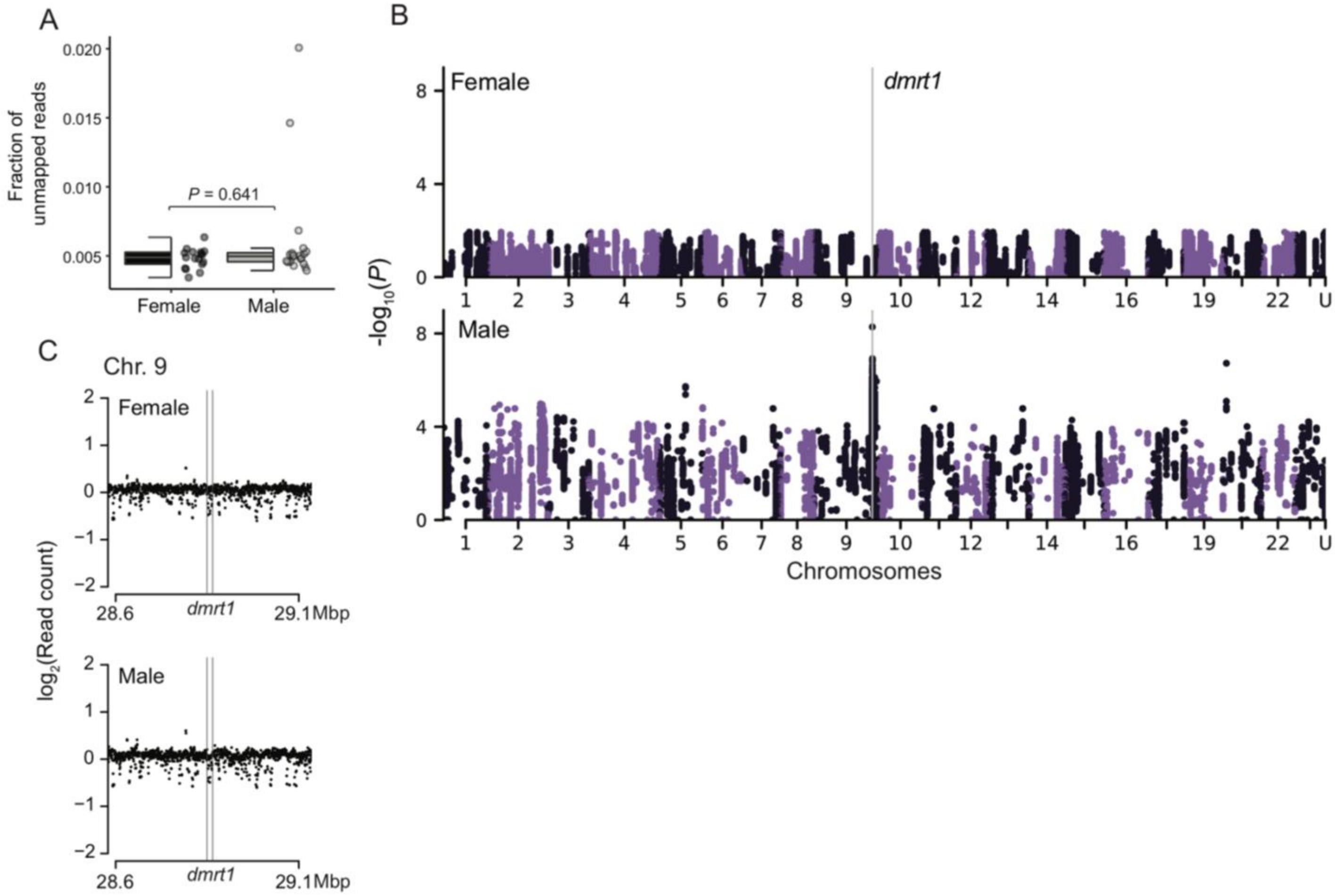


**Fig. S13. K-mer and read-depth analysis across sex. (A)** Number of unmapped over total reads across sex, significance calculated by Mann-Whitney U test. **(B)** k-mer genome-wide association plot with unmapped assembled contigs placed at the end of chromosome 24, denoted U. **(C)** Average $log_2$-normalized read depth in 1 kbp bins with 500 bp slide across *dmrt1* on chromosome 9 for female and male ornamental betta.

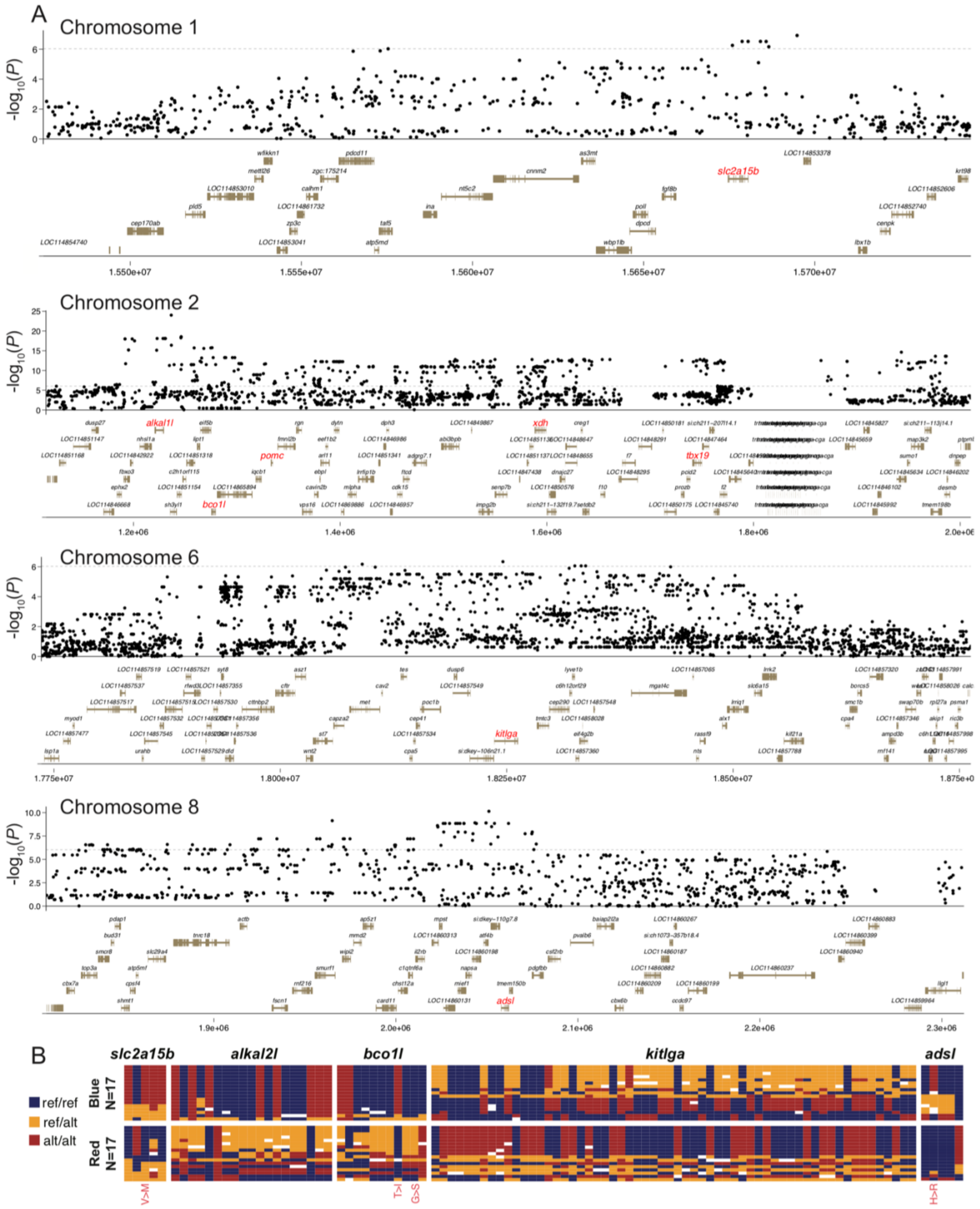


**Fig. S14. Extended red-blue GWAS peak plots. (A)** Expanded view of the red-blue color

GWAS peaks with gene annotations across chromosomes 1, 2, 6, and 8. Candidate genes with high -log10(*P*)values are highlighted in red. **(B)** Genotypes of red and blue fish included in GWAS across genes of interest. SNPs leading to amino acid changes are annotated in red.

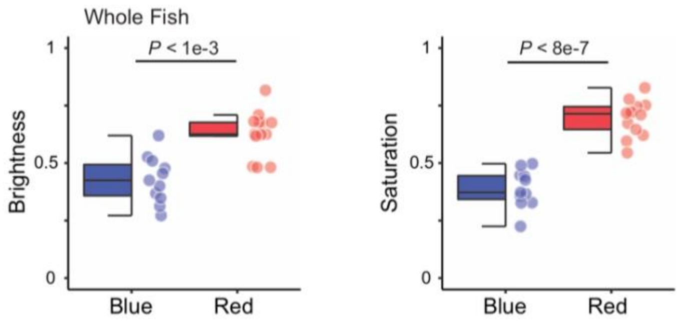


**Fig. S15. Coloration of red and blue P0 Fish.** Average brightness and saturation between P0 red and blue fish. Significance calculated by Mann-Whitney U test.

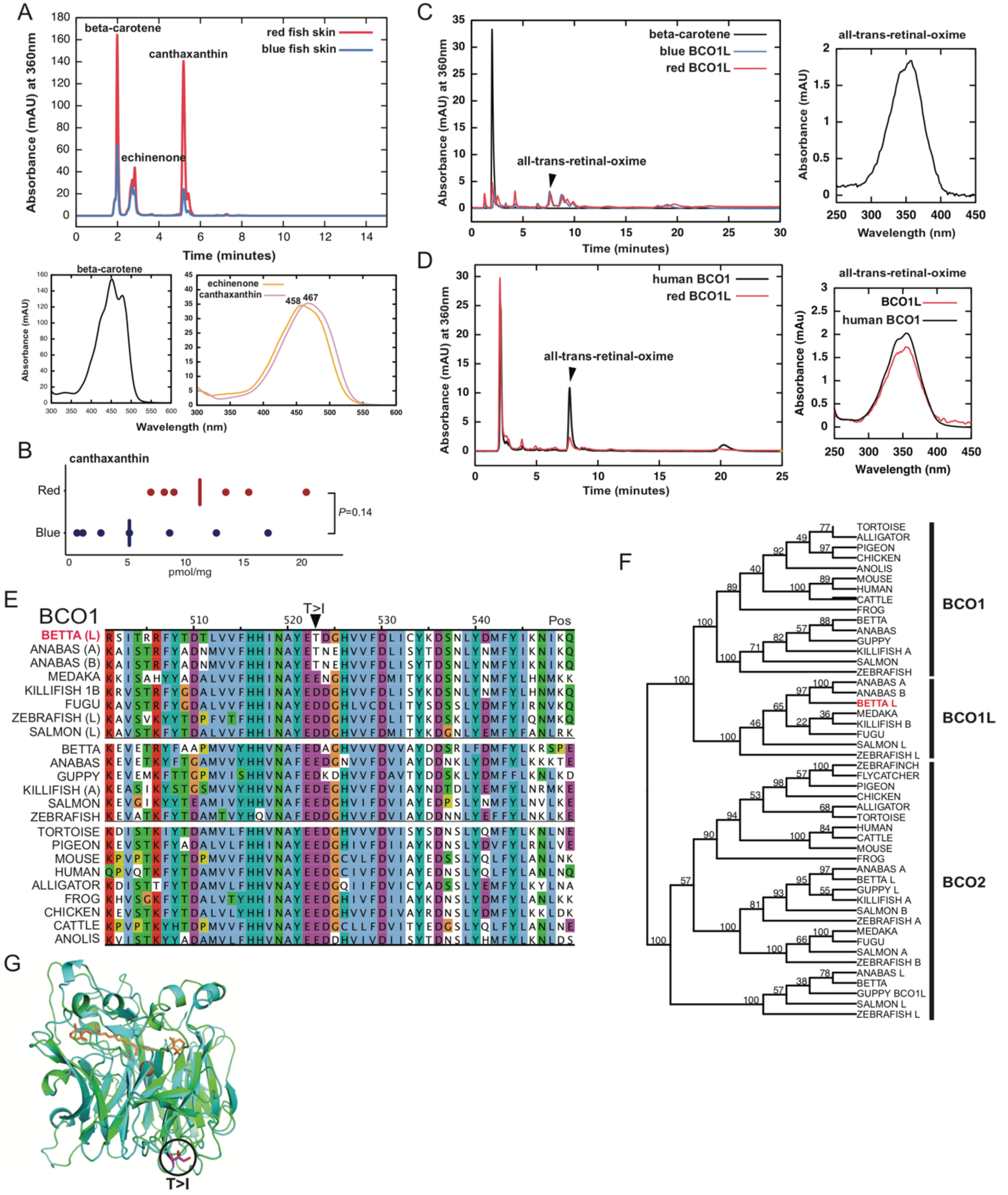

**Fig. S16. Carotenoid and β-carotene oxygenase 1-like biochemistry. (A)** HPLC trace of extracts from red and blue ornamental betta skins. **(B)** Concentration of canthaxanthin in red (n=6) and blue (n=6) ornamental betta skin. Significance by Mann-Whitney U test. **(C)** HPLC trace of lipid extracts from *E. coli* strain that accumulates β-carotene, expressing the recombinant MBP-BCO1L red and blue fusion proteins. **(D)** HPLC trace of products of the purified hBCO1 and red-allele BCO1L enzymes, showing the 15,15′-dioxygenase product all-trans-retinal-oxime. **(E)** Multi-species amino acid alignment of BCO1L using MAFFT, color coded with the Clustal X color scheme with consensus sequence and frequency below. **(F)** Neighbor-joining tree of 100 bootstraps of the BCO amino acid alignment across vertebrates. **(G)** Predicted monomer model of BCO1L, orange shows β-carotene in the catalytic site, magenta indicates the location of the T>I (red, wildtype allele to blue allele) mutation.

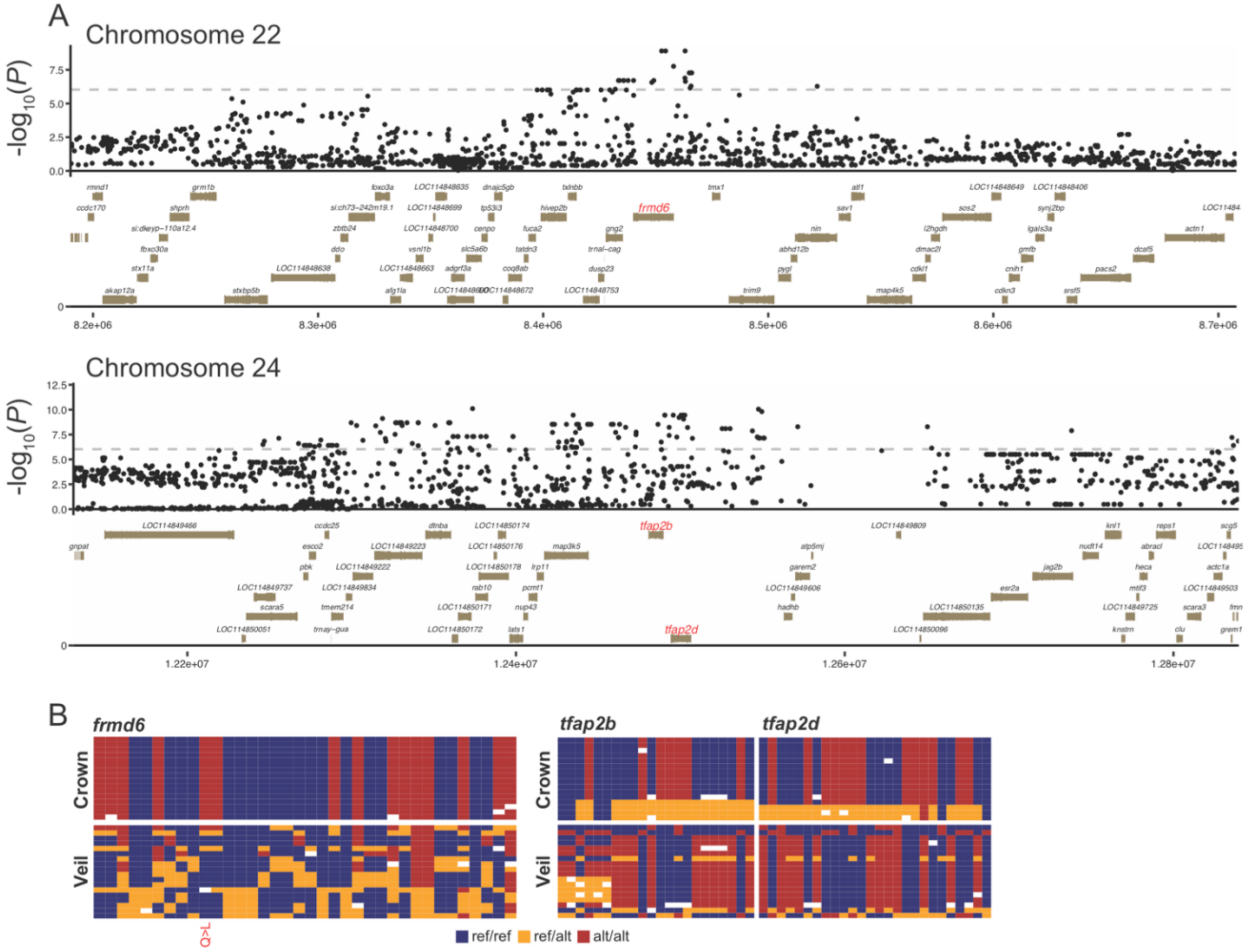


**Fig. S17. Extended crown-veil fin GWAS and genotype plots. (A)** Expanded view of the crown-veil fin GWAS peaks with gene annotations across Chromosomes 22 and 24. Candidate genes with high -log10(*P*) values are highlighted in red. **(B)** Genotypes of crown and veil fish included in GWAS across genes of interest. SNPs leading to amino acid changes are annotated in red.

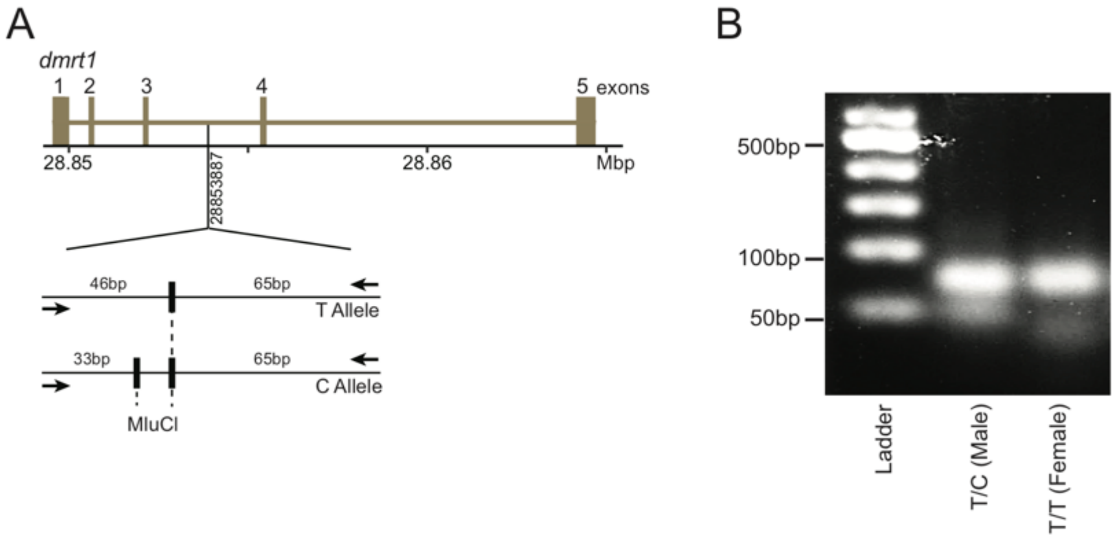


**Fig. S18. Restriction fragment length polymorphism (RFLP) genotyping. (A)** Schematic of the MluCl restriction enzyme cut sites (bolded vertical black bars) and the lengths of the digested PCR product for alleles carrying either the T or C variant at chromosome 9: 288533887. **(B)** 2.5% agarose gel electrophoresis of digested fragments from a male *dmrt1_XY* carrying the T/C genotype and female *dmrt1_XX* carrying the T/T genotype.

**Table S1. Sample metadata.** Metadata information of the whole-genome sequenced samples used in population genetic analyses.

| pub name | secondary ID | paternal ID | maternal ID | sex | color | marble | fin type | species | population |
|---|---|---|---|---|---|---|---|---|---|
| Com0 | Anaban27350502 | - | - | - | - | - | - | compuncta | - |
| Imb0 | 4616STDY6773202 | - | - | - | - | - | - | imbellis | - |
| Imb1 | 4616STDY7068352 | - | - | - | - | - | - | imbellis | - |
| Mah0 | 4616STDY7068355 | - | - | - | - | - | - | mahachaiensis | - |
| Mah1 | Anaban27350532 | - | - | Female | - | - | - | mahachaiensis | - |
| Mah2 | Anaban27350533 | - | - | Male | - | - | - | mahachaiensis | - |
| Sia0 | Anaban27350534 | - | - | - | - | - | - | siamorientalis | southern Vietnam |
| Sia1 | Anaban27350535 | - | - | - | - | - | - | siamorientalis | southern Vietnam |
| Sia2 | Anaban27350536 | - | - | - | - | - | - | siamorientalis | southern Vietnam |
| Sia3 | Anaban27350537 | - | - | - | - | - | - | siamorientalis | southern Vietnam |
| Sia4 | Anaban27350538 | - | - | - | - | - | - | siamorientalis | southern Vietnam |
| Sia5 | Anaban27350539 | - | - | - | - | - | - | siamorientalis | southern Vietnam |
| Sma0 | Anaban27350516 | - | - | - | - | - | - | smaragdina | southern Thailand border Cambodia |
| Sma1 | Anaban27350517 | - | - | - | - | - | - | smaragdina | southern Thailand border Cambodia |
| Sma2 | Anaban27350518 | - | - | - | - | - | - | smaragdina | southern Thailand border Cambodia |
| Sma3 | Anaban27350519 | - | - | - | - | - | - | smaragdina | southern Thailand border Cambodia |
| Sma4 | Anaban27350520 | - | - | - | - | - | - | smaragdina | southern Thailand border Cambodia |
| Sma5 | Anaban27350521 | - | - | - | - | - | - | smaragdina | southern Thailand border Cambodia |
| Sma6 | Anaban27350522 | - | - | - | - | - | - | smaragdina | southern Thailand border Cambodia |
| Sma7 | Anaban27350523 | - | - | - | - | - | - | smaragdina | southern Thailand border Cambodia |
| Sma8 | Anaban27350524 | - | - | - | - | - | - | smaragdina | southern Thailand border Cambodia |
| Sma9 | Anaban27350525 | - | - | - | - | - | - | smaragdina | southern Thailand border Cambodia |
| Sma10 | Anaban27350526 | - | - | - | - | - | - | smaragdina | southern Thailand border Cambodia |
| Sma11 | Anaban27350527 | - | - | - | - | - | - | smaragdina | southern Thailand border Cambodia |

| | | | | | | | | | |
|---|---|---|---|---|---|---|---|---|---|
| Sma12 | Anaban273 50528 | - | - | - | - | - | - | smaragdina | southern Thailand border Cambodia |
| Sma13 | Anaban273 50529 | - | - | - | - | - | - | smaragdina | southern Thailand border Cambodia |
| Sma14 | Anaban273 50530 | - | - | - | - | - | - | smaragdina | southern Thailand border Cambodia |
| Sma15 | Anaban273 50531 | - | - | - | - | - | - | smaragdina | southern Thailand border Cambodia |
| SplBan0 | 4616STDY 7556631 | - | - | Male | - | - | - | splendens | Bang Phlat |
| SplBan1 | 4616STDY 8045966 | - | - | Juvenile | - | - | - | splendens | Bang Phlat |
| SplChi0 | 4616STDY 8045967 | - | - | Male | - | - | - | splendens | Chiang Mai |
| SplChi1 | 4616STDY 8045968 | - | - | Juvenile | - | - | - | splendens | Chiang Mai |
| SplChi2 | 4616STDY 8045969 | - | - | Juvenile | - | - | - | splendens | Chiang Mai |
| SplKan0 | Anaban273 50564 | - | - | Female | - | - | - | splendens | Kanchanaburi |
| SplKan1 | Anaban273 50565 | - | - | Female | - | - | - | splendens | Kanchanaburi |
| SplKan2 | Anaban273 50566 | - | - | Female | - | - | - | splendens | Kanchanaburi |
| SplKan3 | Anaban273 50567 | - | - | Female | - | - | - | splendens | Kanchanaburi |
| SplKan4 | Anaban273 50568 | - | - | Female | - | - | - | splendens | Kanchanaburi |
| SplKan5 | Anaban273 50569 | - | - | Female | - | - | - | splendens | Kanchanaburi |
| SplKan6 | Anaban273 50570 | - | - | - | - | - | - | splendens | Kanchanaburi |
| SplKan7 | Anaban273 50571 | - | - | Female | - | - | - | splendens | Kanchanaburi |
| SplKan8 | Anaban273 50572 | - | - | Male | - | - | - | splendens | Kanchanaburi |
| SplKan9 | Anaban273 50573 | - | - | Male | - | - | - | splendens | Kanchanaburi |
| SplKan10 | Anaban273 50574 | - | - | Male | - | - | - | splendens | Kanchanaburi |
| SplKan11 | Anaban273 50575 | - | - | Male | - | - | - | splendens | Kanchanaburi |
| SplKan12 | Anaban273 50576 | - | - | Male | - | - | - | splendens | Kanchanaburi |
| SplKan13 | Anaban273 50577 | - | - | Male | - | - | - | splendens | Kanchanaburi |
| SplKan14 | Anaban273 50578 | - | - | Male | - | - | - | splendens | Kanchanaburi |
| SplKan15 | Anaban273 50579 | - | - | Male | - | - | - | splendens | Kanchanaburi |
| SplKan16 | Anaban273 50580 | - | - | Male | - | - | - | splendens | Kanchanaburi |
| SplKan17 | Anaban273 50581 | - | - | Male | - | - | - | splendens | Kanchanaburi |
| SplKan18 | Anaban273 50582 | - | - | Male | - | - | - | splendens | Kanchanaburi |
| SplKan19 | Anaban273 50583 | - | - | Male | - | - | - | splendens | Kanchanaburi |
| SplKan20 | Anaban273 50584 | - | - | Male | - | - | - | splendens | Kanchanaburi |
| SplKan21 | Anaban273 50585 | - | - | Male | - | - | - | splendens | Kanchanaburi |
| SplPhe0 | 4616STDY 8045963 | - | - | Male | - | - | - | splendens | Phetchaburi |
| SplPhe1 | 4616STDY 8045964 | - | - | Juvenile | - | - | - | splendens | Phetchaburi |

| | | | | | | | | | |
|---|---|---|---|---|---|---|---|---|---|
| SplPhe2 | 4616STDY 8045965 | - | - | Juvenile | - | - | - | splendens | Phetchaburi |
| Orn0 | 9 | 0 | 0 | Female | blue | n | veil | splendens | ornamental |
| Orn1 | 21 | 0 | 0 | Female | red | n | half-moon | splendens | ornamental |
| Orn2 | 23 | 0 | 0 | Male | red | n | veil | splendens | ornamental |
| Orn3 | 26 | 0 | 0 | Female | red | n | veil | splendens | ornamental |
| Orn4 | 29 | 0 | 0 | Male | blue | n | veil | splendens | ornamental |
| Orn5 | 45 | 0 | 0 | Male | black | n | half-moon | splendens | ornamental |
| Orn6 | 47 | 0 | 0 | Male | blue | n | crown | splendens | ornamental |
| Orn7 | 48 | 0 | 0 | Male | red | n | crown | splendens | ornamental |
| Orn8 | 49 | 0 | 0 | Male | red | n | crown | splendens | ornamental |
| Orn9 | 50 | 0 | 0 | Male | red | n | veil | splendens | ornamental |
| Orn10 | 51 | 0 | 0 | Female | red | n | veil | splendens | ornamental |
| Orn11 | 53 | 0 | 0 | Female | blue | n | veil | splendens | ornamental |
| Orn12 | 55 | 0 | 0 | Male | blue | n | veil | splendens | ornamental |
| Orn13 | 56 | 0 | 0 | Female | blue | n | crown | splendens | ornamental |
| Orn14 | 118 | 0 | 0 | Female | red | n | crown | splendens | ornamental |
| Orn15 | 120 | 0 | 0 | Male | blue | n | crown | splendens | ornamental |
| Orn16 | 122 | 0 | 0 | Male | red | n | crown | splendens | ornamental |
| Orn17 | 129 | 0 | 0 | Female | red | n | half-moon | splendens | ornamental |
| Orn18 | 140 | 0 | 0 | Male | blue | n | veil | splendens | ornamental |
| Orn19 | 141 | 0 | 0 | Male | red | n | crown | splendens | ornamental |
| Orn20 | 175 | 0 | 0 | Male | red | n | veil | splendens | ornamental |
| Orn21 | 176 | 0 | 0 | Male | marble | y | crown | splendens | ornamental |
| Orn22 | 179 | 0 | 0 | Female | red | y | veil | splendens | ornamental |
| Orn23 | 180 | 0 | 0 | Female | orange | y | veil | splendens | ornamental |
| Orn24 | 183 | 0 | 0 | Male | red | y | crown | splendens | ornamental |
| Orn25 | 184 | 0 | 0 | Male | red | n | veil | splendens | ornamental |
| Orn26 | 185 | 0 | 0 | Female | blue | n | crown | splendens | ornamental |
| Orn27 | 187 | 0 | 0 | Male | blue | n | veil | splendens | ornamental |
| Orn28 | 188 | 0 | 0 | Male | blue | n | crown | splendens | ornamental |
| Orn29 | 189 | 45 | 26 | Male | marble | y | - | splendens | ornamental |
| Orn30 | 412 | 0 | 0 | Female | blue | n | veil | splendens | ornamental |
| Orn31 | 486 | 0 | 0 | Male | red | n | veil | splendens | ornamental |
| Orn32 | 642 | 0 | 0 | Female | blue | n | crown | splendens | ornamental |
| Orn33 | 644 | 0 | 0 | Female | red | n | crown | splendens | ornamental |
| Orn34 | 671 | 0 | 0 | Male | blue | n | veil | splendens | ornamental |
| Orn35 | 685 | 0 | 0 | Female | blue | n | veil | splendens | ornamental |
| Orn36 | 686 | 0 | 0 | Female | blue | n | crown | splendens | ornamental |
| Orn37 | 722 | 0 | 0 | Female | blue | n | crown | splendens | ornamental |
| Orn38 | 770 | 671 | 722 | Male | blue | n | crown, veil | splendens | ornamental |
| Orn39 | 772 | 671 | 722 | Female | blue | n | crown, veil | splendens | ornamental |

**Table S2. Chromosomal locations of loci and genes affecting phenotypic traits in betta.**

fBetSpl5.3 genomic coordinates of genes and genomic loci associated with phenotypic traits described in this study and Wang et al. 2021 (*15*).

| Phenotype | Putative gene | Gene name | Chromosome : Start-End |
|---|---|---|---|
| sex determination | dmrt1 | doublesex and mab-3 related transcription factor 1 | 9: 28,849,542-28,864,704 |
| iridophore differentiation | alkal2l | LOC114851404; ALK and LTK ligand 1-like (renamed to alkal2l based on sequence similarity to alkal2) | 2: 1,220,413-1,229,520 |
| red hue in body; proportion of red in body and fins | bco1l | beta-carotene oxygenase 1, like | 2: 1,275,344-1,279,650 |
| iridesence | adsl | adenylosuccinate lyase | 8: 2,057,799-2,062,135 |
| red saturation in fins | slc2a15b | solute carrier family 2 member 15b | 1: 15,674,584-15,680,614 |
| black spread in head | kitlga | kit ligand a | 6: 18,236,164-18,262,486 |
| crowning in anal fin | frmd6 | FERM domain containing 6 | 22: 8,439,990-8,458,042 |
| crowning in caudal fin | tfap2b | transcription factor AP-2 beta | 24: 12,480,700-12,489,712 |
| crowning in caudal fin | tfap2d | transcription factor AP-2 delta | 24: 12,494,194-12,506,588 |
| albinism (*15*) | mitfa | melanocyte inducing transcription factor a | 5: 2,408,544-2,413,248 |
| QTL for spotting on dorsal fin (*15*) | Unknown | - | 8: 1,600,000-1,870,000 |
| QTL for red pigmentation on caudal tail (*15)* | Unknown | - | 1: Not available- Not available |
| QTL for red pigmentation on head (*15)* | Unknown | - | 5: Not available- Not available |
| elephant ear on pectoral fin *(15)* | kcnh8 | potassium voltage-gated channel, subfamily H, member 8 | 11: 7,753,376-7,794,461 |
| double tail on caudal fin (*15)* | zic1 | zic family member 1 | 4: 18,982,259-18,985,650 |
| double tail on caudal fin (*15)* | zic4 | zic family member 4 | 4: 18,988,191-18,993,322 |

**Table S3. Genotyping primer sequences.** Primer sequences used for genotyping.

| Type | Chromosome: position ref>alt | Primer Sequences (5'-…-3') | Amplicon Length (bp) |
|---|---|---|---|
| Sequence-based genotyping | 9: 28850248 G>T | *For:* GTCTCGTGGGCTCGGAGATGTGTATAAGAGACAG TAACGCCGTAGCGTTAGCTT | 201 |
| | | *Rev:* TCGTCGGCAGCGTCAGATGTGTATAAGAGACAG CCTCATAATTCACAAAGGTGAAAG | |
| | 9: 28858562 A>G | *For:* GTCTCGTGGGCTCGGAGATGTGTATAAGAGACAG CGATTTTCCTGCCTTAATCC | 299 |
| | | *Rev:* TCGTCGGCAGCGTCAGATGTGTATAAGAGACAG CTTTCCGACGGCAGTGTG | |
| | 9: 28863849 T>C | *For:* TCGTCGGCAGCGTCAGATGTGTATAAGAGACAG AAATCTGAGCCTTTATTTCTACACA | 322 |
| | | *Rev:* GTCTCGTGGGCTCGGAGATGTGTATAAGAGACAG CGGAATCAGGCACAGTAAAA | |
| | 9: 28864161 G>A | *For:* GTCTCGTGGGCTCGGAGATGTGTATAAGAGACAG AGCACCAACAGCTGTCCTTT | 299 |
| | | *Rev:* TCGTCGGCAGCGTCAGATGTGTATAAGAGACAG CTGCAGGTCATGGTGGAGT | |
| | 9: 28867677 G>A | *For:* GTCTCGTGGGCTCGGAGATGTGTATAAGAGACAG CAAATGCAGCCTAATTTTTCA | 262 |
| | | *Rev:* TCGTCGGCAGCGTCAGATGTGTATAAGAGACAG GGCGTGACACCGAGAAAC | |
| RFLP-based genotyping | 9: 288533887 T>C | *For:* ACTCATCATCTTCACAGGAAGCA | 112 |
| | | *Rev:* AAAGTGGGGTTTAGGGCAGG | |
| Allele-specific expression amplicon | 9: 28864161 G>A | *For*: TGCTGCTGGAAACCTCCTAT | 316 |
| | | *Rev*: CTGCAGGTCATGGTGGAGT | |

**Table S4. PCR cycling and digestion conditions.** PCR cycling and digestion conditions used for genotyping.

#### *PCR conditions for genotyping*

**Touchdown PCR Cycling Conditions**

| Temperature °C | Seconds | Cycles |
|---|---|---|
| 94 | 60 | 1x |
| 94 | 15 | 7x |
| 64 (-1°C/cycle) | 20 | |
| 68 | 40 | |
| 94 | 15 | 30x |
| 57 | 20 | |
| 68 | 40 | |
| 68 | 300 | 1x |
| 4 | - | |

**Index PCR Cycling Conditions**

| Temperature °C | Seconds | Cycles |
|---|---|---|
| 94 | 60 | 1x |
| 94 | 10 | |
| 64 | 15 | |
| 68 | 30 | 12x |
| 68 | 300 | 1x |
| 4 | - | |

#### *Restriction Fragment Length Polymorphism Components*

| PCR components | Concentration | Volume (uL) per rxn |
|---|---|---|
| OneTaq Master Mix with Standard Buffer | 1x | 7.5 |
| 10 µM Forward Primer | 0.2 µM | 0.3 |
| 10 µM Reverse Primer | 0.2 µM | 0.3 |
| H2O | - | 5.9 |
| Genomic DNA | <100ng | 1 |
| Total | | 15 |

| MluCI digestion components | Volume (uL) per rxn |
|---|---|
| 15ul PCR reaction | 15 |
| MluCl | 0.3 |
| 10x CutSmart Buffer | 3 |
| H2O | 5 |
| Total | 8.3 |
